# Cellfoundry: a GPU-accelerated, multi-physics ABM framework for cellular microenvironment and organoid-scale studies

**DOI:** 10.64898/2026.04.22.720218

**Authors:** C. Borau, R. Chisholm, P. Richmond

## Abstract

Advanced *in vitro* systems such as organoids and microfluidic organ-on-a-chip platforms enable physiologically richer experimentation, but their complexity creates large parameter spaces and makes it difficult to disentangle the mechanistic roles of transport, mechanics, and extracellular microstructure. Agent-based modelling provides a natural computational counterpart to these systems by representing heterogeneous cells as discrete entities coupled through local rules and environmental fields. However, realistic microenvironment models often remain limited by scalability, simplified extracellular matrix representations, and the practical difficulty of calibrating large numbers of parameters.

Here we present Cellfoundry, a computational framework built on a FLAMEGPU2-based modelling template for simulating complex cellular microenvironments. The framework integrates multiple interacting agent populations, including cells, fibrous networks, and focal adhesions mediating attachment dynamics and traction-force transmission. It combines mechanically resolved cell–cell and cell–matrix interactions with multi-species diffusion fields that propagate biochemical signals through the extracellular environment and regulate processes such as metabolism, migration, and cell-cycle progression. Cellfoundry also supports customizable behaviours across multiple cell types, enabling the study of heterogeneous multicellular systems within a unified computational setting.

To support reproducible model development and calibration, the framework includes a fibre-network generation module, automated performance benchmarking workflows, post-processing and reporting utilities, and an Optuna-based Bayesian optimization pipeline with configurable single- and multi-objective targets. Two showcase examples illustrate these capabilities: a migration assay calibrated against fibroblast motility descriptors and a multi-objective organoid growth scenario reproducing target population composition and expansion dynamics and over time. Together, these examples demonstrate how Cellfoundry can be used to build, calibrate, and extend mechanistically interpretable models of coupled biochemical and mechanical dynamics in advanced *in vitro* systems.

**Highlights:**

- Highly versatile, GPU-accelerated agent-based framework for cellular microenvironments
- Explicit fibrous ECM networks with dynamic remodelling and focal adhesion agents
- Coupled mechanics and multi-species diffusion regulate cell behaviour in a highly customizable environment
- Modular architecture with automated benchmarking and Bayesian parameter optimization

## 1. Introduction

Advanced *in vitro* platforms are becoming increasingly important in mechanobiology, disease modelling, and therapeutic testing because they provide experimentally tractable systems that better approximate tissue organisation than conventional 2D cultures [1, 2]. Organoids are self-organising three-dimensional tissue-like constructs, typically derived from stem or progenitor cells, that recapitulate key aspects of development and disease *in vitro* [3, 4]. Organ-on-a-chip (OoC) platforms use microfluidic devices to culture engineered or natural miniature tissues under controlled conditions, enabling more precise regulation of microenvironmental cues than static cultures [5–7]. The convergence of these approaches into OoC systems is widely viewed as a route to improved physiological realism through perfusion, controllable gradients, and multi-tissue coupling [8–10], although challenges in reproducibility, validation, and standardisation remain. For this reason, such platforms currently complement rather than replace animal studies in most experimental pipelines. As their design becomes more sophisticated, however, so does the space of controllable and latent variables. Outcomes such as growth, morphology, differentiation, or migration can depend on coupled biochemical transport of oxygen, nutrients, and secreted factors, mechanical constraints and deformation, cell– cell interactions, and reciprocal feedback between cells and the extracellular matrix (ECM). This complexity creates a clear need for computational companions that can formalise mechanistic hypotheses, explore parameter regimes that are impractical to test experimentally, and help interpret incomplete measurements by linking micro-scale rules to macro-scale observables.

Agent-based models (ABMs) are well suited to this role because they represent heterogeneous cell populations as discrete entities whose behaviour depends on internal state and local microenvironment. In multicellular biology, ABMs are widely used to encode cell-type-specific rules such as proliferation, death, motility, and phenotype switching while allowing tissue-level organisation to emerge from local interactions [11–14]. For microenvironment applications, they are often hybridised with extracellular physics, such as reaction–diffusion partial differential equations (PDEs) for soluble factors and force-based interactions for mechanics, enabling comparison with time-resolved experimental readouts including growth curves, spatial gradients, and deformation [15, 16].

Despite this conceptual fit, two major barriers remain when ABMs are applied to organoid and microfluidic experiments. First, biologically relevant scenarios can quickly involve large numbers of agents, repeated neighbourhood searches, and coupled physical processes, making computational runtime and availability of local memory limiting factors. Scaling ABMs towards more realistic biomedical settings therefore often requires high-performance computing strategies [17–19]. Second, environmental representation is frequently simplified. In many models, the ECM is treated as an effective continuum stiffness or density field, whereas in many tissues and engineered matrices it is a fibrous microstructure whose topology and remodelling directly influence force transmission, migration pathways, and transport [20]. This has motivated a substantial body of work on generating and analysing representative fibre networks, including approaches that reproduce experimentally observed collagen-network statistics such as connectivity, free-fibre length, and local orientation correlation [21–23].

These challenges are reflected in the current modelling ecosystem. Several widely used multicellular ABM frameworks, including Chaste [24] and PhysiCell [25], provide mature toolchains and substantial user communities, but are predominantly optimised for central processing units (CPUs) using shared-memory and distributed parallelism such as OpenMP and MPI [26]. In contrast, a newer generation of tools is explicitly designed to exploit graphics processing units (GPUs), including CBMOS [27], Gell [28], and FLAMEGPU2 [29]. Recent comparative studies have highlighted the potential of GPU execution to extend appropriate models towards simulations with millions of agents [17, 30]. A related challenge is calibration: even when an ABM captures the relevant mechanisms, its practical value depends on identifying parameter sets that reproduce experimental observations, often across multiple conditions. This becomes especially challenging when simulations are computationally expensive and the resulting objective functions are effectively black-box. Under these conditions, Bayesian optimisation is particularly attractive, as it aims to identify high-performing regions of parameter space using as few costly model evaluations as possible [31, 32]. Although such optimisation workflows are well suited to parallel and distributed execution, their practical efficiency at scale depends not only on the optimisation strategy itself but also on the supporting hardware. Large simulations demand substantial compute and memory resources, while distributed execution introduces communication overheads for which both bandwidth and latency may become important constraints.

In this paper we introduce Cellfoundry, a GPU-accelerated framework for simulating cellular microenvironments at *in vitro* and organoid scales. Building on a previously introduced modelling template for cell microenvironment simulation [30], Cellfoundry extends this approach into a modular platform that supports multiple interacting cell populations, multi-species extracellular diffusion fields, explicit fibrous ECM networks represented as discrete fibre– node agents, and focal adhesions represented as individual agents that mediate attachment dynamics and traction-force transfer between cells and fibres (Supp.Video.1). The framework also includes utilities for fibre-network generation, post-processing and reporting of simulation outputs, systematic benchmarking, and parameter optimisation through configurable single-objective and multi-objective workflows.

The remainder of this manuscript describes the architecture and core model components of Cellfoundry, including two showcases to illustrate the platform capabilities. The Appendix provides detailed mathematical formulations of the model components (Appendix A), as well as implementation aspects, optimisation workflows, and supporting tools.

## 2. Multiphysics framework

Cellfoundry is implemented utilising FLAMEGPU2 [29], which provides GPU-parallel execution, message-based communication between agents, and a layer-based scheduling scheme in which the different biological processes are resolved in a prescribed computational sequence. Model definition, parameterisation, and simulation setup can be specified through a Python front end, while the underlying library is implemented in C++/CUDA. User-defined functions may be written either directly in C++ or in Python, in which case they are transpiled to C++ for compilation and execution on the GPU directly ensuring a high level of performance. This implementation allows explicit agent-level representations of cells, extracellular structures, and soluble species to coexist within a single three-dimensional domain while retaining the scalability required for large simulations. Simulation outputs are stored in `.pickle` files for efficient post-processing and can optionally be exported automatically as VTK files for direct visualisation and analysis in tools such as ParaView (used here to create all Supplementary Videos). Beyond the simulation core, the framework also includes user-facing tools intended to improve accessibility and reproducibility. A configuration interface (config UI) enables rapid generation of templates for customised models, including new agent types, agent functions, and parameter sets (see Fig. Supp.Fig.1 and Supp.Video.2). Complementarily, an editor interface (editor UI) provides a more accessible environment for users who wish to inspect the default model structure and modify key parameters without interacting directly with the source code (Fig. Supp.Fig.2). Together, these interfaces facilitate both rapid prototyping and reuse while preserving the modular structure of the framework (Appendix B).

At the modelling level, Cellfoundry is organised as a reusable environment in which relevant biological processes can be selectively combined depending on the application. Existing agent definitions and functions can be modified, extended, or replaced, and new agents and interaction mechanisms can be incorporated when required by a specific assay or tissue context. The framework includes four main agent classes (Fig. 1, bottom left): 1) **CELL** agents provide a coarse-grained representation of individual cells, centred on the nucleus and associated internal state variables. One of these variables is *cell_type*, which enables multiple phenotypes and behaviours to be encoded within a single agent class; 2) **ECM** agents define the extracellular background domain in which diffusing chemical species are stored and updated; 3) **FNODE** agents represent the nodes of a discrete fibre network, used to capture extracellular matrix architecture and mechanics, that can be manually defined or automatically generated through a dedicated module (Section A.6); 4) **FOCAD** agents represent dynamic focal adhesions that link cells to nearby fibre nodes and mediate bidirectional force transmission and mechanosensitive coupling. In addition, although they are not formal agents, six parallel planes define the simulation boundaries, enabling geometrical confinement and, when required, prescribed boundary motion and mechanical loading along with different boundary conditions (e.g. fixed, elastic or periodic, sticky or slippery etc.). The full list of agent variables and associated functions is provided in Appendix F.

**Figure 1:**
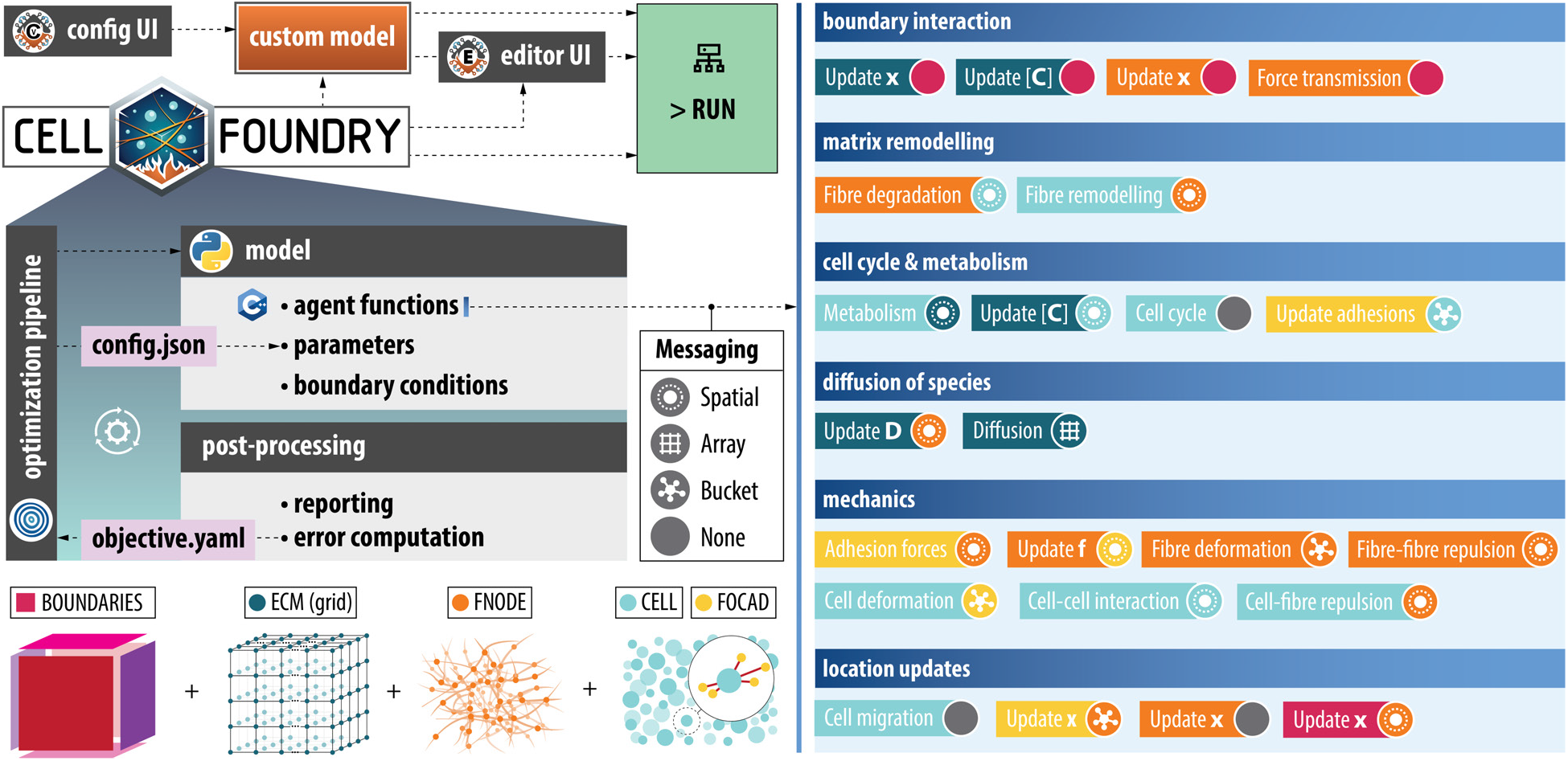
Cellfoundry framework overview. Apart from direct manipulation of the main files, the workflow includes two complementary interfaces: the configuration UI enables rapid generation of templates for fully customized models, including the creation of new agents and agent functions, while the editor UI is designed for less technical users, providing intuitive navigation and convenient adjustment of key parameters within the existing framework. The resulting model is executed on the FLAMEGPU2 platform, leveraging GPU acceleration for large-scale simulations. The architecture is fully modular, allowing complete customization of boundary conditions and all existing agent types (ECM grid, FNODE fibre network, CELL, FOCAD) and their associated functions, as well as the addition of new agents and behaviours. During execution, interactions are coordinated through multiple messaging schemes (spatial, array, and bucket), while the simulation loop is organized into fully customizable functional modules. Right panel shows main agent functions (boxes) and their input messages (circles), colored by calling/broadcasting agent respectively. Post-processing and optimization components support automatic reporting and objective-driven parameter tuning, enabling a flexible and extensible multi-physics simulation framework

Within this architecture, cell behaviour emerges from the coupling of several interacting subsystems, including cell mechanics, focal adhesion dynamics, migration, cell-cycle progression, extracellular diffusion and metabolism (Video Supp.Video.3), and fibre-network mechanics, and remodelling (e.g. matrix degradation [Supp.Video.4], matrix deposition [Supp.Video.5]). These components are described separately in Appendix A. The same modular structure also facilitates integration with calibration and optimisation workflows (Section 3), enabling systematic exploration of high-dimensional parameter spaces and quantitative matching of simulation outputs to experimental data.

### 2.1. Message communication strategies

Communication between agents in FLAMEGPU2 is performed through message lists, where the chosen message type determines how information is indexed and accessed by downstream agent functions. In Cellfoundry, this is important because different interaction patterns require different communication strategies depending on whether agents exchange information through discrete categories, local spatial neighbourhoods, or structured grid locations:

#### Bucket messages

Bucket messages are intended for communication indexed by a discrete integer label. Each emitted message is assigned an integer key within a predefined range, and reader agents retrieve all messages associated with a selected key. This strategy is appropriate when interactions are naturally grouped by category, region identifier, or other discrete partitions, rather than by geometric distance.

#### Spatial messages

Spatial messages are designed for neighbour-based interactions in continuous space. In this case, messages are associated with spatial coordinates and internally organised using a discretisation of the simulation domain into bins. Reader agents query messages around a search origin using a prescribed interaction radius, which makes this strategy well suited, for example, for local cell–cell or cell–matrix interactions. An additional distance check may still be required inside the agent function to reject messages lying outside the true interaction radius.

#### Array messages

Array messages are intended for communication on structured discrete domains. Messages are written to fixed positions in one-, two-, or three-dimensional arrays and are then accessed directly through their corresponding indices. This strategy is especially useful when the communication pattern is naturally aligned with a regular lattice, such as voxelised fields or grid-based environmental data (e.g. diffusion of species).

### 2.2. Core model components

The model is composed of coupled modules describing cell mechanics, focal adhesion turnover and force transmission, cell migration, cell-cycle progression, extracellular species transport, and fibre-network mechanics and remodelling. Note that all modules may be activated or deactivated depending on the specific application, although there may be dependencies between them (e.g. focal adhesion dynamics can not be active if there are no cells or fibre network present). In summary, the model includes: i) Nuclear stress and strain emerging from the traction forces exerted by focal adhesions, which dynamically assemble, attach to the extracellular matrix, detach in a force-dependent manner, and mechanically reinforce under load; ii) Cell motion is governed by the combined action of persistent motility, stochastic fluctuations, chemotactic and chemokinetic responses, durotaxis, and short-range mechanical interactions; iii) Cell proliferation and survival are regulated through a configurable cell-cycle template including growth, stochastic division, damage accumulation, and death pathways; iv) Extracellular chemical fields evolve on an ECM-associated lattice through diffusion, cell-mediated exchange, and intracellular metabolic reactions; v) When required, the extracellular matrix is represented explicitly as a deformable fibrous network with remodelling driven by nearby cells. Detailed mathematical formulations and implementation details are presented in Appendix A.

## 3. Parameter Optimization

Model parameters can be calibrated automatically using Bayesian optimization, implemented with the Optuna framework (v5)[33]. The procedure minimizes one or more user-defined objective (error) functions that quantify the discrepancy between simulation outputs and experimental/target reference data.

### Optimization Framework

Let ***θ*** = (*θ*_1_, …, *θ*_*d*_ ) ∈ Θ denote the vector of *d* tunable model parameters, where Θ is the admissible search space defined by per-parameter bounds. Each parameter may be continuous, integer-valued, or categorical, and may be sampled on a linear or logarithmic scale. The optimization solves:

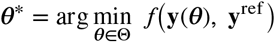

where **y**(***θ*** ) is the simulation output produced with parameter set ***θ*, y**^ref^ is the experimental or target reference data, and *f* is a scalar-valued objective function measuring the error.

For single-objective studies the Tree-structured Parzen Estimator [34] is used as the surrogate model. TPE models the conditional distributions *p*(*θ*_*i*_ ∣ *f* < *f* ^∗^) and *p*(*θ*_*i*_ ∣ *f* ≥ *f* ^∗^) via kernel density estimation and selects the next trial by maximizing the expected improvement ratio *p*(*θ*_*i*_ ∣ *f* < *f* ^∗^) / *p*(*θ*_*i*_ ∣ *f* ≥ *f* ^∗^).

When multiple, potentially conflicting objectives are optimized simultaneously, i.e.

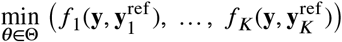

the framework switches to the NSGA-II genetic algorithm [35], which maintains a Pareto front of non-dominated solutions across generations.

### Trial Execution

In the present implementation, Optuna trials are executed sequentially. However, this is a workflow choice rather than a limitation of the framework, as Optuna supports parallel and distributed optimisation through shared study storage, allowing multiple workers to evaluate trials concurrently.

Each optimization trial proceeds as follows:

1. **Parameter suggestion**. The sampler proposes a candidate ***θ***^(*t*)^ based on the history of previous evaluations.
2. **Simulation**. The model is executed as an independent subprocess with the suggested parameters injected as runtime overrides. Simulations are subject to user-defined timeouts and termination criteria, automatically pruning failed or excessively long-running trials.
3. **Result collection**. Simulation outputs are serialized to disk, specifically to a `.pickle` file, at the end of the run.
4. **Error evaluation**. The objective function computes a scalar error from the simulation output and the reference data (see next section and Appendix D for user-defined objectives details).
5. **Database update**. The parameter values and objective value(s) are stored in an SQLite database, enabling study resumption, parallel execution, and post-hoc analysis.

Derived model quantities (e.g. agent’s messages search distances) that depend algebraically on the tuned base parameters are automatically recomputed before each simulation, ensuring internal consistency of the parameter set.

### Objective Functions

The framework provides a library of pre-defined objective functions that can be selected directly from the optimization configuration. New objectives can also be added without modifying the optimization driver, as long as they follow the same input-output interface. The currently implemented objectives span time-series matching, scalar endpoint targets, and specific experiment-oriented targets such as mechanical curve fitting, geometric descriptors (e.g. organoid size), and population-level motility statistics (see Appendix D).

#### Time-series objectives

For outputs recorded as a time series **y** = (*y*_1_, …, *y*_*T*_ ) with a corresponding reference 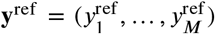, the simulation series is linearly interpolated to match the reference length *M*, and the discrepancy is evaluated through the root-mean-square error

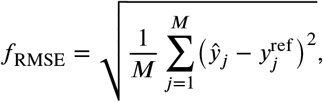

where *ŷ*_*j*_ denotes the interpolated simulation value at the *j*-th reference point. This class of objectives can be used, for example, for cell-population dynamics, focal-adhesion attachment ratios, boundary force histories, and matrix-remodelling metrics such as cumulative degradation, reinforcement, etc.

#### Scalar endpoint objectives

For quantities measured only at the final simulation step, the error is computed as an absolute difference,

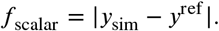

This category includes, for example, the final total alive-cell count or the final average number of focal adhesions per cell. For cell-count objectives, either a single global target or separate targets for individual cell types can be specified. When multiple endpoint targets are provided, the corresponding absolute errors are summed or averaged.

### Implementation Details

The optimization pipeline is implemented in Python and interfaces with the FLAMEGPU2 simulation engine via its `pyflamegpu` Python bindings. All optimization logic resides in a self-contained module that communicates with the simulator through a command-line interface, making the optimization workflow largely independent of the internal simulation implementation.

The full configuration, including search space, objective function, sampler, number of trials, and fixed runtime overrides, is specified declaratively in a single YAML file, facilitating reproducibility and experiment management (see Listing 1 example). This YAML configuration is internally parsed into a JSON file that is passed to the main model via the `–overrides` flag, enabling a clean and consistent replacement of the corresponding simulation parameters without modifying the source code. As stated before, trial results are persisted in an SQLite database, enabling checkpointed long-running studies, resumption of interrupted runs, parallel execution, and interactive inspection through the Optuna Dashboard web interface. Upon completion of the optimization process, an additional JSON file containing the best-performing parameter set is generated, allowing straightforward reproduction of the optimal simulation and further post-processing or analysis if required.

**Listing 1:**
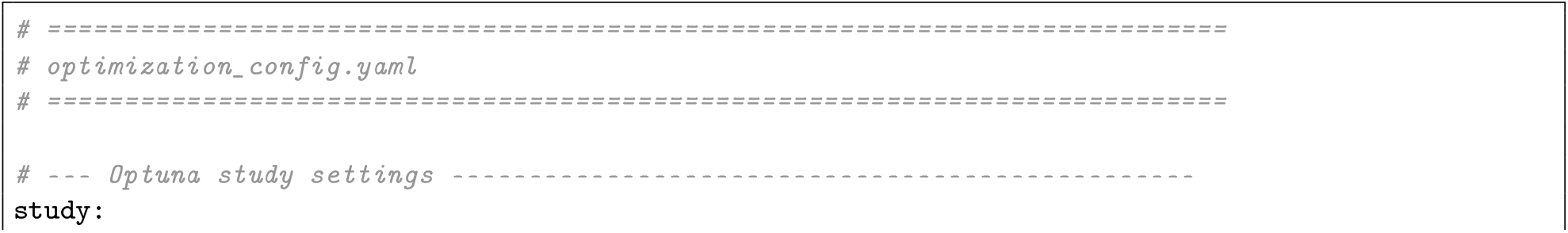
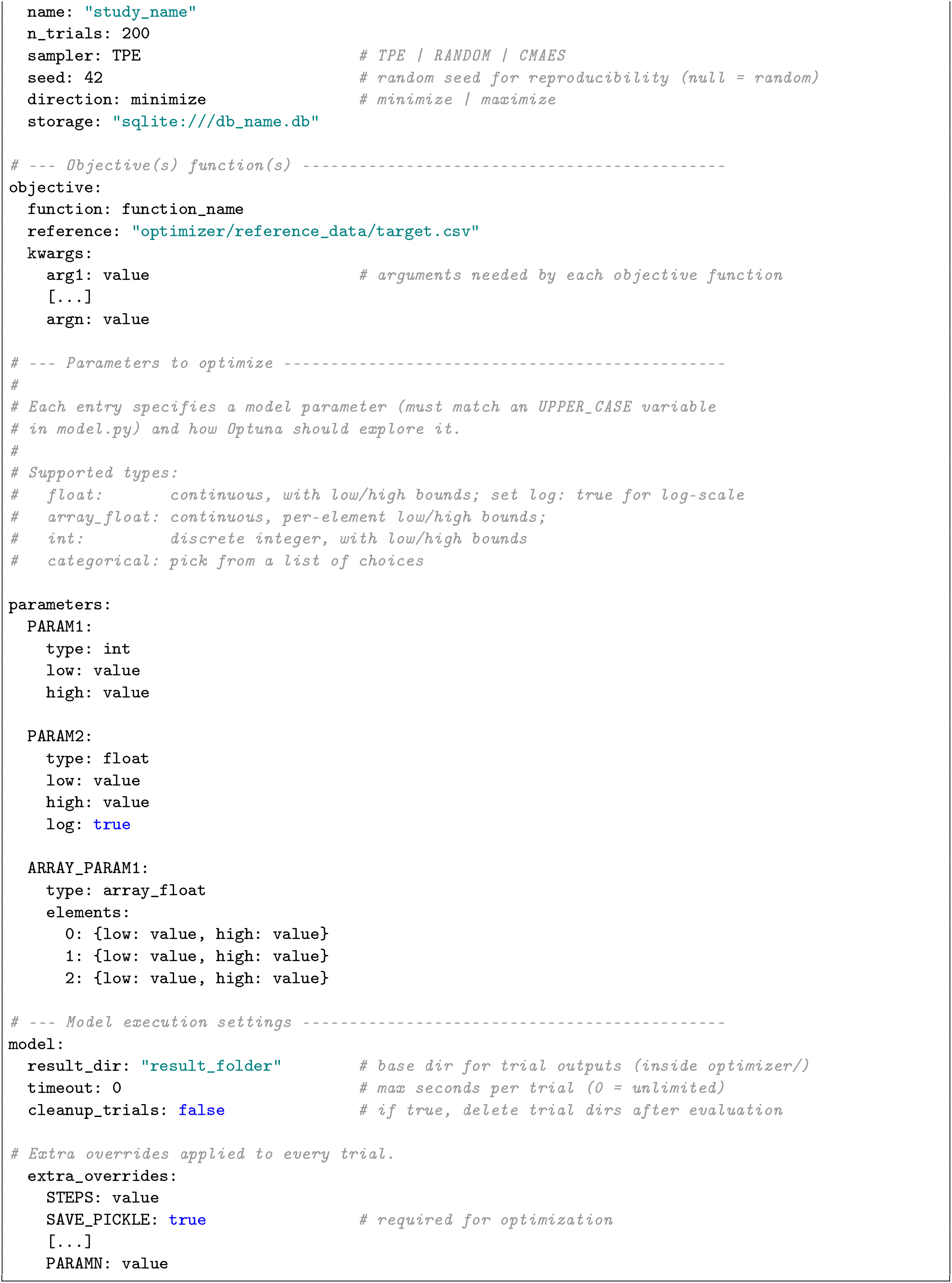
Example YAML configuration file for parameter optimization with Optuna.

## 4. Showcase examples

In this section, two showcase examples are presented to illustrate the capabilities of Cellfoundry. Although not intended to reproduce specific experiments in full detail, these examples demonstrate how the framework can be used to model coupled mechanical and biochemical dynamics, as well as how it can be calibrated against experimental data through the integrated optimization pipeline. The selected cases include: (i) a cell migration assay with chemical inputs, and (ii) an organoid growth scenario. For each case, the model setup, key parameters, and simulation outcomes are described.

### 4.1. Cell migration assay

The first showcase example was designed to capture the key features of a typical cell migration assay. As a reference, we considered the work of [36], where the effects of the growth factor TGF-*β*1 on fibroblast migration in lung adenocarcinoma were investigated using a microfluidic device. In that study, the migration of three cell types (shControl, shSMAD2, and shSMAD3) through a 3D collagen matrix was quantified. Here, the goal was to reproduce the main motility descriptors reported, namely the mean velocity and the effective velocity, across the different experimental conditions (with and without TGF-*β*1).

These metrics were extracted from the published data and used as targets in the optimization pipeline. In particular, the mean velocity (*v*_mean_) corresponds to the average instantaneous speed across all cells and time points, while the effective velocity (*v*_eff_ ) is defined as the ratio between net displacement and total trajectory time, providing a measure of migration directionality. It is worth noting that *v*_eff_ depends on both the total observation time and the temporal resolution, whereas *v*_mean_ is largely independent of these factors. As a consequence, parameters calibrated at a given temporal scale may not directly transfer to simulations with different time step sizes or durations.

Simulations were performed in a 3D cubic domain of 1000 *μ*m per side, with 100 cells migrating for 24 hours using a time step of 1 minute. To remain consistent with the low-density conditions of the reference experiments, cell–cell interactions were disabled, and no fibre network was included. Calibration was carried out in two stages. First, the control condition (absence of TGF-*β*1) was optimized to reproduce the baseline motility of each cell type. Then, the parameters governing baseline speed were fixed, and a second optimization was performed under a uniform TGF-*β*1 concentration (2.5 ng/ml), this time including chemokinesis-related parameters in the search space. In both stages, the optimizer was limited to a maximum of 200 iterations, bounding the computational effort rather than looking for exhaustive convergence

The resulting simulations show good agreement with the experimental migration descriptors indicating successful calibration of the model. Under basal conditions, the shSMAD2 population displays the highest mean velocity and effective velocity, with simulated central values around 1.5 × 10^−3^ *μ*m/s and 6 × 10^−4^ *μ*m/s, respectively (Fig. 2A, panels 1 and 3), together with the highest directionality ratio over time (Fig. 2B, panel 1). This behaviour is consistent with its larger fitted *v*_ref_, combined with lower Brownian noise and rotational diffusion, which favour more persistent trajectories (1). In contrast, shSMAD3 shows the weakest motile phenotype, with lower effective velocity (Fig. 2A, panel 3) and lower directionality ratio (Fig. 2C, panel 1), reflecting a stronger contribution of stochastic reorientation and a reduced persistence .

**Figure 2:**
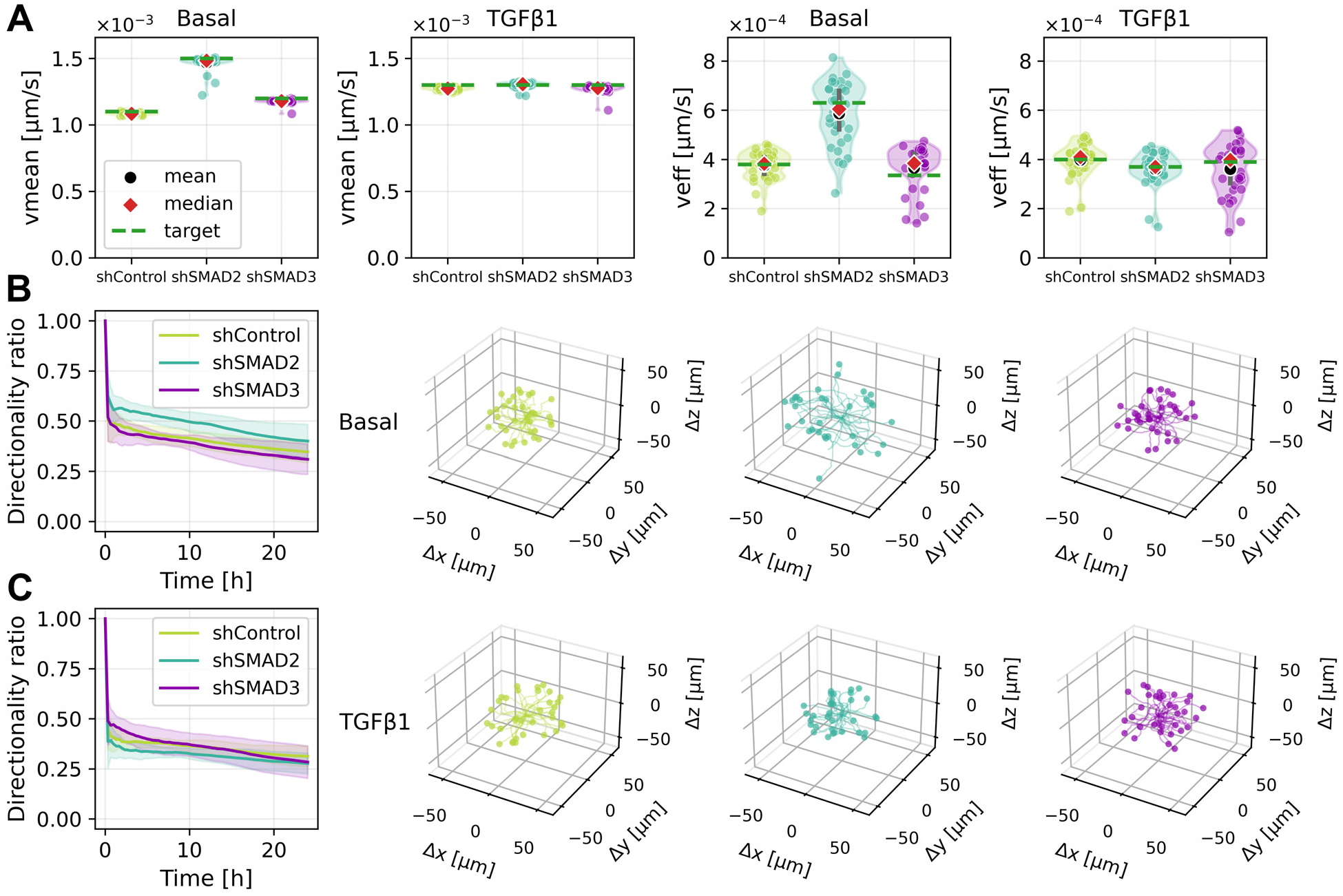
Migration calibration showcase under basal conditions and TGF-*β*1 stimulation. A) Distribution of simulated single-cell mean velocity (*v*_mean_) and effective velocity (*v*_eff_ ) for shControl, shSMAD2, and shSMAD3 populations under basal conditions and in the presence of TGF-*β*1. Black circles indicate the mean, red diamonds the median, and dashed green lines the experimental targets used for calibration. B) Simulated directional persistence under basal conditions. The left panel shows the time evolution of the directionality ratio, computed as net displacement divided by accumulated trajectory length, and the remaining panels show representative 3D cell trajectories over 24 h for each population. C) Simulated directional persistence and trajectories in the presence of TGF-*β*1.

Overall, the mean-velocity distributions are comparatively narrow, whereas the effective velocity exhibits a much larger spread across cells. This difference is expected because mean velocity is computed from the accumulated trajectory length divided by migration time and therefore remains stable even for tortuous paths, while effective velocity is based on net end-to-end displacement divided by time and is thus much more sensitive to directional loss caused here by Brownian motion and rotational diffusion. The representative trajectories in Fig. 2B,C are consistent with this interpretation, showing broader spatial exploration for shSMAD2 and more confined, less persistent motion for shSMAD3. The re-calibration in the presence of TGF-*β*1 further clarifies how these differences arise from distinct chemokinetic responses. Because the adaptation timescale *τ* was chosen to be much longer than the simulation duration, chemokinesis acts here as an approximately sustained modulation of the persistent motility term. In this regime, the sign and magnitude of *α*_ck_ determine whether the persistent component is enhanced or suppressed (1). The optimized parameters reveal a strong inhibitory response for shSMAD2 (*α*_ck_ = −0.761), which markedly reduces effective displacement and directionality relative to basal conditions while leaving the mean velocity much less affected (Fig. 2A, panels 2 and 4). This reproduces the experimentally observed confinement-like behaviour, in which cells remain locally active but lose net migratory efficiency. In contrast, shSMAD3 shows only a weak positive response (*α*_ck_ = 0.167), leading to modest changes relative to basal conditions, while shControl displays an intermediate increase in activity (*α*_ck_ = 0.085). Overall, the simulations capture both the ranking of the cell lines and the distinct effect of TGF-*β*1 on instantaneous motility, net displacement, and directional persistence.

**Table 1.** Calibrated migration parameters for each cell type under control conditions and in the presence of TGF-*β*1 (chemokinesis enabled). Cubical domain: 1000 *μ*m per side. Time step: 1 minute. Simulation duration: 24 hours.

| Condition | Parameter | Units | shControl | shSMAD2 | shSMAD3 |
| --- | --- | --- | --- | --- | --- |
| Control | $v_{\text{ref}}$ | $\mu\text{m/s}$ | $5.07 \times 10^{-4}$ | $8.48 \times 10^{-4}$ | $5.37 \times 10^{-4}$ |
| | $B$ | $\mu\text{m/s}$ | 2.03 | 1.56 | 2.09 |
| | $D_{\text{rot}}$ | $\text{rad}^2/\text{s}$ | $3.5 \times 10^{-5}$ | $3.7 \times 10^{-5}$ | $5.5 \times 10^{-5}$ |
| TGF-β1 | $v_{\text{ref}}$ | $\mu\text{m/s}$ | $5.07 \times 10^{-4}$ | $8.48 \times 10^{-4}$ | $5.37 \times 10^{-4}$ |
| | $B$ | $\mu\text{m/s}$ | 2.44 | 1.52 | 2.26 |
| | $D_{\text{rot}}$ | $\text{rad}^2/\text{s}$ | $3.5 \times 10^{-5}$ | $3.7 \times 10^{-5}$ | $5.5 \times 10^{-5}$ |
| | $\alpha_{\text{ck}}$ | — | 0.085 | -0.761 | 0.167 |

As an additional *in silico* experiment, the calibrated migration model was used without further parameter changes to evaluate a transient, spatially heterogeneous TGF-*β*1 delivery scenario. Instead of imposing a uniform steady concentration of 2.5, TGF-*β*1 was allowed to diffuse from the domain boundaries through a 21 × 21 × 21 ECM lattice, corresponding to voxels of 50 × 50 × 50 *μ*m^3^ (Fig. 3A,B). Two transport regimes were considered. In the first, diffusion was spatially uniform, with a constant *D*_s_ throughout the ECM (see Appendix A.5). In the second, diffusion was hindered by the local fibre-network density, using a spatially varying coefficient of the form

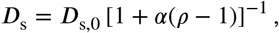

where *α* controls the strength of the reduction and *ρ* is the ratio between the local number of neighbouring fibre nodes and the average number of fibre nodes per voxel. The fibre network was generated with the procedure described in Appendix C; an example of the resulting network and its associated density field is shown in Fig. 3C.

**Figure 3:**
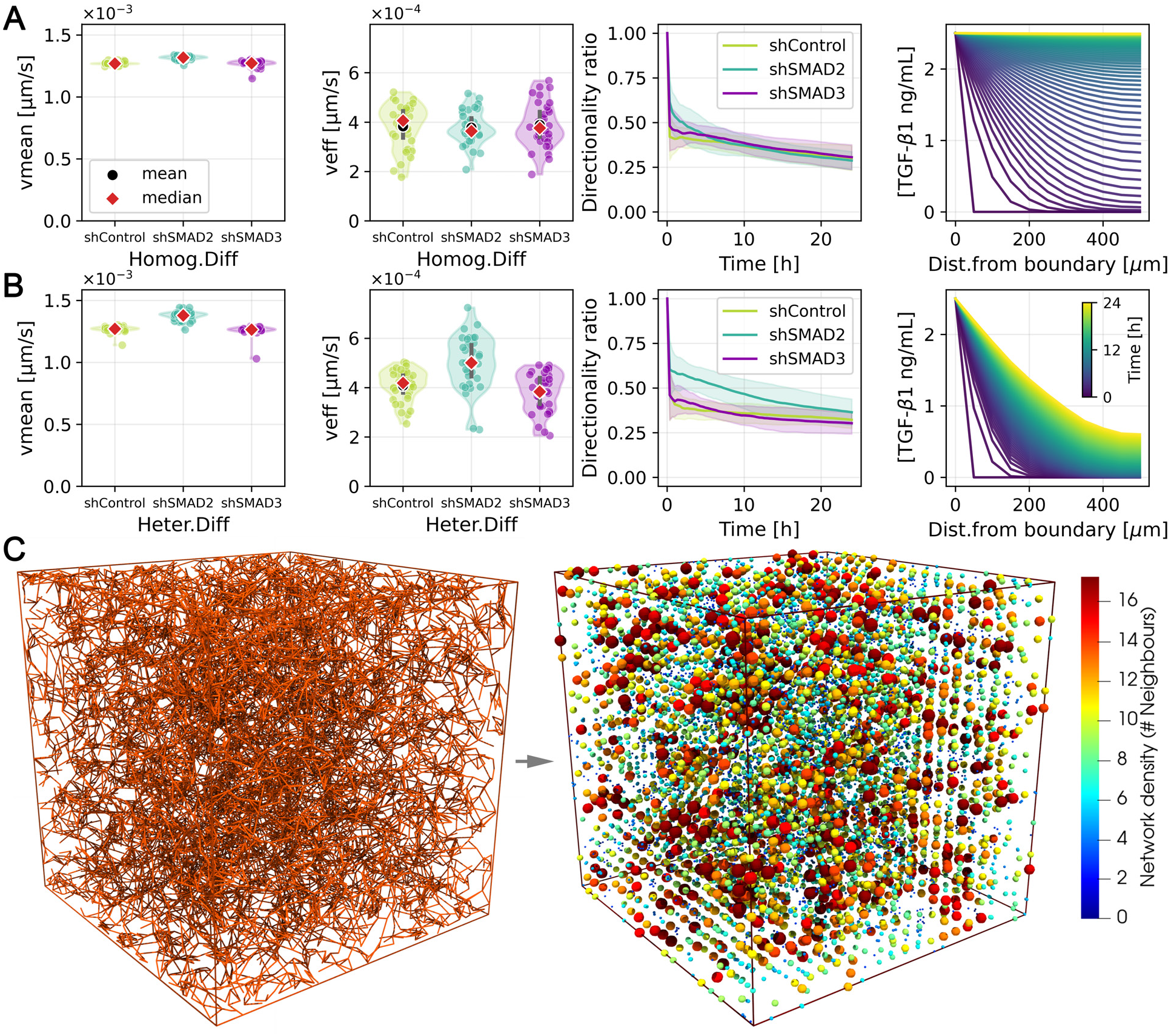
Transient TGF-*β*1 transport scenarios and resulting migration descriptors. A) Homogeneous diffusion case, with spatially uniform *D*_s_. From left to right, the panels show the distributions of mean instantaneous speed (*v*_mean_), effective speed (*v*_eff_ ), directionality ratio over time, and TGF-*β*1 concentration profiles as a function of distance from the domain boundary during the 24 h simulation. B) Heterogeneous diffusion case, in which *D*_s_ is reduced according to the local fibre-network density, shown using the same set of migration and transport descriptors. In the violin plots, black circles indicate means and red diamonds indicate medians. In the concentration plots, colour encodes time. C) Example of the generated fibre network (left) and the corresponding voxel-scale network-density field (right), which is used to define the spatially varying diffusivity in the hindered-diffusion scenario. Each point is colored and scaled according to the number of neighbouring fibre nodes within the corresponding voxel for better visualization.

Under free diffusion, TGF-*β*1 nearly saturated the ECM within the simulated 24 h (Fig. 3A, rightmost panel), and the resulting migration behaviour remained close to that obtained under the previously calibrated uniform-concentration condition, as reflected by similar distributions of migration descriptors and only moderate changes in directionality ratio over time (Fig. 3A, first three panels), although with a broader distribution of effective velocities due to the transient establishment of the field. In contrast, hindered diffusion preserved stronger spatial heterogeneity over time (Fig. 3B, rightmost panel), shifting migration descriptors towards the basal case (Fig. 3B, first three panels). This shift was only partial, however, because cells were still exposed to non-zero TGF-*β*1 levels and persistent gradients throughout the simulation. Together, these results illustrate that, even with identical motility parameters, the spatial and temporal structure of growth-factor transport can substantially alter the apparent migratory phenotype.

More generally, this example highlights the value of the framework as a predictive tool for probing how transport, matrix architecture, and cell motility jointly shape assay-level behaviour. Because the calibration performed here isolates the roles of baseline persistence, stochastic reorientation, and chemokinetic modulation, subsequent simulations can attribute behavioural changes either to altered transport conditions or to changes in the underlying cell parameters. In turn, the same staged calibration strategy can be extended to progressively incorporate additional mechanisms such as proliferation, apoptosis, matrix remodelling, or cell–cell interactions, enabling controlled model expansion while preserving interpretability.

### 4.2. Organoid growth scenario

The second showcase example focuses on organoid growth dynamics. This case was inspired by the work of [37], where long-term live imaging of murine pancreas-derived organoids revealed substantial heterogeneity in growth trajectories, even among organoids with similar initial cell numbers. Their analysis highlighted how differences in proliferation dynamics can lead to markedly distinct morphological and volumetric outcomes over time.

In the present example, the objective was to reproduce a realistic organoid growth scenario over a period of five days. A representative growth curve was extracted from their results and used as a target. This curve was expressed in terms of the radius of gyration (*R*_*g*_) and its equivalent sphere radius (*R*_eq_), allowing direct comparison with simulation outputs (see Appendix D for definitions). In addition to matching the global growth trend, a compact morphology was enforced by minimizing the mean nearest-neighbour cell distance towards a prescribed value corresponding to the cell diameter (Supp.Video.6). Finally, an arbitrary heterogeneous population structure (not present in the original study) was imposed by defining three cell types with distinct final quantities, chosen to illustrate different proliferation dynamics (target populations: 1000, 400, and 200 cells for types 0, 1, and 2, respectively). These targets were intentionally selected to represent a solid organoid, in contrast to the lumen-containing structures reported in the reference study, which required approximately double the experimental cell count. The optimization problem was therefore formulated with three objectives: (i) minimizing the error with respect to the target growth curve, (ii) minimizing the deviation of the mean cell–cell distance from the desired compact configuration, and (iii) minimizing the discrepancy in both total and per-type cell populations at the end of the simulation. The simulation was performed in a cubic domain of 1000 *μ*m per side, with a random initial population *N*_cells_ (treated as a calibrated parameter) placed in a compact cluster near the domain centre. Due to the higher complexity compared to the first scenario, a total of 500 simulations were allowed during the optimization process. The temporal discretization was fixed to 2400 steps with a time step Δ*t* = 180 s, corresponding to a total simulated time of 120 hours (5 days). A subset of parameters was fixed to reduce the dimensionality of the search space and enforce biologically consistent behaviour. In particular, the durations of the S, G2, and M phases (*T*_S_ = 8 h, *T*_G2_ = 4 h, *T*_M_ = 2 h) were kept constant across all cell types. In contrast, key parameters controlling motility, cell–cell mechanical interactions, and proliferation were calibrated. These included the reference migration speed *v*_ref_, the rotational diffusion coefficient *D*_rot_, the cell–cell adhesion and repulsion stiffnesses (*K*_adh_ and *K*_rep_), and the G1 phase duration *T*_G1_ for each cell type.

Table 2 summarizes the calibrated parameters. The optimized values reveal a regime dominated by strong repulsion (*K*_rep_) relative to adhesion (*K*_adh_), which, in combination with proliferation dynamics, ensures that cells remain closely packed while avoiding excessive overlap, thereby satisfying the compactness constraint. In fact, the calibrated migration parameters (*v*_ref_ and *D*_rot_ ) indicate that cell motility plays a secondary role compared to proliferation-driven expansion, with structural rearrangements largely induced by growth and local crowding. In practice, motility remains subdominant as long as it does not exceed a threshold at which individual cells can escape the aggregate and compromise organoid cohesion. This regime is effectively controlled by the additional constraint enforcing mean cell–cell distances close to the cell diameter, which prevents dispersion and maintains a compact structure. As could be expected, since it is the only proliferation-related free parameter in the optimization, the duration of the G1 phase (*T*_G1_) dominates the fate of the organoid and the resulting per cell-type populations, with shorter *T*_G1_ leading to faster expansion and ultimately larger population fractions.

**Table 2.**
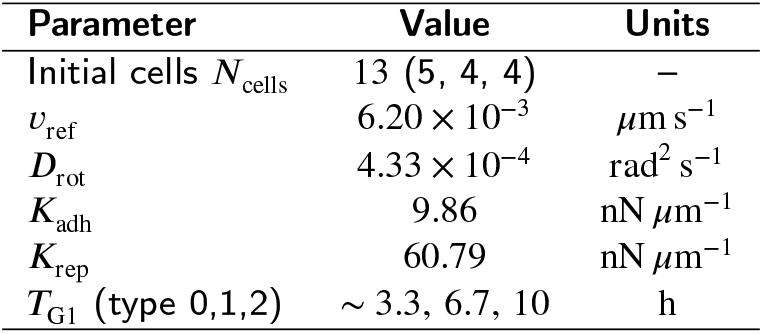
Calibrated parameters for the organoid growth scenario.

The simulated growth dynamics capture the main trend of the target trajectory (Fig. 4A), with the equivalent radius reaching approximately 420 *μ*m at 120 h. Although the agreement is not exact at all intermediate times, the simulation reproduces the overall expansion profile and final size reasonably well, given the complexity of the system and the limited number of parameters available for tuning. The initial lag phase is slightly underestimated, while the later growth phase is somewhat overestimated, which may be attributed to the simplified proliferation model and the absence of additional regulatory mechanisms such as nutrient limitations or mechanical feedback. The sphericity (Fig. 4B) remains relatively high overall (Ψ ≈ 0.6–0.8 at late times), although it exhibits noticeable fluctuations, particularly at early and intermediate stages. This noisy behaviour arises from the definition of Ψ based on the ratio of extreme eigenvalues of the covariance matrix (see Appendix D), which makes it sensitive to transient anisotropies, local rearrangements, and stochastic proliferation events. In particular, small clusters and uneven growth fronts at early times can temporarily distort the shape, leading to rapid variations in the metric despite the organoid remaining globally compact during growth (reaching a maximum span of ∼ 1300 *μ*m), while maintaining a controlled spatial distribution of cells (Fig. 4C). The population dynamics (Fig. 4D) show the emergence of the prescribed heterogeneous composition, with final counts approaching almost exactly the target values of ∼ 1000, 400, and 200 cells for types 0, 1, and 2, respectively. These differences arise directly from the calibrated G1 phase durations, which set distinct proliferation rates and drive the divergence in population sizes over time. This behaviour illustrates how the framework links single-cell cycle parameters with emergent, population-level outcomes while simultaneously satisfying global geometric constraints.

**Figure 4:**
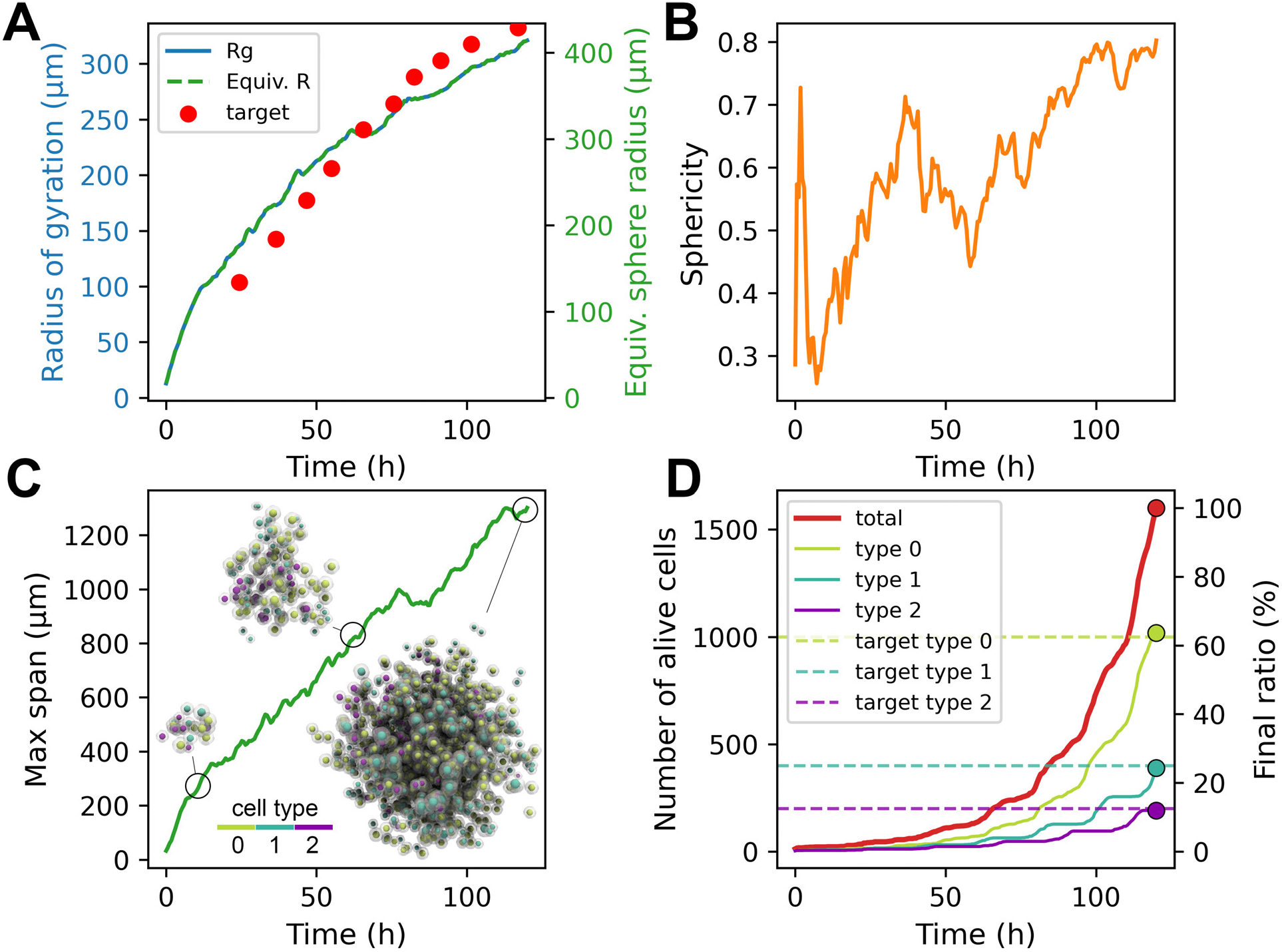
Organoid growth simulation. A) Evolution of the radius of gyration (left y-axis) and equivalent sphere radius (right y-axis) compared to the target growth curve. B) Sphericity metric over time. C) Maximum spatial extent of the organoid, showing monotonic expansion. D) Total and per-type cell populations over time, along with target final proportions.

More generally, this example illustrates how multi-objective optimization can be used to constrain both structural and dynamical properties of organoid systems. As in previous showcases, the approach can be extended to incorporate additional experimental observables, such as spatial patterning, mechanical stresses, or nutrient gradients. Furthermore, once a baseline growth model is established, additional processes such as cell death, differentiation, or extracellular matrix interactions can be introduced and calibrated incrementally, enabling systematic exploration of increasingly complex biological scenarios.

## 5. Benchmarking

### Benchmark protocol

Benchmark simulations were performed to characterize the computational scaling of Cellfoundry across a representative range of model sizes and GPU architectures. A fixed cubic domain of 1*mm*^3^ was used for all simulations with a varying number of agents (up to ∼ 5 × 10^7^). The extracellular matrix was discretized as a cubic lattice of ECM agents with total population

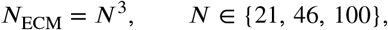

corresponding approximately to 10^4^, 10^5^, 10^6^ ECM agents.

The number of simulated cells was varied over four orders of magnitude,

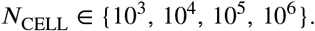

Each CELL agent initialized a fixed number of focal-adhesion agents,

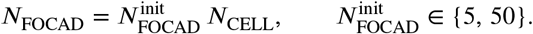

Cell size was varied through

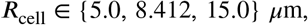

which directly affects interaction range and neighbourhood occupancy.

The fibre scaffold was represented using three pre-generated networks (Appendix C) of increasing density, with

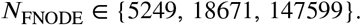

The full parameter sweep comprised up to 216 configurations. Each simulation was run for 100 steps, and two runtime metrics were recorded separately: initialization time and mean runtime per simulation step. Distinguishing these two costs is important because they capture different aspects of the workflow. Initialization time includes model construction, host-side data preparation (raw Python), and GPU memory allocation and transfer, and is therefore influenced not only by the simulation size but also by CPU performance and runtime overheads. In contrast, the per-step runtime reflects the cost of the GPU-executed simulation loop. This distinction is particularly relevant for workloads involving many short simulations, such as parameter sweeps, sensitivity analyses, or model calibration.

### Hardware platforms

Benchmarks were executed on four NVIDIA GPU architectures:

- GTX 1050Ti (desktop GPU, 4GB GDDR5)
- RTX 5000 (workstation GPU, 32GB GDDR6)
- A100 (data-center GPU, 80GB HBM2)
- H100 NVL (data-center GPU, 94GB HBM3)

Due to memory limitations, GTX 1050Ti and RTX 5000 runs were restricted to 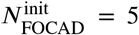, whereas A100 and H100 NVL enabled both 5 and 50 focal adhesions per cell, reaching simulations with more than 5 × 10^7^ agents. Supp.Fig.5 shows example snapshots of the largest configurations used during benchmarking, illustrating the extreme density of agents and interactions that can be simulated on the data-center platforms.

### Scaling behaviour across platforms

Figure 5 summarizes initialization time and mean runtime per step as a function of total agent count. In all cases, runtime increases with problem size, but the spread observed at similar total agent counts shows that total population alone is not sufficient to explain performance. The composition of the system, namely how agents are distributed among ECM, CELL, FOCAD, and FNODE populations, also has a clear impact on runtime due to the different communication and interaction patterns generated by each agent type.

**Figure 5:**
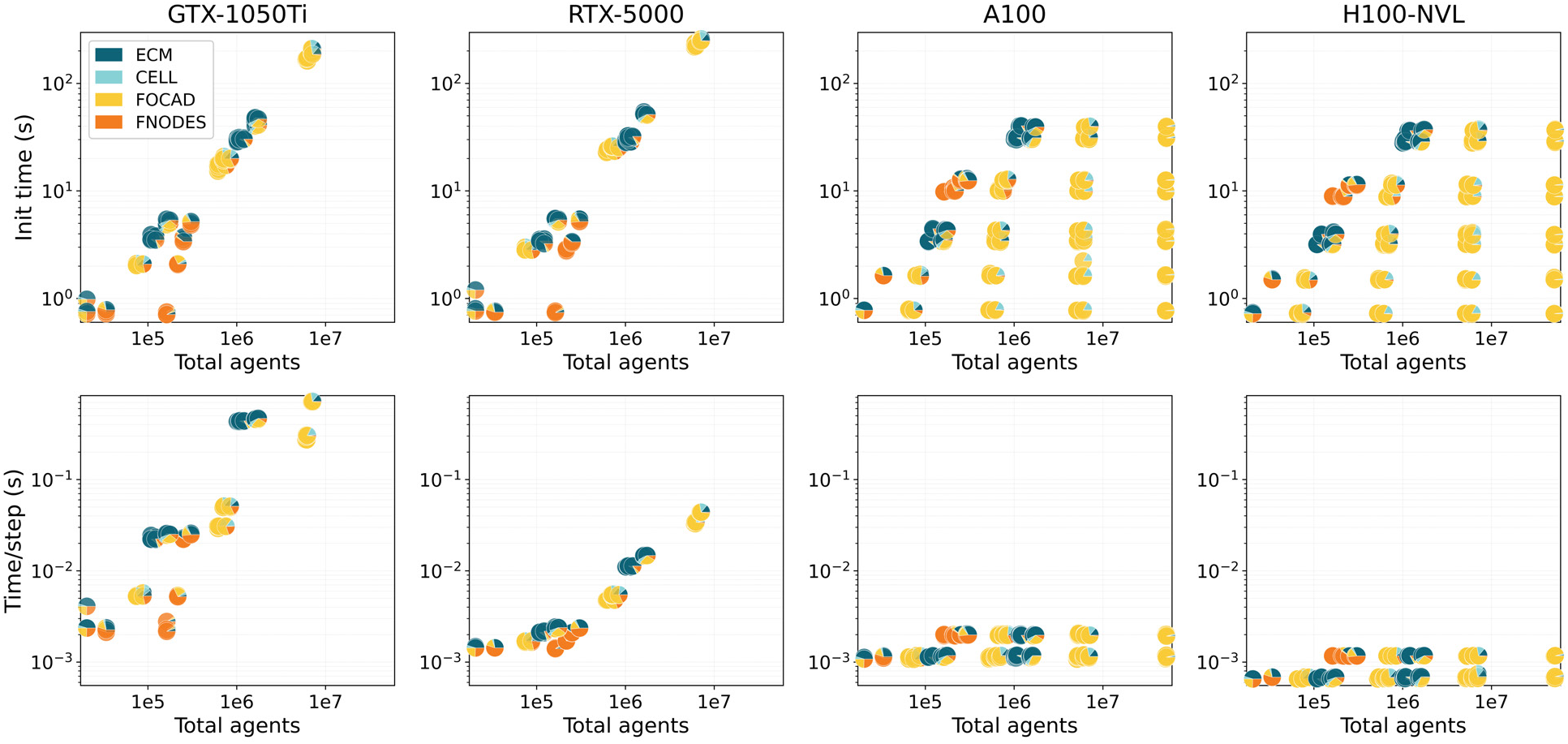
Benchmark runtime (s) across GPU architectures as a function of total agent count, defined as the sum of ECM, CELL, FOCAD, and FNODE agents. Top row: initialization time. Bottom row: mean runtime (s) per simulation step (averaged over 100 steps). Pie charts indicate the proportion of each agent type. Both GTX-1050Ti and RTX-5000 show an exponential growth of both initialization and step times with agent count. On the other hand, both A100 and H100-NVL present a vertical spread at similar total agent counts showing that runtime is influenced not only by the total number of agents, but also by how those agents are distributed across interacting populations.

Initialization time spans a much wider range than per-step runtime. On the GTX 1050Ti and RTX 5000, initialization increases from approximately 10^0^ to 2 × 10^2^ s across the tested configurations, while step times range from milliseconds to tenths of a second (Table 3). These platforms remain suitable for small and medium-scale studies, but become restrictive when either very fine temporal resolution or many repeated runs are required.

**Table 3.** Summary of tested simulation scale, hardware characteristics, and computational cost across GPU architectures. Reported times correspond to the minimum and maximum values observed across the benchmark configurations.

| GPU | Memory | Min-Max agents | Min-Max init time (s) | Min-Max step time (s) |
| --- | --- | --- | --- | --- |
| GTX 1050Ti | 4GB GDDR5 | $\sim 2 \times 10^4 - 7 \times 10^6$ | $\sim 1 \times 10^0 - 2 \times 10^2$ | $\sim 2 \times 10^{-3} - 7 \times 10^{-1}$ |
| RTX 5000 | 32GB GDDR6 | $\sim 2 \times 10^4 - 7 \times 10^6$ | $\sim 1 \times 10^0 - 2 \times 10^2$ | $\sim 1 \times 10^{-3} - 4 \times 10^{-2}$ |
| A100 | 80GB HBM2 | $\sim 2 \times 10^4 - 5 \times 10^7$ | $\sim 8 \times 10^{-1} - 4 \times 10^1$ | $\sim 1 \times 10^{-3} - 2 \times 10^{-3}$ |
| H100 NVL | 94GB HBM3 | $\sim 2 \times 10^4 - 5 \times 10^7$ | $\sim 8 \times 10^{-1} - 4 \times 10^1$ | $\sim 6 \times 10^{-4} - 1 \times 10^{-3}$ |

In contrast, the A100 and H100 NVL sustain substantially larger simulations while maintaining very low stepping cost. Across the explored range, both data-center GPUs support configurations exceeding 5 × 10^7^ agents, with step times remaining in the low-millisecond or sub-millisecond regime. Initialization time still increases with scale, but remains below a few tens of seconds even for the largest cases. The H100 NVL consistently outperforms the A100, especially during the stepping phase, and shows the lowest absolute runtimes at the largest scales.

To quantify these trends, multivariate log–log regression models were fitted separately for initialization time and mean runtime per step, using the benchmark parameters as predictors. The detailed fits and pairwise contour maps are provided in Appendix E (Supp.Fig.4 and Supp.Fig.3). In brief, the desktop and workstation GPUs are mainly influenced by the combined growth of CELL and ECM populations, whereas on the A100 and H100 NVL the remaining variation is more strongly associated with ECM and fibre-network size. For the data-center GPUs, this apparent dominance should be interpreted with caution, because absolute step times remain extremely small across the full sweep.

Overall, the results show that Cellfoundry can span a wide performance envelope, from desktop-scale exploratory simulations to very large data-center workloads. At moderate scales, cost is largely associated with CELL and ECM populations. The number of ECM only plays a role when diffusion of species is activated, so any other problem will be mostly limited by the number of CELL agents. At the largest scales, fibre-network resolution and interaction structure become increasingly relevant, reflecting the higher communication and neighbourhood-query load generated by scaffold-rich and adhesion-rich configurations. This behaviour is consistent with the model design and highlights the importance of considering both population size and interaction topology when estimating simulation cost.

## 6. Code Availability

The Cellfoundry framework is openly available and maintained through public GitHub repositories. The main simulation code, together with the editor interface for model exploration and parameter tuning, can be accessed at: https://github.com/cborau/cellfoundry. The configuration interface (configurator), designed to generate templates for customized models, agents, and functions, is provided in a separate repository at: https://github.com/cborau/flamegpu2_uiconfig.

Both repositories include the necessary source code, example configurations, and documentation to reproduce the results presented in this work and to facilitate extension of the framework to new applications.

## 7. Discussion

Cellfoundry was developed as a high-performance and extensible framework for simulating cellular microenvironments in which mechanics, transport, and extracellular structure are tightly coupled. The results presented here show that this combination can be achieved while maintaining realistic computational costs at scales that are typically challenging for detailed agent-based models. In particular, the benchmarking results demonstrate that large three-dimensional systems (tens of millions of agents) can be simulated efficiently on modern GPU hardware, making tasks such as parameter sweeps and optimization practically feasible. A key feature of the framework is the explicit representation of extracellular structure and force transmission. By modelling fibre networks and focal adhesions as dedicated agent populations, Cellfoundry enables direct investigation of how matrix architecture, connectivity, and mechanical properties influence cell behaviour. This contrasts with approaches in which the extracellular environment is treated in a more implicit or continuum manner, and is particularly relevant for applications where mechanotransduction, force propagation, and matrix remodelling are central to the system dynamics. In this context, the framework provides a natural way to couple local interactions at the cell–matrix interface with emergent, system-level mechanical behaviour. Another important aspect is the integration of parameter calibration within the modelling workflow. The Optuna-based pipeline allows systematic exploration of high-dimensional parameter spaces, including multi-objective formulations in which different experimental observables are matched simultaneously. Importantly, the calibrated parameters do not act as black-box fitting coefficients alone, but retain direct mechanistic meaning within the model. Embedding this capability directly into the framework simplifies reproducibility, reduces manual intervention, and enables a more quantitative connection between simulations and experimental data, especially in complex *in vitro* systems where multiple outputs must be considered jointly. This tight coupling between simulation and calibration is particularly valuable for progressively refining models as additional experimental constraints become available.

Cellfoundry is not intended to compete with existing platforms such as PhysiCell, which provide mature ecosystems, extensive biological models, and broad community support. Instead, it is designed to complement them by targeting a different regime: large-scale GPU-accelerated simulations with an emphasis on mechanical interactions, explicit extracellular structure, and integrated calibration workflows. In this sense, the framework extends the range of problems that can be addressed within agent-based modelling, rather than replacing such established tools. The current implementation also has several limitations. First, the reliance on FLAMEGPU2 implies a dependency on NVIDIA hardware and the CUDA ecosystem, which restricts portability compared to CPU-based frameworks. Nevertheless, ongoing development efforts within FLAMEGPU2 are extending support for AMD hardware. Second, although Python is used for orchestration and optimization, advanced agent function customization still requires interaction with C++ components and GPU-oriented concepts (e.g. atomic operations), which may increase the entry barrier for some users. Third, the framework is still relatively young, lacking in biological scope and the extensive community, model libraries, and validation cases available in more established platforms. Although the framework already integrates many biological and mechanical mechanisms, the present work should be interpreted as a methodological foundation rather than a fully comprehensive biological platform. In fact, the showcase cases were deliberately chosen to demonstrate capability, not to provide exhaustive reconstructions of particular experiments. Thus, the current results validate the flexibility of the framework and its calibration workflow, but they should not be over-interpreted as comprehensive biological validation of all included processes.

These limitations naturally suggest several future directions. On the modelling side, future work should focus on application-driven studies that leverage the full multiphysics structure of the framework in more biologically realistic settings, particularly in the context of microfluidic research. For example, simplified scenarios such as the migration showcase could be extended to explore the effects of different growth factors, interactions between heterogeneous cell populations (e.g. healthy and tumoral), quantify how matrix density and architecture influence cell motility, or assess how inhibiting or enhancing matrix remodelling alters migration patterns. Likewise, organoid applications could be expanded toward lumen formation, matrix remodelling, necrotic core emergence, or mechanically constrained growth in microfluidic devices. Because the framework already combines discrete cells, extracellular mechanics, and soluble factors, these extensions are conceptually straightforward, although they will require careful calibration and validation against richer datasets.

On the computational side, future work should continue to improve usability, reproducibility, and model sharing. Continuous refinement of the configuration UI, more standardized example projects, expanded documentation, richer post-processing templates, and broader libraries of reusable objective functions would make the framework more accessible and easier to adopt. A growing collection of reference models would also help establish modelling conventions and facilitate more systematic cross-study comparisons. Additionally, in the present optimization pipeline, trials are executed sequentially. However, Optuna is compatible with parallel and distributed execution, suggesting a future extension based on FLAMEGPU ensemble functionality or multi-GPU workflows. This could further reduce the computational cost of large-scale calibration and enable broader exploration of complex parameter spaces. In summary, Cellfoundry provides a flexible and scalable framework for studying complex cellular microenvironments enabling new classes of large-scale and mechanistically detailed simulations that are difficult to address with current tools.

## CRediT authorship contribution statement

**C. Borau:** Conceptualization, Methodology, Software, Writing - original draft, Writing - review and editing. **R. Chisholm:** Software, Methodology, Writing - review and editing. **P. Richmond:** Software, Methodology, Writing - review and editing.

