## Appendix for "Cellfoundry: a GPU-accelerated, multi-physics ABM framework for cellular microenvironment and organoid-scale studies"

### Appendix Contents


- **Appendix A:** Detailed Model Formulation
- **Appendix B:** Configurator and Editor UI
- **Appendix C:** Fibre Network Generation Implementation
- **Appendix D:** Pre-defined Optimization Targets
- **Appendix E:** Benchmark Data
- **Appendix F:** Agent API Summary
- **Appendix G:** Agents' C++ Functions Summary
- **Appendix H:** Video Captions

### A. Detailed Model Formulation

#### A.1. Cell mechanics

Each CELL agent represents a cell of radius $R_{cell}$ containing a deformable nucleus with centre position $\mathbf{x}_c = (x, y, z)^\top$ and undeformed radius $R_{nuc}$. The mechanical state of each agent evolves over time as a result of interactions with other agents and the extracellular environment.

Nuclear mechanics are explicitly represented, while the mechanical role of the cytoskeleton is captured implicitly through a set of focal adhesion agents (described in Section A.2). Each FOCAD belongs to exactly one CELL and applies a traction force $\mathbf{f}_i$ at an anchor position $\mathbf{x}_i$ located on the nucleus surface. The number of anchor points is user-defined and remains fixed throughout the simulation (increasing this number provides a higher spatial resolution). Communication between FOCAD and CELL agents is implemented via bucket messages keyed by the CELL identifier, enabling each CELL to efficiently aggregate the mechanical contributions from all of its associated adhesions.

##### *Stresslet-Based Coarse-Grained Stress*

For each adhesion $i$, the lever arm from the nucleus centre to the adhesion site is $\mathbf{r}_i = \mathbf{x}_i - \mathbf{x}_c$. A symmetric stresslet tensor is accumulated over all adhesions:

$$\mathbf{S} = \sum_i \mathrm{sym}(\mathbf{r}_i \otimes \mathbf{f}_i), \qquad \mathrm{sym}(\mathbf{A}) = \tfrac{1}{2}(\mathbf{A} + \mathbf{A}^\top).$$

The average Cauchy stress tensor in the nucleus is obtained by dividing by the nuclear volume $V = \frac{4}{3}\pi R_{nuc}^3$:

$$\boldsymbol{\sigma} = \frac{\mathbf{S}}{V}.$$

##### *Linear Elastic Compliance & Near-Incompressibility Projection*

The elastic strain $\boldsymbol{\varepsilon}^{\mathrm{el}}$ is obtained from $\boldsymbol{\sigma}$ by assuming isotropic linear elasticity. The shear modulus and Lamé parameter are, respectively,

$$G = \frac{E}{2(1+\nu)}, \qquad \lambda = \frac{E\nu}{(1+\nu)(1-2\nu)}.$$

Inverting the isotropic Hooke relation $\boldsymbol{\sigma} = 2G\boldsymbol{\varepsilon} + \lambda\,\mathrm{tr}(\boldsymbol{\varepsilon})\mathbf{I}$ gives

$$\boldsymbol{\varepsilon}^{\mathrm{el}} = \frac{1}{2G}\boldsymbol{\sigma} - \alpha\,\mathrm{tr}(\boldsymbol{\sigma})\mathbf{I}, \qquad \alpha = \frac{\lambda}{2G(3\lambda + 2G)}.$$

To enforce near-incompressibility at the strain level, the volumetric part is removed from the elastic strain:

$$\tilde{\boldsymbol{\varepsilon}} = \boldsymbol{\varepsilon}^{\mathrm{el}} - \frac{\mathrm{tr}(\boldsymbol{\varepsilon}^{\mathrm{el}})}{3}\mathbf{I}.$$

##### *Viscoelastic Relaxation*

The nucleus stores a viscoelastic strain state $\boldsymbol{\varepsilon}$, which relaxes towards the deviatoric elastic strain $\tilde{\boldsymbol{\varepsilon}}$ via first-order linear kinetics with time constant $\tau$:

$$\frac{d\boldsymbol{\varepsilon}}{dt} = \frac{1}{\tau}\left(\tilde{\boldsymbol{\varepsilon}} - \boldsymbol{\varepsilon}\right).$$

This is integrated with an explicit Euler step of size $\Delta t$:

$$\boldsymbol{\varepsilon}^{t+\Delta t} = \boldsymbol{\varepsilon}^t + \frac{\Delta t}{\tau}\left(\tilde{\boldsymbol{\varepsilon}}^t - \boldsymbol{\varepsilon}^t\right).$$

To preserve the validity of the small-strain assumption and prevent numerical runaway from force–geometry feedback, each component of $\boldsymbol{\varepsilon}$ is bounded by a user-defined clamp: $|\varepsilon_{ij}| \le \varepsilon_{\max}$,

***Update of Nucleus Surface Anchors***

Each CELL stores $N$ reference unit vectors $\hat{\mathbf{u}}_a$ ($a = 1, \ldots, N$) defining anchor directions on the unit sphere. The current anchor positions are updated via the small-strain mapping:

$$\mathbf{x}_a = \mathbf{x}_c + R\,(\mathbf{I} + \boldsymbol{\varepsilon})\,\hat{\mathbf{u}}_a,$$

which produces an ellipsoidal deformation consistent with the current strain tensor. The resulting redistribution of anchor positions alters the spatial arrangement of focal adhesions, thereby modulating traction force transmission and influencing the emergence of cell polarity.

#### A.2. Focal adhesions

FOCAD agents are modeled as discrete, stochastic, force-transmitting links between extracellular matrix fibre nodes (FNODEs, described in Section **??**) and the aforementioned prescribed anchor points on the nuclear surface. Each adhesion belongs to a given cell and connects a fibre node at position $\mathbf{x}_{\mathrm{FNODE}}$ to a nucleus anchor point $\mathbf{x}_i$. For an attached adhesion, the instantaneous end-to-end distance is $\ell = \|\mathbf{x}_{\mathrm{FNODE}} - \mathbf{x}_i\|$. All transmitted forces are tension-only and act along $\mathbf{u}$, the unit vector along the adhesion line of action:

$$\mathbf{u} = \frac{\mathbf{x}_{\mathrm{FNODE}} - \mathbf{x}_i}{\ell}, \qquad \ell > 0.$$

The traction force applied to the fibre node is therefore $\mathbf{F}_{\mathrm{FNODE}} = -F\mathbf{u}$, pulling the fibre node toward the cell. Force computation if explained in Section A.2-Mechanics.

***Turnover dynamics***

Focal adhesions assemble, attach to the extracellular matrix, and detach dynamically. In the model this turnover is governed by mechanochemical formation rules, polarity-biased attachment kinetics, and force-dependent detachment.

*Formation*

Focal adhesions are created dynamically by each cell, coupling adhesion assembly to both biochemical availability and mechanical load. At every time step each cell estimates the number of adhesions currently associated with it, denoted $N_{\mathrm{FA}}$, by counting focal-adhesion messages addressed to that cell. Formation events are only considered when a refractory timer has elapsed.

Adhesion formation is regulated by two saturating gates. The mechanical gate depends on tensile nuclear stress, $\sigma_+ = \max(0, \sigma_1)$, where $\sigma_1$ is the largest eigenvalue of the nuclear stress tensor. A Hill-type activation function maps this stress into the interval $[0, 1]$,

$$h_\sigma(\sigma_+) = \frac{\sigma_+^{n_\sigma}}{K_\sigma^{n_\sigma} + \sigma_+^{n_\sigma}}.$$

In parallel a biochemical gate is computed from the concentration $c$ of a selected chemical species,

$$h_c(c) = \frac{c^{n_c}}{K_c^{n_c} + c^{n_c}}.$$

The overall formation activation is taken as the product $h_{\mathrm{birth}} = h_\sigma h_c$, and the instantaneous formation rate is $k_{\mathrm{birth}} = k_0 + k_{\max} h_{\mathrm{birth}}$,, which produces the probability of formation within one time step $\Delta t$:

$$P_{\mathrm{birth}} = 1 - \exp(-k_{\mathrm{birth}} \Delta t).$$

A target adhesion number $N_{\mathrm{FA}}^*$ is defined by interpolating between a prescribed minimum $N_{\min}$ and maximum $N_{\max}$ according to the activation level $h_{\mathrm{birth}}$. Formation attempts only occur when $N_{\mathrm{FA}} < N_{\mathrm{FA}}^*$, ensuring that adhesion numbers remain bounded. New adhesions are initialized detached near the cell leading edge,

$$\mathbf{x}_{\mathrm{lead}} = \mathbf{x}_c + R_{\mathrm{cell}}\mathbf{o}.$$

and are assigned to the closest anchor point on a discrete set of reference directions on the nuclear surface.

*Polarity-biased attachment*

Adhesion kinetics depend on the relative orientation between the cell polarity vector $\mathbf{o}$ and the vector from the cell centre to the anchor point $\mathbf{a} = \mathbf{x}_i - \mathbf{x}_c$.

$$p = \frac{\mathbf{o} \cdot \mathbf{a}}{\|\mathbf{o}\| \|\mathbf{a}\|}, \qquad p \in [-1, 1].$$

Positive values correspond to front-oriented adhesions and negative values correspond to rear-oriented adhesions. The score is decomposed into front and rear contributions $p_{\text{front}} = \max(0, p)$ and $p_{\text{rear}} = \max(0, -p)$. Polarity increases the attachment rate at the leading edge, $k_{\text{on}}^{\text{eff}} = k_{\text{on}} \left(1 + G_{\text{front}} p_{\text{front}}\right)$, with the attachment probability defined as:

$$P_{\text{on}} = 1 - \exp(-k_{\text{on}}^{\text{eff}} \Delta t).$$

*Force-dependent detachment*

The baseline detachment rate is also polarity dependent,

$$k_{\text{off},0}^{\text{eff}} = k_{\text{off},0} \left(1 - R_{\text{front}} p_{\text{front}} + G_{\text{rear}} p_{\text{rear}}\right).$$

where $R_{\text{front}}$ controls the reduction of detachment rates at the cell front, and $G_{\text{rear}}$ increases detachment rates for rear-oriented adhesions.

Once attached, detachment becomes force dependent under either slip-bond or catch-slip regimes (user-defined). Let $F$ denote the magnitude of the traction force transmitted through the adhesion. The parameter $F_c$ is a characteristic force scale for slip-bond rupture. In the catch–slip formulation, $a_c$ and $a_s$ weight the catch and slip contributions, while $F_{\text{catch}}$ and $F_{\text{slip}}$ define the corresponding characteristic force scales.

*Slip-bond regime*:

$$k_{\text{off}}(F) = k_{\text{off},0}^{\text{eff}} \exp\left(\frac{|F|}{F_c}\right).$$

*Catch–slip regime*:

$$k_{\text{off}}(F) = k_{\text{off},0}^{\text{eff}} \left(a_c e^{-|F|/F_{\text{catch}}} + a_s e^{|F|/F_{\text{slip}}}\right).$$

In both cases the probability of detachment within a time step is

$$P_{\text{off}} = 1 - \exp(-k_{\text{off}} \Delta t).$$

***Mechanics***

Mechanical forces transmitted through focal adhesions arise from actomyosin contractility within the cell and from elastic resistance in the adhesion–cytoskeleton–nucleus system. The model therefore separates three components: active contractility, force transmission through the adhesion, and optional viscoelastic coupling to the nucleus.

*Active contractility*

When actomyosin contractility is engaged, the adhesion actively shortens its rest length. This represents myosin-driven retrograde actin flow pulling the adhesion toward the cell interior. The rest length $L(t)$ therefore decreases at rate $v_c$,

$$L(t + \Delta t) = \max(L_{\text{min}}, L(t) - v_c \Delta t).$$

The lower bound $L_{\text{min}}$ prevents the adhesion rest length from collapsing to zero and represents geometric limits of the molecular linkage.

*Force transmission*

Let $\ell$ denote the current distance between the adhesion anchor on the nucleus surface and the attached fibre node. In the simplest representation the focal adhesion behaves as a tension-only linear spring,

$$F = k_{\mathrm{FA}} \max(0, \ell - L),$$

where $k_{\mathrm{FA}}$ is the adhesion stiffness and $L$ is the current rest length. This formulation ensures that attached adhesions only transmit tensile forces and cannot push on the extracellular matrix. It is worth noting that CELL agents themselves push FNODE agents away when within an interaction distance, and de-attached adhesions move away from the cell centre, probing the matrix for a potential attachment.

*LINC-mediated nuclear coupling*

In many cells, forces transmitted through focal adhesions are not only balanced by the cytoskeleton but are also coupled to the nucleus through the LINC complex. This protein assembly mechanically connects cytoskeletal filaments to the nuclear envelope, allowing forces generated at adhesions to deform the nucleus and participate in mechanotransduction.

To represent this pathway, the model optionally introduces a viscoelastic LINC element between the nucleus anchor point and the focal adhesion spring. The total length $\ell$ is decomposed into the length of the LINC element $L_0$ and the focal adhesion spring $L_1$ ($\ell = L_0 + L_1$). Force continuity along the series elements yields

$$F = k_{\mathrm{FA}}(L_1 - L)$$

for the focal adhesion spring and

$$F = k_{\mathrm{LINC}}(L_0 - L_{\mathrm{LINC}}) + d_{\mathrm{LINC}}\dot{L}_0$$

for the Kelvin–Voigt LINC element, where $k_{\mathrm{LINC}}$ and $d_{\mathrm{LINC}}$ denote its elastic stiffness and viscous damping coefficient. Using backward Euler discretization for the LINC deformation rate $\dot{L}_0$, the updated LINC length becomes:

$$L_0^{n+1} = \frac{k_{\mathrm{FA}}(\ell^{n+1} - L^{n+1}) + k_{\mathrm{LINC}}L_{\mathrm{LINC}} + \dfrac{d_{\mathrm{LINC}}}{\Delta t}L_0^n}{k_{\mathrm{FA}} + k_{\mathrm{LINC}} + \dfrac{d_{\mathrm{LINC}}}{\Delta t}}.$$

When the LINC coupling is disabled the model reduces to the simpler focal adhesion spring described above. This option is useful when nuclear mechanics are not expected to significantly influence adhesion forces.

***Maturation and reinforcement***

Focal adhesions are known to strengthen under mechanical load through recruitment of additional structural proteins such as talin and vinculin. To represent this maturation process, adhesion stiffness evolves according to a bounded reinforcement law,

$$\frac{dk_{\mathrm{FA}}}{dt} = k_{\mathrm{reinf}}\frac{F}{F + F_{\mathrm{reinf}}}, \qquad k_{\mathrm{FA}} \le k_{\mathrm{FA}}^{\max}.$$

This formulation increases stiffness with increasing load but saturates at a maximum value $k_{\mathrm{FA}}^{\max}$, preventing unbounded stiffening.

#### A.3. Cell migration

Cell migration is implemented as an explicit Euler update of cell position using a velocity vector assembled from multiple motility cues. At each simulation step, each cell agent stores its position $\mathbf{x} = \mathbf{x}_c = (x, y, z)$ and a polarity (orientation) unit vector $\mathbf{o} = (\mathrm{orx}, \mathrm{ory}, \mathrm{orz})$, which is updated in a separate routine and treated as given during the movement update. The cell position is advanced as:

$$\mathbf{x}(t + \Delta t) = \mathbf{x}(t) + \Delta t\, \mathbf{v}(t),$$

where $\mathbf{v}$ is the resulting velocity after combining persistent self-propulsion, Brownian motility, chemotaxis, chemokinesis, durotaxis, and short-range interaction corrections.

***Velocity assembly with intermediate accumulators***

To avoid ordering effects when multiple cues are active, intermediate accumulators are used. A base velocity $\mathbf{v}_{\text{base}}$ collects components that define an initial speed and direction, namely persistent self-propulsion along the current cell orientation and a stochastic Brownian contribution. Two additional accumulators are defined: (i) a steering vector $\mathbf{s}$ collecting contributions that only modify direction while preserving speed, and (ii) an additive velocity increment $\Delta\mathbf{v}$ collecting contributions that may change both direction and speed. The final velocity is computed as

$$\mathbf{v}_0 = \mathbf{v}_{\text{base}},$$

$$\mathbf{v}_{\text{steered}} = \|\mathbf{v}_0\| \frac{\mathbf{v}_0/\|\mathbf{v}_0\| + \mathbf{s}}{\left\|\mathbf{v}_0/\|\mathbf{v}_0\| + \mathbf{s}\right\|},$$

$$\mathbf{v} = \mathbf{v}_{\text{steered}} + \Delta\mathbf{v}$$

If $\|\mathbf{v}_0\|$ is negligible, a user-defined reference speed is used and the initial direction is taken from $\mathbf{s}$. This formulation ensures that "direction-only" effects do not depend on whether a preceding cue has already increased or decreased the speed.

***Persistent and Brownian motility***

In the absence of external directional cues, cells still exhibit active motility through a persistent self-propulsion term aligned with their current polarity vector, combined with a stochastic Brownian component. The base velocity is modulated by a dimensionless chemokinesis factor $f_{\text{ck}}$ (defined below) and is written as:

$$\mathbf{v}_{\text{base}} = (f_{\text{ck}}\, v_{\text{ref}})\, \mathbf{o} + B\, \boldsymbol{\eta},$$

where $v_{\text{ref}}$ is the cell-specific reference speed stored by each agent, $B$ is a cell type-specific Brownian strength parameter, and $\boldsymbol{\eta} = (\eta_x, \eta_y, \eta_z)$ is a vector of three independent random samples drawn uniformly from $[-1, 1]$ at each time step.

The persistent term introduces directional memory by driving motion along the current cell orientation, whereas the Brownian term introduces random exploratory fluctuations. The balance between the modulated speed $f_{\text{ck}}\, v_{\text{ref}}$ and $B$ controls the degree of migration persistence.

***Rotational diffusion***

Because the orientation vector $\mathbf{o}$ is otherwise only updated by the mechanical reorientation mechanism (see Section A.3), free-migrating cells (i.e. when mechanical cues are deactivated) would follow nearly ballistic trajectories. A rotational diffusion step is therefore applied to the orientation vector at each time step before computing $\mathbf{v}_{\text{base}}$:

$$\mathbf{o}^* = \mathbf{o}^n + \sigma_{\text{rot}}\, \boldsymbol{\eta}, \qquad \mathbf{o}^{n+1} = \frac{\mathbf{o}^*}{|\mathbf{o}^*|},$$

where $\boldsymbol{\eta}$, as before, is randomly sampled and the noise amplitude is

$$\sigma_{\text{rot}} = \sqrt{2\, D_{\text{rot}}\, \Delta t},$$

with $D_{\text{rot}}$ the cell type-specific rotational diffusion coefficient (units of $\text{rad}^2/\text{s}$).

This produces a persistent random walk whose directional correlation decays with a characteristic persistence time $\tau_p \sim 1/(2\, D_{\text{rot}})$. For $D_{\text{rot}} = 0$ the orientation remains fixed, while increasing $D_{\text{rot}}$ yields progressively more tortuous trajectories that eventually approach pure diffusive motion.

***Chemokinesis and speed adaptation***

Chemokinesis models the non-directional modulation of a cell's persistent speed in response to absolute local chemical concentrations. The modulation factor $f_{\text{ck}}$ incorporates soft signal saturation, temporal adaptation, and bounded Hill gating across multiple chemical species.

For a given cell at position $\mathbf{x}$, each chemical species $s$ elicits a raw local signal weighted by a species-specific chemokinesis sensitivity $\kappa_s$. A positive sensitivity ($\kappa_s > 0$) contributes to a promotive channel that increases speed, whereas a negative sensitivity ($\kappa_s < 0$) contributes to an inhibitory channel that decreases speed. The raw absolute signal is defined as:

$$u_s = |\kappa_s| C_s(\mathbf{x}),$$

where $C_s(\mathbf{x})$ denotes the concentration of species $s$ at the grid location corresponding to the cell position. To account for receptor saturation, the signal undergoes a soft, Michaelis-Menten-like saturation:

$$\tilde{u}_s = C_{\text{sat},s} \frac{u_s}{C_{\text{sat},s} + u_s},$$

where $C_{\text{sat},s}$ is the maximum saturation level for species $s$.

Cells adapt to sustained chemical stimuli over a characteristic timescale $\tau$. This is modeled using an internal adaptation state variable $A_s(t)$ for each species, which evolves according to:

$$\frac{dA_s}{dt} = \frac{1}{\tau}(\tilde{u}_s - A_s).$$

In the discrete implementation, $A_s$ is updated via an explicit Euler step. The effective, desensitized signal driving the cell's chemokinetic response is then:

$$u_s^* = \tilde{u}_s - A_s$$

The effective signals are linearly summed into total promotive and inhibitory pools depending on the sign of their respective sensitivities:

$$S_P = \sum_{\{s|\kappa_s>0\}} u_s^*, \qquad S_I = \sum_{\{s|\kappa_s<0\}} u_s^*.$$

Finally, these pools are passed through a bounded Hill function, $h_{ck}(S) = S^n/(K_{ck}^n + S^n)$, where $K_{ck}$ is the half-activation threshold and $n$ is the Hill coefficient. The overall chemokinetic multiplier modifying the reference speed is computed as:

$$f_{\text{ck}} = 1 + \alpha_{ck} \left[ h_{ck}(S_P) - h_{ck}(S_I) \right],$$

where $\alpha_{ck}$ is a global amplitude parameter controlling the maximum fractional increase or decrease in speed. If the final multiplier drops below zero, it is truncated to ensure cell speed remains non-negative.

##### *Chemotaxis from macro-scale concentration fields*

Chemotaxis is driven by gradients of macro-scale chemical concentrations fields of the ECM grid. As before, $C_s(\mathbf{x})$ denotes the concentration of species $s$ at the grid location corresponding to the cell position. The cell position is mapped to grid indices $(i, j, k)$, and a 26-neighborhood box was used to approximate a direction of steepest ascent or descent.

For each neighbor offset $(d_i, d_j, d_k) \in \{-1, 0, 1\}^3 \setminus \{(0, 0, 0)\}$, the physical displacement to the neighbor is

$$\Delta\mathbf{r} = (d_i\,\Delta x,\ d_j\,\Delta y,\ d_k\,\Delta z),$$

with grid spacings $(\Delta x, \Delta y, \Delta z)$. A distance-weighted, multi-species gradient proxy was computed as

$$\mathbf{g} = \sum_{(d_i,d_j,d_k)} \left( \frac{1}{\|\Delta\mathbf{r}\|} \sum_{s=1}^{N_s} \chi_s \left[ C_s(\mathbf{x} + \Delta\mathbf{r}) - C_s(\mathbf{x}) \right] \right) \frac{\Delta\mathbf{r}}{\|\Delta\mathbf{r}\|},$$

where $N_s$ is the number of species and $\chi_s \in [-1, 1]$ is a per-cell type chemotactic sensitivity that allows attraction ($\chi_s > 0$), repulsion ($\chi_s < 0$), or deactivation ($\chi_s = 0$) per species. The chemotactic unit direction is therefore

$$\hat{\mathbf{g}} = \frac{\mathbf{g}}{\|\mathbf{g}\|}$$

Chemotaxis contributes either as a direction-only steering term (added to $\mathbf{s}$) or as a speed-changing increment (added to $\Delta\mathbf{v}$), depending on a user-defined configuration flag:

$$\mathbf{s} \mathrel{+}= \chi\,\hat{\mathbf{g}} \qquad \text{(direction-only mode)}$$

$$\Delta\mathbf{v} \mathrel{+}= \chi\,\hat{\mathbf{g}} \qquad \text{(speed-changing mode)}$$

where $\chi$ is a global chemotaxis strength parameter.

***Durotaxis from mechanical state variables***

Durotaxis is implemented as a preference to move toward mechanically "stiffer" directions, represented here by stress and strain tensors stored per cell,

$$\boldsymbol{\sigma} = \begin{bmatrix} \sigma_{xx} & \sigma_{xy} & \sigma_{xz} \\ \sigma_{xy} & \sigma_{yy} & \sigma_{yz} \\ \sigma_{xz} & \sigma_{yz} & \sigma_{zz} \end{bmatrix}, \qquad \boldsymbol{\varepsilon} = \begin{bmatrix} \varepsilon_{xx} & \varepsilon_{xy} & \varepsilon_{xz} \\ \varepsilon_{xy} & \varepsilon_{yy} & \varepsilon_{yz} \\ \varepsilon_{xz} & \varepsilon_{yz} & \varepsilon_{zz} \end{bmatrix}.$$

Principal values and vectors of both tensors are also stored in agent variables:

$$\boldsymbol{\sigma}\,\mathbf{v}^{\sigma}_{m} = \lambda^{\sigma}_{m}\,\mathbf{v}^{\sigma}_{m}, \qquad \boldsymbol{\varepsilon}\,\mathbf{v}^{\varepsilon}_{m} = \lambda^{\varepsilon}_{m}\,\mathbf{v}^{\varepsilon}_{m}, \qquad m \in \{1, 2, 3\},$$

with $\lambda_1 \geq \lambda_2 \geq \lambda_3$ and corresponding unit eigenvectors.

Two complementary durotactic directions were defined: (i) a traction-based direction aligned with the current cell orientation,

$$\mathbf{t} = \boldsymbol{\sigma}\,\mathbf{o}, \qquad \hat{\mathbf{t}} = \frac{\mathbf{t}}{\|\mathbf{t}\|},$$

and (ii) a principal-axis direction given by the dominant eigenvector,

$$\hat{\mathbf{p}} = \begin{cases} \mathbf{v}^{\sigma}_{1}, & \text{stress-based durotaxis,} \\ \mathbf{v}^{\varepsilon}_{1}, & \text{strain-based durotaxis.} \end{cases}$$

To prevent sign flips due to eigenvector ambiguity, $\hat{\mathbf{p}}$ is oriented to maintain continuity with the orientation direction by enforcing $\hat{\mathbf{p}} \cdot \mathbf{o} \geq 0$ (otherwise $\hat{\mathbf{p}} \leftarrow -\hat{\mathbf{p}}$).

In addition to contributing to the migration velocity, the dominant mechanical direction can optionally reorient the cell polarity itself. In this case, the target direction is taken as the first principal eigenvector of either the stress or strain tensor, depending on the selected mode. After resolving the sign ambiguity as above to ensure continuity with the current polarity, the orientation vector is updated by a bounded linear relaxation toward that target,

$$\mathbf{o}^{n+1,*} = (1 - \alpha)\,\mathbf{o}^{n} + \alpha\,\hat{\mathbf{p}}, \qquad \alpha = k_{\text{align}}\,\Delta t$$

followed by normalization,

$$\mathbf{o}^{n+1} = \frac{\mathbf{o}^{n+1,*}}{\|\mathbf{o}^{n+1,*}\|}.$$

Here $k_{\text{align}}$ is a user-defined orientation-alignment rate. This mechanism provides a gradual mechanical alignment of cell polarity with the direction of maximum principal stress or strain, rather than imposing an instantaneous reorientation.

A blended durotactic direction is then defined as

$$\hat{\mathbf{d}} = \frac{(1 - \beta)\hat{\mathbf{t}} + \beta\,\hat{\mathbf{p}}}{\left\|(1 - \beta)\hat{\mathbf{t}} + \beta\,\hat{\mathbf{p}}\right\|},$$

where $\beta \in [0, 1]$ interpolates between traction-only ($\beta = 0$) and principal-axis-only ($\beta = 1$) durotaxis.

The magnitude of the durotactic response is scaled using two complementary quantities derived from the local mechanical state. The first is a positive stress–strain coupling term,

$$E = \max(0,\ \boldsymbol{\sigma} : \boldsymbol{\varepsilon}) = \max\left(0,\ \sigma_{xx}\varepsilon_{xx} + \sigma_{yy}\varepsilon_{yy} + \sigma_{zz}\varepsilon_{zz} + 2(\sigma_{xy}\varepsilon_{xy} + \sigma_{xz}\varepsilon_{xz} + \sigma_{yz}\varepsilon_{yz})\right),$$

which acts as an energy-like measure of mechanical loading and vanishes in compressive or mechanically inactive regions. To account for the directional organization of the mechanical field, a dimensionless anisotropy factor is computed from the principal values $\lambda_1, \lambda_2, \lambda_3$ (obtained from either the stress or strain tensor depending on the selected durotaxis mode). The anisotropy measure is defined as

$$A = \frac{\lambda_1 - \lambda_3}{|\lambda_1| + |\lambda_2| + |\lambda_3|},$$

which increases as the mechanical state becomes more anisotropic and vanishes when the principal values are nearly equal.

The resulting durotactic strength is computed as

$$S_{\text{duro}} = \mu_{\text{duro}} (E + A),$$

where $\mu_{\text{duro}}$ is a global strength parameter controlling the sensitivity of cell migration to mechanical cues.

As with chemotaxis, durotaxis contributes either as a direction-only steering term or as a speed-changing increment:

$$\mathbf{s} \mathrel{+}= S_{\text{duro}} \, \hat{\mathbf{d}} \qquad \text{(direction-only mode)}$$

$$\Delta \mathbf{v} \mathrel{+}= S_{\text{duro}} \, \hat{\mathbf{d}} \qquad \text{(speed-changing mode)}$$

***Short-range mechanical interaction velocity terms***

In addition to Brownian motility, chemotaxis, and durotaxis, cell motion includes two short-range mechanical contributions computed at each time step: a CELL–CELL interaction term $\Delta \mathbf{v}_{cc}$ and a CELL–FNODE exclusion term $\Delta \mathbf{v}_{cf}$. The velocity used in the explicit Euler position update therefore reads

$$\mathbf{v} = \mathbf{v}_{\text{steered}} + \Delta \mathbf{v} + \Delta \mathbf{v}_{cc} + \Delta \mathbf{v}_{cf}.$$

In both interaction models, pairwise mechanical forces are first accumulated and then converted into velocity increments using an effective drag coefficient ($\Delta \mathbf{v} = \mathbf{F}/\gamma$).

*CELL–CELL interactions.* For neighboring cells within the spatial search radius, a pairwise interaction is applied along the inter-cell unit vector. When the distance between cell centers is smaller than the sum of their radii, a linear repulsive force proportional to the overlap depth prevents interpenetration. For separations slightly larger than contact, a weak finite-range attractive interaction promotes aggregate cohesion. The contributions from all neighbors are summed to obtain $\mathbf{F}_{cc}$ and converted to the velocity increment $\Delta \mathbf{v}_{cc}$.

*CELL–FNODE exclusion.* Interactions with nearby fibre nodes are purely repulsive. When a fibre node enters the exclusion region surrounding the cell, a repulsive force is generated along the cell–node direction to maintain a minimum separation. The equal and opposite reaction displaces the fibre node away from the cell. The summed interaction force $\mathbf{F}_{cf}$ is converted to the velocity increment $\Delta \mathbf{v}_{cf}$ using the same drag formulation.

***Boundary handling***

CELL agents are confined to a rectangular computational domain with bounds $x_{i,\min}(t) \leq x_i \leq x_{i,\max}(t), i \in \{x, y, z\}$, where the boundary positions may vary over time depending on the simulation configuration.

When periodic boundary conditions are enabled, positions are wrapped independently along each coordinate direction. If a cell exits the domain, its position is shifted by the corresponding domain length ($L_i = x_{i,\max} - x_{i,\min}$) according to:

$$x_i \leftarrow \begin{cases} x_i + L_i, & x_i < x_{i,\min}, \\ x_i - L_i, & x_i > x_{i,\max}. \end{cases}$$

This operation produces toroidal boundary conditions, allowing cells to migrate continuously across the domain without accumulation at the boundaries. When periodic boundaries are not enabled, cell positions are simply clamped to the domain limits:

$$x_i \leftarrow \min\left(x_{i,\max}, \max\left(x_{i,\min}, x_i\right)\right),$$

Note that periodic wrapping can only be applied when focal adhesions are disabled, since relocating a cell while its adhesion complexes remain anchored to the extracellular matrix on the opposite side of the domain would produce unphysical behaviour.

#### A.4. Cell cycle

Each CELL agent evolves according to an internal cell-cycle program that controls growth, division, and survival. The framework provides a generic template based on the canonical eukaryotic phases (G1, S, G2, and M), represented by a scalar clock variable $t_c$ that advances with the simulation time step $\Delta t$,

$$t_c(t + \Delta t) = t_c(t) + \Delta t.$$

Phase boundaries are specified through user-defined start times, allowing the current phase of each cell to be determined by comparing $t_c$ with these thresholds. The template provides a structural organization of the cycle, while the specific biological behaviours associated with each phase (e.g., growth, checkpoints, DNA replication, or mitosis) can be defined and customized by the user for each cell type. This design enables flexible modelling of diverse cell-cycle dynamics without imposing a fixed biological interpretation.

***Cell growth prior to mitosis***

Following division, cells begin at approximately half their baseline size and gradually grow during interphase until reaching their reference dimensions prior to mitosis. Both the cell radius $R_{\text{cell}}$ and the nucleus radius $R_{\text{nuc}}$ increase linearly in time until the start of M phase at time $T_M$:

$$R_{\text{cell}}(t + \Delta t) = R_{\text{cell}}(t) + \frac{R^0_{\text{cell}}}{2T_M}\,\Delta t$$

$$R_{\text{nuc}}(t + \Delta t) = R_{\text{nuc}}(t) + \frac{R^0_{\text{nuc}}}{2T_M}\,\Delta t$$

where $R^0_{\text{cell}}$ and $R^0_{\text{nuc}}$ denote the baseline radii of the cell and nucleus.

***Stochastic division during M phase***

Once a cell enters the M phase, division is treated as a stochastic event that must occur during this phase but whose exact timing is random. Let

$$t_M = t_c - T_M$$

denote the elapsed time since entry into M phase, and let $T_{M,\text{dur}}$ be the duration of this phase. The corresponding number of discrete simulation steps is

$$N_M = T_{M,\text{dur}}/\Delta t.$$

The division time is drawn from a discrete uniform distribution over the $N_M$ steps of the M phase. Equivalently, the probability of division at step $t$ is computed conditionally on the event that division has not yet occurred. The conditional probability of division at the current step therefore reads:

$$P(D^c_t \mid \cap_{i=1}^{t-1}\overline{D^c_i}) = \frac{1/N_M}{(N_M - t + 1)/N_M} = \frac{1}{N_M - t + 1},$$

which produces a monotonically increasing division probability as the number of remaining steps decreases. This conditional formulation guarantees that the division time is uniformly distributed across the discrete steps of the M phase and is therefore independent of the simulation time step. Cell-type-dependent multipliers may further scale the effective division probability.

When division occurs, the parent cell and daughter cell are placed symmetrically along the cell orientation vector $\mathbf{o}$:

$$\mathbf{x}'_{\text{parent}} = \mathbf{x} + \frac{R^0_{\text{cell}}}{2}\mathbf{o}$$

$$\mathbf{x}_{\text{daughter}} = \mathbf{x} - \frac{R^0_{\text{cell}}}{2}\mathbf{o}.$$

*Partitioning of internal state*

Following division, internal variables are redistributed between the parent and daughter cells, who are assigned unique identifiers through an atomic increment of a global counter.

Some scalar internal quantities are partitioned symmetrically. For example, the cumulative damage variable (defined in the next subsection) is divided equally between the two cells. Per-cell chemical masses $M_s$ are halved while concentrations $C_s$ remain unchanged:

$$M_s \leftarrow \frac{1}{2}M_s \qquad C_s \leftarrow C_s,$$

Dynamic and mechanical state variables, including velocity, stress, and strain, are reset to zero following division. In addition, the FOCAD agents previously associated with the parent cell are randomly partitioned between the parent and daughter cell so that both inherit a subset of the pre-existing adhesion complexes.

To prevent artificial synchronization of the cell population, the parent cell-cycle clock is reset to a small random offset within early G1. A completed-cycle counter stored within the parent agent is incremented for bookkeeping.

##### *Cumulative damage and death pathways*

Cell death can occur either through acute lethal conditions or through progressive damage accumulation. Acute death may be triggered by extreme microenvironmental conditions such as severe hypoxia, extreme nutrient deprivation, or excessive mechanical stress.

In addition, each cell maintains a scalar damage variable $d \in [0, 1]$ that integrates sub-lethal stress over time while allowing partial recovery. Damage accumulation depends on three environmental proxies: oxygen availability $O$, nutrient availability $N$, and tensile stress magnitude $S$, defined as

$$S = \max(0, \sigma_1),$$

where $\sigma_1$ is the maximum principal stress of the nucleus.

Damage increases when oxygen or nutrient levels fall below prescribed thresholds and when tensile stress exceeds its threshold. Defining normalized severities

$$\mathrm{sev}_O = \frac{O_{\mathrm{th}} - O}{O_{\mathrm{th}}},\ \mathrm{sev}_N = \frac{N_{\mathrm{th}} - N}{N_{\mathrm{th}}},\ \mathrm{sev}_S = \frac{S - S_{\mathrm{th}}}{S_{\mathrm{th}}}$$

and considering only positive severities, the damage update reads

$$d(t + \Delta t) = d(t) + \Delta t \left[k_O\, \mathrm{sev}_O^+ + k_N\, \mathrm{sev}_N^+ + k_S\, \mathrm{sev}_S^+ - k_{\mathrm{rep}}\right]$$

where $k_O$, $k_N$, and $k_S$ are accumulation rates for the respective damage pathways, and $k_{\mathrm{rep}}$ represents basal repair. Cell-type-dependent multipliers may modulate damage accumulation, repair rates, and death thresholds to represent phenotypic heterogeneity.

When a cell is marked as dead, its velocity is set to zero and a death-cause code is recorded for downstream analysis. Importantly, dead cells remain in the simulation by default, and continue interacting mechanically with other agents. A user-defined flag can be activated to, instead, eliminate them from the simulation after death in order to reduce computational cost.

#### A.5. Diffusion of species and cell metabolism

Extracellular chemical species are represented on a structured Cartesian lattice of ECM agents that coexist with the CELL agent population. Each ECM agent stores the concentrations of $N_s$ diffusing species and interacts locally with cells occupying the same spatial region. For clarity of exposition, ECM agents will be referred to in this section as *voxels*, although the grid is implemented explicitly as a population of agents rather than a fixed numerical mesh.

This agent-based representation allows the positions of ECM agents to be adapted when domain boundaries move, enabling deformation of the diffusion lattice without requiring remeshing. Within each simulation step the model sequentially updates extracellular diffusion, CELL-ECM exchange, and intracellular metabolic reactions.

##### *Reaction–diffusion formulation*

Let $C_s(\mathbf{x}, t)$ denote the extracellular concentration of species $s \in \{1, \dots, N_s\}$ stored by each ECM voxel (agent) located at position $\mathbf{x}$ on a structured lattice with spacings $(\Delta x, \Delta y, \Delta z)$. The governing reaction–diffusion equation is:

$$\frac{\partial C_s}{\partial t} = D_s \nabla^2 C_s + R_s^{\mathrm{cells}} + R_s^{\mathrm{bg}}$$

where $D_s$ is the diffusion coefficient, $R_s^{\mathrm{cells}}$ represents cell-mediated exchange, and $R_s^{\mathrm{bg}}$ denotes optional background reactions. Using second-order finite differences and a forward Euler time update, the discrete evolution for voxel $(i, j, k)$ reads:

$$C_{i,j,k}^{t+1} = C_{i,j,k}^t + F_x(C_{i+1,j,k}^t - 2C_{i,j,k}^t + C_{i-1,j,k}^t) + F_y(C_{i,j+1,k}^t - 2C_{i,j,k}^t + C_{i,j-1,k}^t) + F_z(C_{i,j,k+1}^t - 2C_{i,j,k}^t + C_{i,j,k-1}^t) + R_{i,j,k}^t \Delta t$$

where the diffusion numbers are:

$$F_x = \frac{D_s \Delta t}{\Delta x^2}, \qquad F_y = \frac{D_s \Delta t}{\Delta y^2}, \qquad F_z = \frac{D_s \Delta t}{\Delta z^2}.$$

Stability of the explicit scheme requires:

$$F_x + F_y + F_z \leq \frac{1}{2}.$$

When this condition is violated, a centre-weighted semi-implicit update is applied instead, where neighbouring voxel concentrations are evaluated at time $t$ while the central term is treated implicitly:

$$\left(1 + 2(F_x + F_y + F_z)\right) C_{i,j,k}^{t+1} = C_{i,j,k}^{t} + F_x \left(C_{i+1,j,k}^{t} + C_{i-1,j,k}^{t}\right) + F_y \left(C_{i,j+1,k}^{t} + C_{i,j-1,k}^{t}\right) + F_z \left(C_{i,j,k+1}^{t} + C_{i,j,k-1}^{t}\right) + R_{i,j,k}^{t} \Delta t$$

Importantly, two alternative formulations are considered for the diffusion coefficient $D_s$, allowing the model to represent either homogeneous transport or transport hindered by the local extracellular matrix microstructure.

*Constant diffusion:* In the simplest formulation, the diffusion coefficient $D_s$ for each species is taken as constant throughout the domain. This assumption corresponds to a homogeneous medium and is appropriate when the ECM is either absent, weakly structured, or when transport limitations due to fibre architecture are not expected to play a dominant role.

*Fibre-dependent diffusion:* To account for the effect of matrix microstructure on transport, the diffusion coefficient can be locally modulated as a function of fibre density. This reflects the physical observation that dense fibrous networks hinder molecular diffusion through steric obstruction and increased tortuosity.

In this formulation, a local fibre density measure $\rho(\mathbf{x})$ is estimated by counting FNODEs within each ECM voxel and normalizing by an average reference density (computed globally at the start of the simulation). The effective diffusion coefficient is then scaled as

$$D_s^{\text{eff}}(\mathbf{x}) = D_s \, m(\rho),$$

where $m(\rho) \in (0, 1]$ is a decreasing function of density. A saturating mapping is used so that diffusion remains unchanged at low densities and progressively decreases as the local fibre concentration increases,

$$m(\rho) = \frac{1}{1 + \alpha_h \, \max(\rho - 1, 0)},$$

with an additional lower bound to prevent complete arrest of diffusion. The parameter $\alpha_h$ controls the strength of the hindrance effect.

This formulation enables spatially heterogeneous transport properties to emerge directly from the evolving fibre network, allowing dense regions to act as diffusion barriers while maintaining efficient transport in sparse regions.

##### *CELL-ECM exchange*

Cells exchange chemical species with the surrounding extracellular ECM voxels through production and uptake processes. For a voxel containing at least one cell, the extracellular kinetics of species $s$ are written as

$$\frac{dC_s}{dt} = \alpha \left(k_p C_{s,\text{sat}} - (k_p + k_c) C_s\right),$$

where $k_p$ and $k_c$ are cell production and consumption rates, $C_{s,\text{sat}}$ is a saturation concentration, and $\alpha = V_{\text{cell}}/V_{\text{ECM}}$ with $V_{\text{ECM}} = \Delta x \, \Delta y \, \Delta z$. Applying a backward Euler discretization yields the closed-form update:

$$C_s^{t+1} = \frac{C_s^t + \Delta t \, \alpha \, k_p \, C_{s,\text{sat}}}{1 + \Delta t \, \alpha \, (k_p + k_c)}.$$

This formulation is unconditionally stable and preserves non-negative concentrations.

*Mass conservation*

For each species $s$, the extracellular mass contained in a voxel is

$$M_{s,\mathrm{ECM}} = C_s V_{\mathrm{ECM}}.$$

Let $C_{s,\mathrm{prop}}^{t+1}$ denote the concentration obtained from the reaction update described above. The corresponding proposed mass exchange during the time step is

$$\Delta M_{s,\mathrm{ECM}}^{\mathrm{prop}} = \left(C_{s,\mathrm{prop}}^{t+1} - C_s^t\right) V_{\mathrm{ECM}}.$$

To prevent negative concentrations and ensure local mass conservation, the actual exchanged mass is limited by the available mass in both compartments. The accepted exchange therefore satisfies

$$-M_{s,\mathrm{ECM}}^t \le \Delta M_{s,\mathrm{ECM}} \le M_{s,\mathrm{cell}}^t,$$

and is chosen as the closest admissible value to the proposed exchange $\Delta M_{s,\mathrm{ECM}}^{\mathrm{prop}}$.

The intracellular mass of species $s$ is then updated as

$$M_{s,\mathrm{cell}}^{t+1} = M_{s,\mathrm{cell}}^t - \Delta M_{s,\mathrm{ECM}}.$$

This formulation guarantees local mass conservation between the cell and its surrounding ECM while preventing negative extracellular concentrations or intracellular depletion beyond the available mass. Notably, all updates are performed atomically at the voxel level, so that multiple cells interacting with the same voxel during a time step may compete for the available extracellular mass.

***Intracellular metabolism***

Intracellular species are assumed to be well mixed within each cell (i.e. spatial gradients inside the cell are neglected). Their concentrations evolve according to

$$\frac{dC_s^{\mathrm{cell}}}{dt} = \mathcal{F}_s\left(C^{\mathrm{cell}}, C^{\mathrm{ecm}}, \boldsymbol{\sigma}, \boldsymbol{\varepsilon}, \text{cell state}\right),$$

where $\mathcal{F}_s$ denotes a user-defined reaction operator describing the intracellular kinetics of species $s$. This operator may incorporate metabolic pathways, signaling reactions, or mechanically regulated biochemical processes.

***Modularity***

The number of species $N_s$, diffusion coefficients $D_s$, and reaction operators $\mathcal{F}_s$ are user-configurable. This allows the framework to represent processes such as oxygen transport, nutrient uptake, signaling gradients, waste accumulation, synthetic reaction–diffusion systems, and heterogeneous metabolic phenotypes. The framework therefore provides a flexible scaffold rather than a fixed biochemical model.

#### A.6. Fibre network

As an optional component of the framework, the extracellular matrix can be represented as a three-dimensional disordered fibre network composed of interconnected nodes (FNODE agents) that also interact with CELL and FOCAD agents. This feature is used when the biological application requires an explicit fibrous scaffold, but it can be omitted in models where such structure is not needed.

***Network generation***

The network generation pipeline follows and extends previously published approaches for collagen-like and biopolymer networks [21–23], and is implemented here in Python for direct integration with the present agent-based matrix representation.

Briefly, nodes are distributed throughout a rectangular domain according to a prescribed density, with additional boundary-associated nodes introduced when anchoring at selected faces is required. Fibre connectivity is then established by linking nearby nodes while enforcing a prescribed valency distribution. The resulting network is subsequently refined through stochastic optimization steps that regularize fibre lengths and improve local branching geometry. Optional boundary-processing operations are then applied, including snapping of near-boundary nodes to selected planes and pruning of artificial in-plane connections. Finally, long fibres can be subdivided into shorter segments of prescribed maximum length to improve fibre representation accuracy. The final output consists of node coordinates and adjacency information suitable for direct use in subsequent simulations. Full algorithmic details and representative diagnostics are provided in Appendix C.

##### *Fibre mechanics*

The extracellular matrix fibre network is represented by FNODE agents connected by discrete fibre segments. In addition to passive mechanics, the network can remodel through (i) degradation and reinforcement driven by nearby cells and (ii) cell-driven secretion of new FNODEs (representing deposition or local fibre nucleation). Remodelling is controlled by a global enable flag, so that the full remodelling pipeline can be activated or deactivated without changing other model components.

Mechanical interactions between connected FNODE pairs represent individual fibre segments and are modelled using a Kelvin–Voigt formulation, that is, a linear elastic spring acting in parallel with a viscous dashpot. For a connected pair of nodes $i$ and $j$, with current separation $\ell_{ij}$ and equilibrium length $\ell^0_{ij}$, the axial force along the fibre direction is given by

$$F_{ij} = k^{\text{eff}}_{ij}\,(\ell_{ij} - \ell^0_{ij}) + d\,\dot{\ell}_{ij},$$

where $d$ is a viscous damping coefficient and $\dot{\ell}_{ij}$ is the relative velocity projected along the fibre axis. The dashpot term provides rate-dependent dissipation and improves numerical stability of the network dynamics.

###### *Remodelling-dependent stiffness*

The stiffness of each fibre segment is dynamically modulated by local remodelling state variables, so that degradation reduces load-bearing capacity whereas reinforcement increases the effective elastic response of the matrix. The nodal stiffness is defined as:

$$k_i = k_0(1 - \phi_d + \phi_r)$$

where $k_0$ denotes the baseline fibre-segment stiffness, and $\phi_d$ and $\phi_r$ are bounded degradation and reinforcement variables, respectively (see next section).

For a fibre segment connecting two nodes, the effective stiffness is computed as the equivalent stiffness of two springs in series:

$$k_{ij} = \frac{k_i k_j}{k_i + k_j}$$

###### *Strain-dependent nonlinear response*

To capture the non-linear mechanical behaviour of fibrous biopolymer networks, the segment stiffness is further modulated by the local axial strain:

$$\varepsilon_{ij} = \frac{\ell_{ij} - \ell^0_{ij}}{\ell^0_{ij}}.$$

A piecewise exponential law is used to represent fibre buckling under compression and strain stiffening under tension [38]:

$$k^{\text{eff}}_{ij} = k_{ij}\, f(\varepsilon_{ij})$$

where the strain-dependent factor $f(\varepsilon)$ is defined as

$$f(\varepsilon) = \begin{cases} \exp\left(\dfrac{\varepsilon}{d_0}\right), & \varepsilon < 0, \\ 1, & 0 \le \varepsilon \le \varepsilon_s, \\ \exp\left(\dfrac{\varepsilon - \varepsilon_s}{d_s}\right), & \varepsilon > \varepsilon_s, \end{cases}$$

with $d_0$ controlling the rate of stiffness decay in compression (buckling regime), $\varepsilon_s$ the onset strain for stiffening, and $d_s$ the rate of exponential stiffening. The stiffness increase is capped to prevent unbounded growth.

This formulation enables the network to exhibit asymmetric mechanical behaviour, with reduced resistance under compression and increased stiffness under tension, consistent with experimental observations in collagen and other biological networks.

***Degradation and reinforcement from nearby cells***

Each FNODE maintains two remodeling state variables: degradation level $\phi_d \in [0, 1]$ and reinforcement level $\phi_r \geq 0$. The reinforcement variable accounts for deposition of new matrix material, cross-linking, or other strengthening processes defined by the user. By default, both variables are initialized to zero.

At each step, a FNODE counts the number of live cells ($n_{\text{cell}}$) within a prescribed radius, and degradation and reinforcement evolve proportionally to local cellular presence,

$$\phi_d(t + \Delta t) = \phi_d(t) + \Delta t \, k_{\text{deg}} \, n_{\text{cell}},$$

$$\phi_r(t + \Delta t) = \phi_r(t) + \Delta t \, k_{\text{dep}} \, n_{\text{cell}},$$

where $k_{\text{deg}}$ is the degradation rate and $k_{\text{dep}}$ is the deposition (reinforcement) rate, which may depend on cell type.

FNODEs are removed from the simulation once their degradation exceeds unity, representing complete local loss of structural integrity.

***Deposition and secretion of new matrix fibres***

Live CELL agents can probabilistically generate new fibre nodes (FNODEs), representing the local deposition or secretion of extracellular matrix material. To avoid unrealistically frequent deposition events, each CELL maintains a refractory timer $T_{\text{cool}}$, which is decremented by $\Delta t$ and must reach zero before another birth event can occur. At most one FNODE can be generated per cell and simulation step.

The birth propensity is modulated by both mechanical and biochemical signals. As a mechanical proxy we use the positive part of the maximum principal stress in the nucleus,

$$\sigma_+ = \max(0, \sigma_1),$$

where $\sigma_1$ is the largest eigenvalue of the nucleus stress tensor. The biochemical proxy is the non-negative concentration $c$ of a user-selected extracellular species,

$$c = \max(0, c_{\text{raw}}).$$

Each signal is mapped to a saturating Hill response function,

$$h_\sigma(\sigma_+) = \frac{\sigma_+^{n_\sigma}}{K_\sigma^{n_\sigma} + \sigma_+^{n_\sigma}}, \qquad h_c(c) = \frac{c^{n_c}}{K_c^{n_c} + c^{n_c}},$$

where $K_\sigma$ and $K_c$ are activation thresholds and $n_\sigma$ and $n_c$ are Hill exponents controlling the steepness of the response.

The combined activation level is defined multiplicatively as

$$h_{\text{birth}} = h_\sigma \, h_c,$$

so that both mechanical and biochemical cues must be present to strongly promote fibre deposition.

The effective birth rate is then

$$k_{\text{birth}} = k_0 + k_{\max} \, h_{\text{birth}},$$

where $k_0$ represents a baseline deposition rate and $k_{\max}$ sets the maximal additional rate achievable under full activation.

Assuming a Poisson process for FNODE creation, the corresponding per-step probability of a birth event during a time step $\Delta t$ is

$$P_{\text{birth}} = 1 - \exp\left(-k_{\text{birth}} \Delta t\right).$$

If a uniform random draw is below $P_{\text{birth}}$, the CELL emits a newborn FNODE agent and resets the refractory timer to its prescribed cooldown value.

*Integration of newborn FNODE agents*

The newborn FNODE is placed at a fixed radial distance $R_{\mathrm{birth}}$ from the cell center along a random unit direction $\hat{\mathbf{d}}$:

$$\mathbf{x}_{\mathrm{new}} = \mathbf{x}_c + R_{\mathrm{birth}}\hat{\mathbf{d}}, \qquad \|\hat{\mathbf{d}}\| = 1.$$

To integrate the newborn node into the existing network, the CELL selects up to two nearby potential partner FNODEs using spatial neighbor queries. Candidates must (i) lie within a maximum link distance $R_{\mathrm{link}}$ and (ii) have available connectivity capacity (degree strictly below a maximum connectivity). For each selected partner $p$, an initial equilibrium distance is set to the current separation to create initially stress-free links:

$$\ell_{\mathrm{eq}}^{(p)} \leftarrow \left\| \mathbf{x}_{\mathrm{new}} - \mathbf{x}_p \right\|$$

#### B. Configurator and Editor UI

Cellfoundry provides two complementary user interfaces to facilitate model development and exploration: a configuration interface (configurator UI) for generating new FLAMEGPU2 models in general, and an editor interface (editor UI) for interacting with existing ones.

The configurator UI is designed as a rapid template-generation tool that allows users to define the main components of a model in a structured and visual manner (Fig. Supp.Fig.1). In addition to specifying agent types, variables, and functions, a key feature is the ability to manually and graphically connect agents to represent their interactions. This visual linking reflects the underlying communication and dependency structure of the model, providing an intuitive overview of how different subsystems are coupled. The interface supports saving and loading configurations, and also allows importing existing FLAMEGPU2 Python model files, including those not originally created with the tool, to be used as templates. Based on the configured elements, the tool automatically generates a consistent project structure (a main .py file including model configuration, and a set of .cpp template files to define agent behavior) with the required code and parameter definitions, serving as a starting point for further development. An example workflow is illustrated in Video Supp.Video.2.

The editor UI provides a more accessible interface for inspecting and modifying existing models without directly editing the source code (Fig. Supp.Fig.2). Given that the main configuration file of Cellfoundry typically spans several thousand lines, direct navigation may be challenging for users unfamiliar with the codebase. The editor therefore offers a structured view of the model, enabling users to explore agent definitions and adjust parameters through a centralized interface. Both global environment properties and agent-dependent parameters can be modified, making it a suitable entry point when the goal is to use the base model while adjusting a limited set of parameters.

Together, these tools support a streamlined workflow in which models can be quickly prototyped using the configurator and subsequently explored and refined through the editor, while preserving full flexibility for direct code-level customization when required

#### C. Fibre Network Generation Implementation

Three-dimensional disordered fibre networks are generated using a custom Python implementation based on previously published approaches for collagen-like and biopolymer networks [21–23]. The implementation is also inspired by publicly available open-source routines for fibre network generation [39].

The present workflow builds on these earlier frameworks for mechanically relaxed fibre networks while adapting the generation pipeline to the agent-based representation of the extracellular matrix used in this work. In particular, the original MATLAB routines are translated into Python and extended with additional procedures for network optimization, boundary handling, and fibre subdivision. These extensions enable direct integration of the generated networks with the FNODE agents used in the simulations while preserving control over structural properties such as network density, connectivity, fibre length distribution, and segment resolution.

The following sections describe the main stages of the generation pipeline, including node placement, fibre connectivity construction, stochastic network relaxation, boundary processing, and fibre subdivision.

##### Initial node placement

A rectangular domain of size $L_x \times L_y \times L_z$ is defined, and the total number of initial nodes is prescribed through a target number density $\rho$ according to

$$N = \text{round}\left(\rho\, L_x L_y L_z\right).$$

A fraction of nodes is placed on selected domain faces in order to provide boundary anchoring, while the remaining nodes are sampled uniformly within the interior. In the present implementation, 5% of nodes per face are assigned to each pair of opposite boundaries. This procedure yields an initially isotropic and spatially homogeneous point cloud.

Each node is further assigned a target valency ($z_i \in \{2, 3, 4, 5, 6\}$), sampled from a prescribed probability distribution biased toward low connectivity, consistent with experimentally observed collagen and fibrin network architectures [21].

##### Initial fibre connectivity

An initial network connectivity is first constructed from the node positions by assigning fibres between nearby nodes. For each node $i$, candidate neighbours are identified within an expanding cubic search region until a sufficient number of nearby nodes is available. These candidates are then ordered by Euclidean distance, and fibres are created by randomly selecting neighbours while respecting the prescribed target valencies of both endpoints.

The resulting connectivity is subsequently post-processed to remove redundant connections. In particular, zero-length fibres are discarded, duplicate and mirrored edges are eliminated, and nodes with valency smaller than two are supplemented with additional nearest-neighbour connections where necessary. This yields an initial network with admissible connectivity and a well-defined set of fibre segments.

To quantify the geometric quality of this initial network, a quadratic fibre-length energy is defined as

$$E_{\text{length}} = \sum_{e=1}^{N_f} \left(\ell_e - \ell_0\right)^2,$$

where $N_f$ is the number of fibres, $\ell_e$ is the length of fibre $e$, and $\ell_0$ is the prescribed preferred fibre length. This energy serves as the objective for the first relaxation stage.

##### Fibre length optimization

The initial network is next relaxed by a stochastic optimization procedure aimed at reducing deviations of fibre lengths from the preferred value $\ell_0$. For each fibre connecting nodes $i$ and $j$, the local contribution to the fibre-length energy is

$$E_{\text{length},e} = \left(\|\mathbf{x}_i - \mathbf{x}_j\| - \ell_0\right)^2,$$

so that the total fibre-length energy is obtained by summing over all fibres.

Two classes of stochastic updates are considered. First, interior nodes are displaced by small random perturbations. For a proposed displacement, only the fibres incident to the selected node are modified, and the corresponding local contribution to $E_{\text{length}}$ is re-evaluated. The move is accepted whenever this local energy decreases. In addition, slightly unfavorable moves may also be accepted with small probability, which introduces a simulated-annealing-like mechanism that helps the optimization avoid trapping in shallow local minima.

Second, local edge-swapping operations are attempted. In these moves, pairs of neighbouring fibres are rewired to form an alternative local connectivity pattern, and the fibre-length energy of the modified pair is compared with that of the original configuration. A swap is accepted only if it reduces the associated local fibre-length energy.

The characteristic amplitude of the random displacement proposals is reduced progressively over the course of the optimization, so that early iterations explore the configuration space more broadly while later iterations become increasingly conservative. If geometric constraints are enabled, proposed displacement moves can additionally be required to remain inside the computational domain by either rejecting out-of-bounds proposals or clipping them back to the admissible region.

#### Branch alignment optimization

After fibre-length relaxation, a second optimization stage is applied to improve the local branching geometry. For each node $i$, unit direction vectors are computed for all connected fibres as

$$\mathbf{t}_k = \frac{\mathbf{x}_{j_k} - \mathbf{x}_i}{\|\mathbf{x}_{j_k} - \mathbf{x}_i\|},$$

where $\mathbf{x}_i$ is the position of node $i$ and $\mathbf{x}_{j_k}$ is the position of the neighbouring node connected by fibre $k$. All pairwise dot products between these directions are then evaluated, and the pair with minimum dot product is identified as the most opposing pair of branches.

Based on this local geometry, a nodal branching energy is defined as

$$E_{\text{branch},i} = w\left(1 + \min_{a<b} \mathbf{t}_a \cdot \mathbf{t}_b\right) + \left[(n_i - 2) - \max\left(\sum_{k\in R_i} \mathbf{t}_k \cdot \mathbf{t}_{a^*}, \sum_{k\in R_i} \mathbf{t}_k \cdot \mathbf{t}_{b^*}\right)\right],$$

where $n_i$ is the valency of node $i$, $(a^*, b^*)$ denotes the pair of fibres with minimum dot product, and $R_i$ is the set of remaining fibres after this pair has been removed. The first term penalizes configurations in which no pair of fibres is close to antiparallel, while the second penalizes poor alignment of the remaining fibres with whichever member of the selected opposing pair provides the stronger overall alignment. The coefficient $w$ is an empirical weighting factor controlling the relative importance of these two contributions.

Branch alignment is optimized through stochastic local displacements involving the selected node together with its directly connected neighbours. For each proposed move, the summed branching energy over this local neighbourhood is recomputed, and the move is accepted whenever this local branching energy decreases. As in the fibre-length optimization stage, a small fraction of mildly unfavorable moves may also be accepted probabilistically in order to reduce trapping in poor local minima. Optional rejection-based or clipping-based boundary handling can again be used to keep proposed node positions inside the computational domain.

#### Boundary snapping and connectivity pruning

To impose geometric constraints at the boundaries, nodes located within a prescribed distance of selected faces can optionally be snapped exactly onto the corresponding planes. Following this operation, edges connecting pairs of nodes lying on the same boundary plane are removed in order to avoid artificial in-plane loops. Nodes that become coincident within a specified tolerance are merged, and the corresponding connectivity graph is updated accordingly.

#### Fibre subdivision for agent-based modelling

To improve spatial resolution in the subsequent FNODE agent-based simulations, long fibres are subdivided into shorter segments of maximum length $\ell_{\text{edge}}$.

For a fibre connecting nodes $i$ and $j$ with length $\ell_{ij}$, subdivision is triggered whenever

$$\ell_{ij} > \ell_{\text{edge}}.$$

The number of intermediate nodes introduced along that fibre is

$$n_{\text{new}} = \left\lceil \frac{\ell_{ij}}{\ell_{\text{edge}}} \right\rceil - 1.$$

These intermediate nodes are placed uniformly along the original segment according to

$$\mathbf{x}_i^{(k)} = \mathbf{x}_i + \frac{k}{n_{\text{new}} + 1} \left( \mathbf{x}_j - \mathbf{x}_i \right), \qquad k = 1, \dots, n_{\text{new}}.$$

The original edge is then replaced by a chain of shorter edges connecting the original endpoints through the inserted nodes.

#### Export and diagnostics

The final network is stored as node coordinates and adjacency lists and exported in VTK format for visualization and inspection. Standard network statistics, including nodal valency distributions, fibre-length distributions, and pore-size histograms, are computed for quality control.

Overall, this generation and refinement pipeline yields statistically isotropic, mechanically relaxed fibre networks with controllable density, connectivity, fibre-length distribution, and segment resolution.

#### D. Pre-defined Optimization Targets

In addition to the generic time-series and scalar endpoint objectives described in Section 3, the framework provides a set of pre-defined targets tailored to common calibration tasks. These objectives are designed to facilitate direct comparison between simulation outputs and experimental measurements across multiple scales, including mechanical characterization, cell migration behaviour, and organoid-level descriptors. The available targets include formulations for fitting stress–strain responses and differential moduli, matching population-level cell motility statistics, and reproducing geometric measures such as final organoid size. Each objective follows a consistent interface and can be directly selected within the optimization configuration. The following subsections summarize the main pre-defined optimization targets currently implemented in the framework.

*Stress–strain curve fitting*

For mechanical calibration, engineering strain $\varepsilon$ and a boundary-derived stress proxy $\sigma$ are extracted from the simulated boundary displacement and force signals, respectively. Depending on the assay, both normal and shear loading modes are supported. The stress signal may be normalized either by the geometric boundary area or by the total cross-sectional area of attached fibres, enabling comparison against different experimental stress definitions.

The stress–strain objective combines discrepancies in both stress and strain through a weighted sum of mean-squared errors:

$$f_{\sigma\varepsilon} = \mathrm{MSE}\big(\hat{\boldsymbol{\sigma}}, \boldsymbol{\sigma}^{\mathrm{ref}}\big) + w\,\mathrm{MSE}\big(\hat{\boldsymbol{\varepsilon}}, \boldsymbol{\varepsilon}^{\mathrm{ref}}\big)$$

where $\hat{\boldsymbol{\sigma}}$ and $\hat{\boldsymbol{\varepsilon}}$ denote the simulated stress and strain curves, and $\boldsymbol{\sigma}^{\mathrm{ref}}$, $\boldsymbol{\varepsilon}^{\mathrm{ref}}$ are the corresponding reference data. The weighting parameter $w$ controls the relative importance of matching stress versus strain. In practice, the strain term acts mainly as a coverage penalty, ensuring that the simulated loading path spans the reference strain range.

*Differential-modulus fitting*

To calibrate nonlinear matrix mechanics more directly, the framework also supports fitting the differential modulus $K(\varepsilon) = d\sigma/d\varepsilon$ as a function of strain. The simulated stress–strain curve is first differentiated numerically, optionally smoothed, and then interpolated onto the reference strain grid. The resulting error is computed as a mean-squared difference between simulated and reference differential-modulus curves, with an optional penalty for insufficient strain-range coverage. As with stress–strain fitting, both normal and shear variants are available.

*Cell-speed objectives*

To capture cell motility phenotypes, the framework includes objectives based on per-cell trajectory statistics aggregated at the population level. For each cell, two complementary velocity measures are computed from its tracked trajectory:

- Mean speed $v_{\mathrm{mean}}$, defined as the cumulative path length divided by the trajectory duration.
- Effective speed $v_{\mathrm{eff}}$, defined as the net displacement divided by the trajectory duration.

These quantities distinguish between persistent and random motion, allowing the model to capture both directional migration and diffusive-like behaviour.

For each cell type, the selected population statistic (mean or median) is compared against reference targets. The error is defined as a sum (or average) of relative absolute errors,

$$f_{\mathrm{speed}} = \frac{1}{N_t}\sum_{k=1}^{N_t} \frac{|v_k - v_k^{\mathrm{ref}}|}{|v_k^{\mathrm{ref}}|},$$

where $v_k$ represents either $v_{\mathrm{mean}}$ or $v_{\mathrm{eff}}$ for a given cell type or metric, and $N_t$ is the number of target quantities. Optional filtering can be applied based on trajectory duration or cell viability, enabling robust comparison with experimental tracking data.

*Organoid size*

Spheroid or organoid size is quantified from the three-dimensional point cloud of alive-cell centroids extracted from the final simulation snapshot. Several geometric descriptors are available. The default metric is the radius of gyration,

$$R_g = \sqrt{\frac{1}{N}\sum_{i=1}^{N} \|\mathbf{r}_i - \bar{\mathbf{r}}\|^2},$$

where $\mathbf{r}_i$ is the position of the $i$-th alive cell and $\bar{\mathbf{r}}$ is the centroid of the alive-cell cloud. The equivalent uniform-sphere radius $R_{\mathrm{eq}} = R_g\sqrt{5/3}$ and the maximum inter-cell Euclidean distance are also available as alternative size metrics.

In addition, a dimensionless sphericity measure is computed from the covariance matrix of the centred positions. Let $\lambda_{\min}$ and $\lambda_{\max}$ denote the minimum and maximum eigenvalues of $\mathrm{cov}(\mathbf{r}_i - \bar{\mathbf{r}})$, respectively. The sphericity is defined as

$$\Psi = \frac{\lambda_{\min}}{\lambda_{\max}},$$

with $\Psi \in [0, 1]$, where $\Psi = 1$ corresponds to an isotropic distribution. For $N < 4$, $\Psi$ is set to 1. These objectives require final VTK output so that cell-centre positions can be reconstructed from the saved geometry.

### E. Benchmark Data

This appendix includes additional figures to better visualize the benchmark results, describes the multivariate model used to support the discussion in the main text, and includes the raw timing data obtained from the simulations.

#### Per GPU-results

To further characterise platform-specific performance, pairwise response surfaces were constructed using representative combinations of benchmark parameters (Fig. Supp.Fig.3). These plots highlight how initialization and runtime depend on model composition, revealing notably stable per-step performance across large regions of the parameter space for the A100 and H100 NVL.

#### Multivariate Model

A multivariate model is formulated in log–log form to stablish a relationship between agent population and model initialization or step times. Let $y$ denote either the initialization time or the mean simulation step time. The independent benchmark inputs are defined as

$$x_1 = N_{\mathrm{ECM}}, \qquad x_2 = N_{\mathrm{CELL}}, \qquad x_3 = N_{\mathrm{FNODES}}, \qquad x_4 = R_{\mathrm{cell}}.$$

Because the number of focal adhesions per cell is fixed in the benchmark configurations, the focal adhesion population is derived from the cell count and therefore is not included as an independent regressor.

Each input variable is transformed to logarithmic scale and standardized according to

$$z_i = \frac{\log_{10}(x_i) - \mu_i}{\sigma_i},$$

where $\mu_i$ and $\sigma_i$ denote the sample mean and standard deviation of $\log_{10}(x_i)$ over the fitted benchmark dataset.

The regression model includes both main effects and pairwise interactions:

$$\log_{10}(y) = \beta_0 + \sum_{i=1}^{4} \beta_i z_i + \sum_{1 \le i < j \le 4} \beta_{ij} z_i z_j + \varepsilon.$$

A positive main coefficient $\beta_i$ indicates that increasing the corresponding benchmark variable tends to increase runtime after accounting for the other variables. A negative interaction coefficient $\beta_{ij}$ indicates that the joint effect of increasing both variables is smaller than the sum of their individual contributions in the standardized log-transformed space. For example, a negative ECM–FNODE interaction implies that simultaneously increasing ECM and FNODE counts produces a sub-additive effect relative to the individual contributions of each variable (Fig. Supp.Fig.3).

#### Raw Data

This section presents the raw benchmarking results obtained across the different GPU architectures evaluated. For each device, detailed tables report the full set of simulation parameters alongside the measured execution times, including initialization and runtime per step. To ensure a fair comparison, compilation overhead ($t_{\mathrm{RTC}}$) has been excluded from the reported initialization time, isolating the true cost of model setup. A representative rendering of one of the most demanding benchmark configurations is shown in Fig. Supp.Fig.5, illustrating the extreme agent density reached in these tests and the severity of the computational workload imposed on the model.

(tables 4–5–6–7).

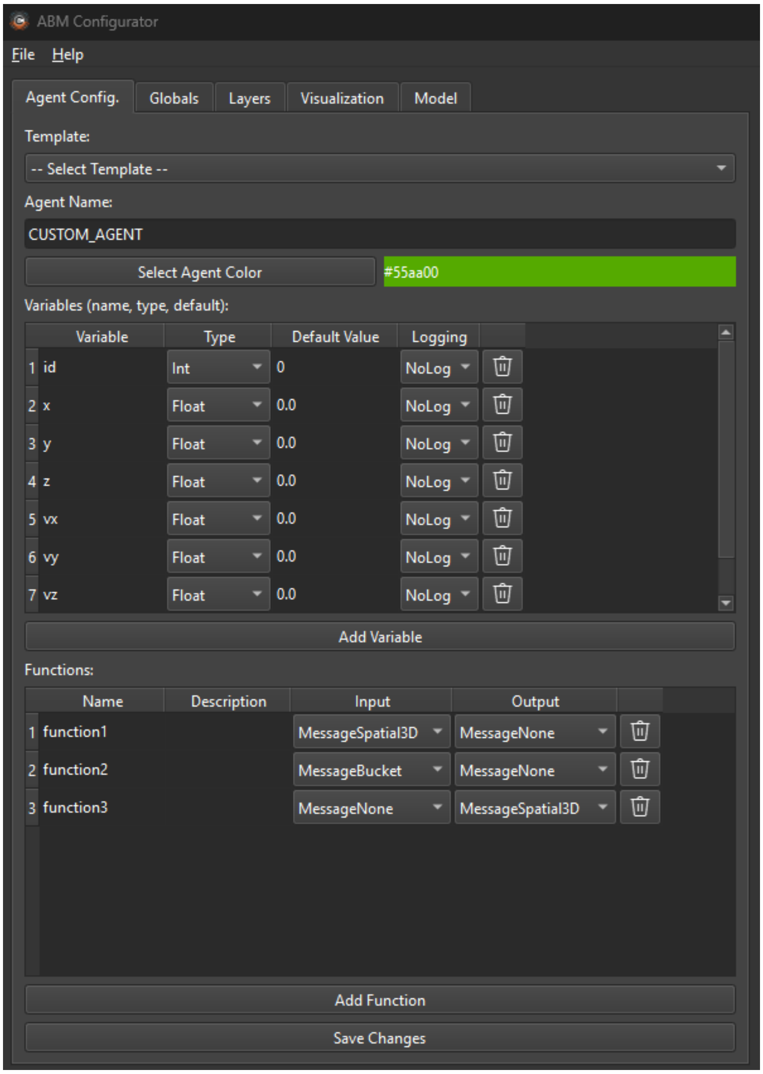


**Figure Supp.Fig.1: Overview of the agent configurator and other tabs.** The configurator tab allows users to define agent types, variables, and functions, and to visually link agents to represent interactions. The generated model structure is automatically created based on the defined configuration.

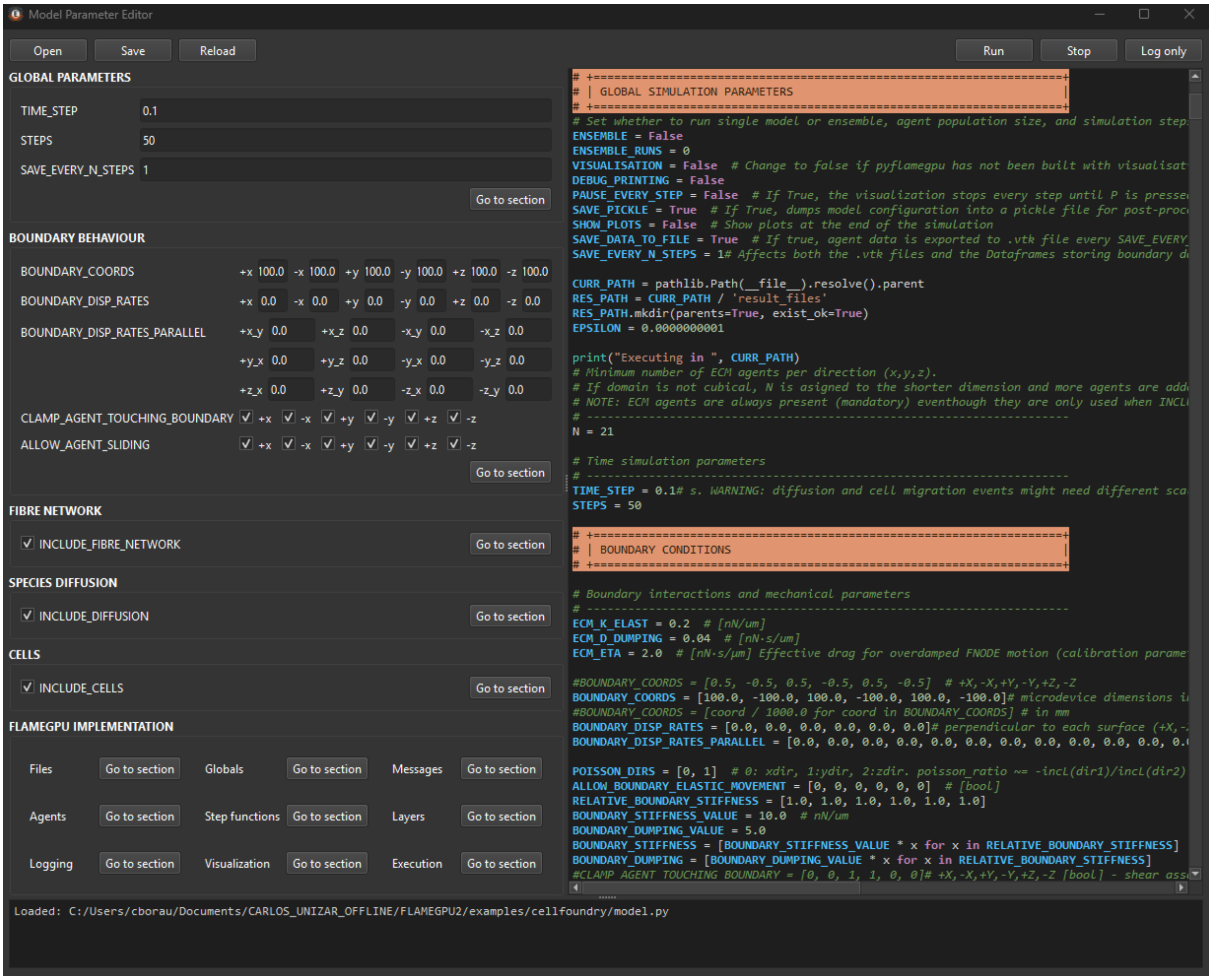


**Figure Supp.Fig.2: Overview of the agent editor.** The editor provides a structured view of the model, enabling users to explore agent definitions and adjust parameters through a centralized interface.

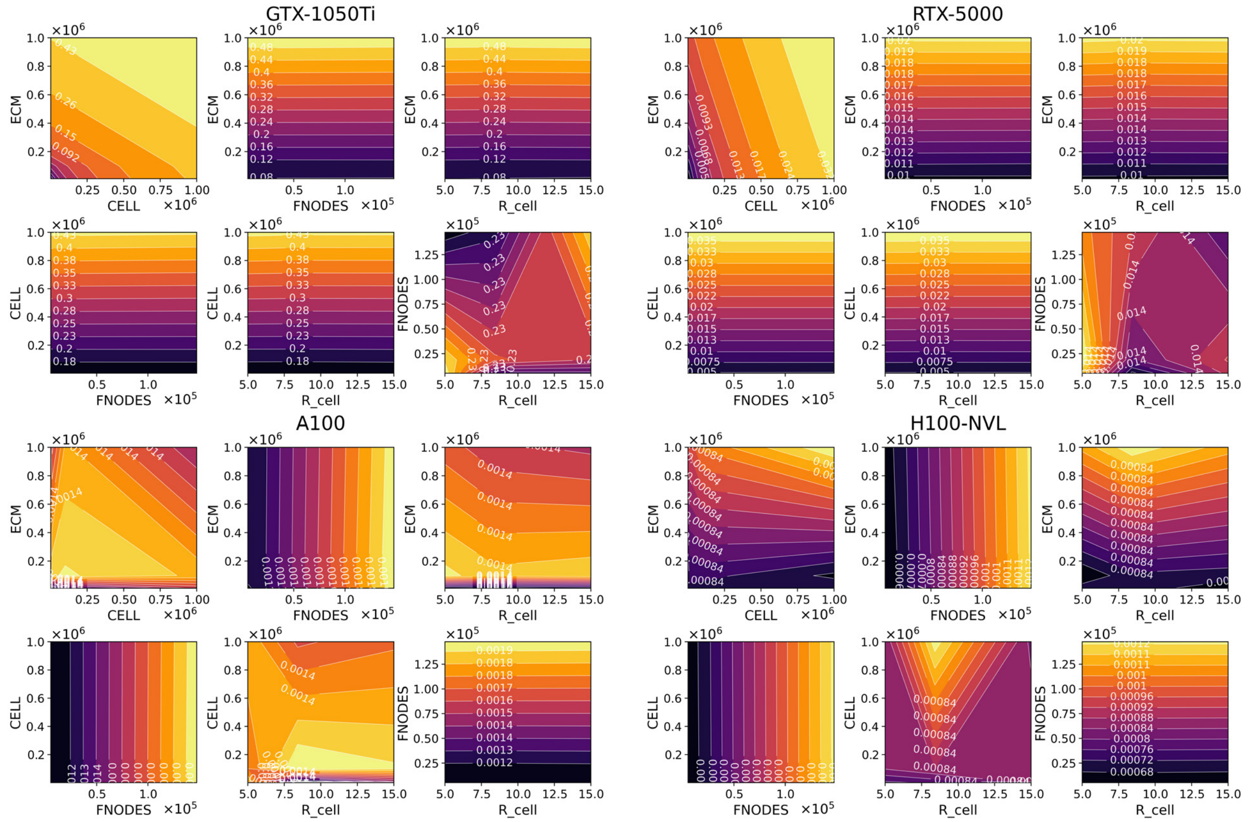


**Figure Supp.Fig.3: Per GPU-results.** For each platform, the upper row shows initialization time and the lower row shows mean runtime per step, plotted for representative pairs of benchmark parameters. Overall, runtime depends on model composition beyond total agent count alone. Notably, per-step times on the A100 and H100 NVL remain nearly flat across large portions of the explored parameter space.

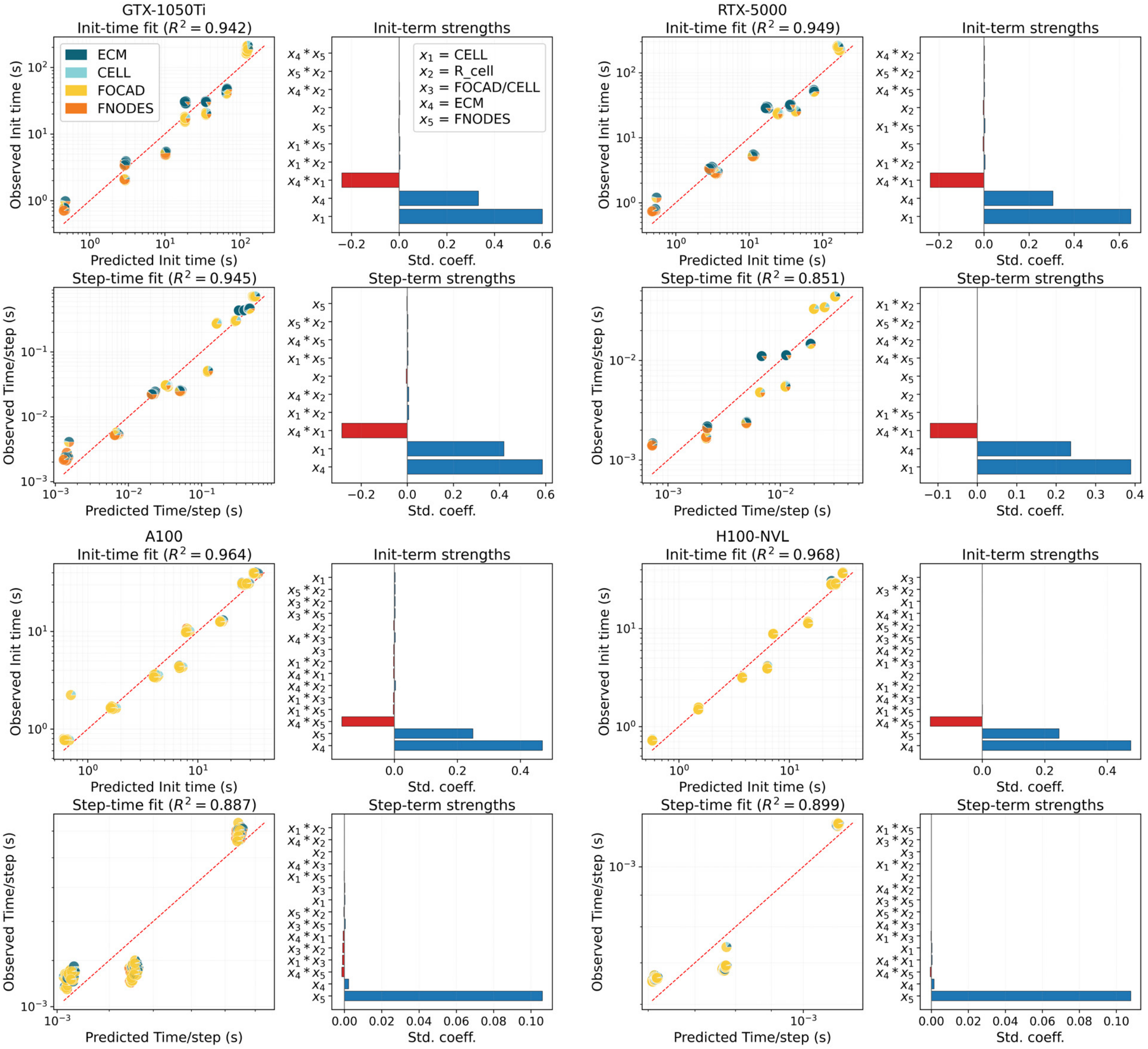


**Figure Supp.Fig.4: Multivariate log–log regression analysis of initialization time and mean runtime per step for each GPU architecture.** For each platform, the left panels compare observed and predicted values, with the dashed red line indicating perfect agreement; the coefficient of determination is reported in each title. The right panels show standardized regression coefficients for the retained terms, where $x_1 = N_{\text{CELL}}$, $x_2 = R_{\text{cell}}$, $x_3 = N^{\text{init}}_{\text{FOCAD}}$, $x_4 = N_{\text{ECM}}$, and $x_5 = N_{\text{FNODE}}$. These fits quantify how model composition, beyond total agent count alone, controls both initialization and stepping cost.

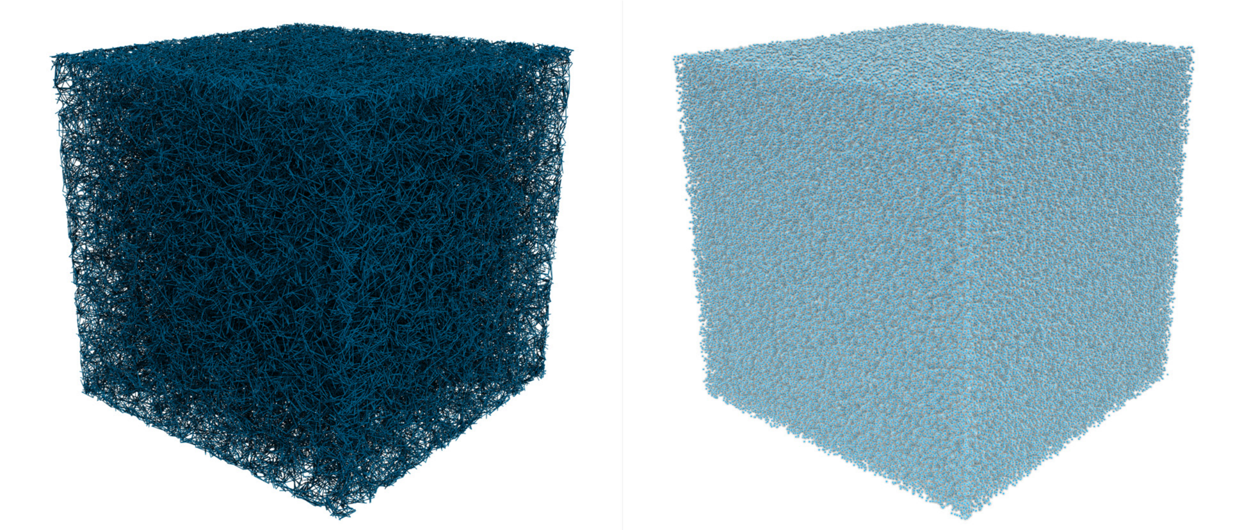

**Figure Supp.Fig.5: 3D rendering of a high-density benchmarking configuration showing the densest fibre network and cell population considered in this study.** The snapshot illustrates the extreme packing achieved within a $1\ \mathrm{mm}^3$ simulation domain, containing $10^6$ `CELL` agents (associated to $10^7$ `FOCAD` agents, not rendered) and 147,599 `FNODE` agents. It is intended to convey how strongly the framework is being stressed under these conditions, both in terms of agent count and the resulting communication workload required to resolve their interactions.

Table 4: GTX5000Ti raw benchmark results. Times are in seconds. Compilation time ($t_{RTC}$) is subtracted from $t_{init}$ to compute actual initialization time.

| Run | $N$ | ECM | CELL | FOCAD/CELL | FOCAD | FNODE | $R_c$ | $R_s$ | Steps | $t_{init}$ | $t_{sim}$ | $t_{RTC}$ | $t_{initfn}$ | $t_{exit}$ | $t_{total}$ | $t_{step}$ |
|---|---|---|---|---|---|---|---|---|---|---|---|---|---|---|---|---|
| 1 | 21 | 9261 | 1000 | 5 | 5000 | 5249 | 5.0 | 15.0 | 100 | 55.100 | 0.408 | 54.119 | 0.404 | 0.000 | 56.051 | 0.004 |
| 2 | 21 | 9261 | 1000 | 5 | 5000 | 18671 | 5.0 | 15.0 | 100 | 1.018 | 0.237 | 0.296 | 0.384 | 0.000 | 1.818 | 0.002 |
| 3 | 21 | 9261 | 1000 | 5 | 5000 | 147599 | 5.0 | 15.0 | 100 | 0.781 | 0.278 | 0.073 | 0.378 | 0.000 | 1.582 | 0.003 |
| 4 | 21 | 9261 | 1000 | 5 | 5000 | 5249 | 8.4 | 25.2 | 100 | 0.814 | 0.236 | 0.095 | 0.383 | 0.000 | 1.629 | 0.002 |
| 5 | 21 | 9261 | 1000 | 5 | 5000 | 18671 | 8.4 | 25.2 | 100 | 0.834 | 0.212 | 0.083 | 0.397 | 0.000 | 1.572 | 0.002 |
| 6 | 21 | 9261 | 1000 | 5 | 5000 | 147599 | 8.4 | 25.2 | 100 | 0.820 | 0.228 | 0.072 | 0.413 | 0.000 | 2.366 | 0.002 |
| 7 | 21 | 9261 | 1000 | 5 | 5000 | 5249 | 15.0 | 45.0 | 100 | 0.828 | 0.236 | 0.072 | 0.384 | 0.000 | 1.636 | 0.002 |
| 8 | 21 | 9261 | 1000 | 5 | 5000 | 18671 | 15.0 | 45.0 | 100 | 0.872 | 0.226 | 0.088 | 0.398 | 0.000 | 1.684 | 0.002 |
| 9 | 21 | 9261 | 1000 | 5 | 5000 | 147599 | 15.0 | 45.0 | 100 | 0.782 | 0.217 | 0.073 | 0.383 | 0.000 | 1.537 | 0.002 |
| 10 | 21 | 9261 | 10000 | 5 | 50000 | 5249 | 5.0 | 15.0 | 100 | 2.082 | 0.535 | 0.072 | 1.695 | 0.000 | 3.238 | 0.005 |
| 11 | 21 | 9261 | 10000 | 5 | 50000 | 18671 | 5.0 | 15.0 | 100 | 2.139 | 0.542 | 0.072 | 1.751 | 0.000 | 3.158 | 0.005 |
| 12 | 21 | 9261 | 10000 | 5 | 50000 | 147599 | 5.0 | 15.0 | 100 | 2.161 | 0.524 | 0.072 | 1.707 | 0.000 | 3.200 | 0.005 |
| 13 | 21 | 9261 | 10000 | 5 | 50000 | 5249 | 8.4 | 25.2 | 100 | 2.179 | 0.534 | 0.071 | 1.763 | 0.000 | 3.176 | 0.005 |
| 14 | 21 | 9261 | 10000 | 5 | 50000 | 18671 | 8.4 | 25.2 | 100 | 2.155 | 0.573 | 0.072 | 1.749 | 0.000 | 3.243 | 0.006 |
| 15 | 21 | 9261 | 10000 | 5 | 50000 | 147599 | 8.4 | 25.2 | 100 | 2.132 | 0.536 | 0.071 | 1.728 | 0.000 | 3.232 | 0.005 |
| 16 | 21 | 9261 | 10000 | 5 | 50000 | 5249 | 15.0 | 45.0 | 100 | 2.097 | 0.525 | 0.071 | 1.698 | 0.000 | 3.170 | 0.005 |
| 17 | 21 | 9261 | 10000 | 5 | 50000 | 18671 | 15.0 | 45.0 | 100 | 2.159 | 0.529 | 0.072 | 1.711 | 0.000 | 3.245 | 0.005 |
| 18 | 21 | 9261 | 10000 | 5 | 50000 | 147599 | 15.0 | 45.0 | 100 | 2.164 | 0.520 | 0.071 | 1.736 | 0.000 | 3.172 | 0.005 |
| 19 | 21 | 9261 | 100000 | 5 | 500000 | 5249 | 5.0 | 15.0 | 100 | 15.310 | 2.909 | 0.073 | 14.858 | 0.000 | 18.750 | 0.029 |
| 20 | 21 | 9261 | 100000 | 5 | 500000 | 18671 | 5.0 | 15.0 | 100 | 15.898 | 3.080 | 0.072 | 15.503 | 0.000 | 19.466 | 0.031 |
| 21 | 21 | 9261 | 100000 | 5 | 500000 | 147599 | 5.0 | 15.0 | 100 | 18.229 | 3.075 | 0.073 | 17.813 | 0.000 | 21.859 | 0.031 |
| 22 | 21 | 9261 | 100000 | 5 | 500000 | 5249 | 8.4 | 25.2 | 100 | 17.769 | 3.072 | 0.073 | 17.343 | 0.000 | 21.372 | 0.031 |
| 23 | 21 | 9261 | 100000 | 5 | 500000 | 18671 | 8.4 | 25.2 | 100 | 17.182 | 3.070 | 0.072 | 16.778 | 0.000 | 20.781 | 0.031 |
| 24 | 21 | 9261 | 100000 | 5 | 500000 | 147599 | 8.4 | 25.2 | 100 | 17.543 | 3.105 | 0.073 | 17.144 | 0.000 | 21.989 | 0.031 |
| 25 | 21 | 9261 | 100000 | 5 | 500000 | 5249 | 15.0 | 45.0 | 100 | 17.001 | 3.091 | 0.072 | 16.592 | 0.000 | 20.603 | 0.031 |
| 26 | 21 | 9261 | 100000 | 5 | 500000 | 18671 | 15.0 | 45.0 | 100 | 17.750 | 3.079 | 0.074 | 17.345 | 0.000 | 21.384 | 0.031 |
| 27 | 21 | 9261 | 100000 | 5 | 500000 | 147599 | 15.0 | 45.0 | 100 | 17.310 | 3.065 | 0.090 | 16.841 | 0.000 | 20.898 | 0.031 |
| 28 | 21 | 9261 | 1000000 | 5 | 5000000 | 5249 | 5.0 | 15.0 | 100 | 162.796 | 28.078 | 0.074 | 162.358 | 0.000 | 191.418 | 0.281 |
| 29 | 21 | 9261 | 1000000 | 5 | 5000000 | 18671 | 5.0 | 15.0 | 100 | 164.064 | 28.046 | 0.072 | 163.624 | 0.000 | 192.690 | 0.280 |
| 30 | 21 | 9261 | 1000000 | 5 | 5000000 | 147599 | 5.0 | 15.0 | 100 | 163.524 | 28.082 | 0.073 | 163.102 | 0.000 | 192.158 | 0.281 |
| 31 | 21 | 9261 | 1000000 | 5 | 5000000 | 5249 | 8.4 | 25.2 | 100 | 164.382 | 28.039 | 0.073 | 163.988 | 0.000 | 193.014 | 0.280 |
| 32 | 21 | 9261 | 1000000 | 5 | 5000000 | 18671 | 8.4 | 25.2 | 100 | 163.886 | 28.089 | 0.073 | 163.452 | 0.000 | 192.559 | 0.281 |
| 33 | 21 | 9261 | 1000000 | 5 | 5000000 | 147599 | 8.4 | 25.2 | 100 | 162.360 | 27.411 | 0.072 | 161.921 | 0.000 | 190.357 | 0.274 |
| 34 | 21 | 9261 | 1000000 | 5 | 5000000 | 5249 | 15.0 | 45.0 | 100 | 163.534 | 27.226 | 0.073 | 163.091 | 0.000 | 191.305 | 0.272 |
| 35 | 21 | 9261 | 1000000 | 5 | 5000000 | 18671 | 15.0 | 45.0 | 100 | 163.913 | 27.216 | 0.071 | 163.495 | 0.000 | 191.723 | 0.272 |
| 36 | 21 | 9261 | 1000000 | 5 | 5000000 | 147599 | 15.0 | 45.0 | 100 | 164.344 | 27.272 | 0.072 | 163.904 | 0.000 | 192.987 | 0.273 |
| 37 | 46 | 97336 | 1000 | 5 | 5000 | 5249 | 5.0 | 15.0 | 100 | 62.248 | 2.441 | 58.757 | 3.099 | 0.000 | 65.301 | 0.024 |
| 38 | 46 | 97336 | 1000 | 5 | 5000 | 18671 | 5.0 | 15.0 | 100 | 4.185 | 2.331 | 0.360 | 3.525 | 0.000 | 7.868 | 0.023 |
| 39 | 46 | 97336 | 1000 | 5 | 5000 | 147599 | 5.0 | 15.0 | 100 | 3.608 | 2.282 | 0.073 | 3.215 | 0.000 | 6.421 | 0.023 |
| 40 | 46 | 97336 | 1000 | 5 | 5000 | 5249 | 8.4 | 25.2 | 100 | 3.985 | 2.216 | 0.072 | 3.529 | 0.000 | 7.517 | 0.022 |
| 41 | 46 | 97336 | 1000 | 5 | 5000 | 18671 | 8.4 | 25.2 | 100 | 3.866 | 2.202 | 0.072 | 3.429 | 0.000 | 7.396 | 0.022 |
| 42 | 46 | 97336 | 1000 | 5 | 5000 | 147599 | 8.4 | 25.2 | 100 | 3.796 | 2.213 | 0.073 | 3.337 | 0.000 | 6.533 | 0.022 |
| 43 | 46 | 97336 | 1000 | 5 | 5000 | 5249 | 15.0 | 45.0 | 100 | 3.616 | 2.219 | 0.073 | 3.243 | 0.000 | 6.351 | 0.022 |
| 44 | 46 | 97336 | 1000 | 5 | 5000 | 18671 | 15.0 | 45.0 | 100 | 3.566 | 2.239 | 0.072 | 3.178 | 0.000 | 7.097 | 0.022 |
| 45 | 46 | 97336 | 1000 | 5 | 5000 | 147599 | 15.0 | 45.0 | 100 | 3.447 | 2.218 | 0.073 | 3.040 | 0.000 | 6.184 | 0.022 |
| 46 | 46 | 97336 | 10000 | 5 | 50000 | 5249 | 5.0 | 15.0 | 100 | 5.125 | 2.520 | 0.071 | 4.731 | 0.000 | 8.175 | 0.025 |
| 47 | 46 | 97336 | 10000 | 5 | 50000 | 18671 | 5.0 | 15.0 | 100 | 5.154 | 2.516 | 0.078 | 4.744 | 0.000 | 8.199 | 0.025 |
| 48 | 46 | 97336 | 10000 | 5 | 50000 | 147599 | 5.0 | 15.0 | 100 | 4.897 | 2.511 | 0.073 | 4.495 | 0.000 | 7.886 | 0.025 |
| 49 | 46 | 97336 | 10000 | 5 | 50000 | 5249 | 8.4 | 25.2 | 100 | 4.925 | 2.457 | 0.072 | 4.492 | 0.000 | 7.969 | 0.025 |
| 50 | 46 | 97336 | 10000 | 5 | 50000 | 18671 | 8.4 | 25.2 | 100 | 5.074 | 2.500 | 0.073 | 4.651 | 0.000 | 8.119 | 0.025 |
| 51 | 46 | 97336 | 10000 | 5 | 50000 | 147599 | 8.4 | 25.2 | 100 | 5.050 | 2.574 | 0.089 | 4.597 | 0.000 | 8.157 | 0.026 |
| 52 | 46 | 97336 | 10000 | 5 | 50000 | 5249 | 15.0 | 45.0 | 100 | 5.534 | 2.556 | 0.089 | 5.128 | 0.000 | 8.664 | 0.026 |

*Continued on next page*

(continued from previous page)

| Run | $N$ | ECM | CELL | FOCAD/CELL | FOCAD | FNODE | $R_c$ | $R_s$ | Steps | $t_{init}$ | $t_{sim}$ | $t_{RTC}$ | $t_{initfn}$ | $t_{exit}$ | $t_{total}$ | $t_{step}$ |
|---|---|---|---|---|---|---|---|---|---|---|---|---|---|---|---|---|
| 53 | 46 | 97336 | 10000 | 5 | 50000 | 18671 | 15.0 | 45.0 | 100 | 5.430 | 2.511 | 0.093 | 4.958 | 0.000 | 9.331 | 0.025 |
| 54 | 46 | 97336 | 10000 | 5 | 50000 | 147599 | 15.0 | 45.0 | 100 | 5.250 | 2.487 | 0.073 | 4.829 | 0.000 | 8.286 | 0.025 |
| 55 | 46 | 97336 | 100000 | 5 | 500000 | 5249 | 5.0 | 15.0 | 100 | 19.401 | 4.952 | 0.072 | 18.934 | 0.000 | 25.683 | 0.050 |
| 56 | 46 | 97336 | 100000 | 5 | 500000 | 18671 | 5.0 | 15.0 | 100 | 19.326 | 4.937 | 0.074 | 18.900 | 0.000 | 24.754 | 0.049 |
| 57 | 46 | 97336 | 100000 | 5 | 500000 | 147599 | 5.0 | 15.0 | 100 | 19.641 | 4.947 | 0.072 | 19.196 | 0.000 | 25.139 | 0.049 |
| 58 | 46 | 97336 | 100000 | 5 | 500000 | 5249 | 8.4 | 25.2 | 100 | 19.423 | 5.037 | 0.072 | 18.984 | 0.000 | 25.797 | 0.050 |
| 59 | 46 | 97336 | 100000 | 5 | 500000 | 18671 | 8.4 | 25.2 | 100 | 20.973 | 5.077 | 0.076 | 20.558 | 0.000 | 26.586 | 0.051 |
| 60 | 46 | 97336 | 100000 | 5 | 500000 | 147599 | 8.4 | 25.2 | 100 | 19.931 | 5.030 | 0.092 | 19.492 | 0.000 | 25.495 | 0.050 |
| 61 | 46 | 97336 | 100000 | 5 | 500000 | 5249 | 15.0 | 45.0 | 100 | 20.015 | 4.951 | 0.073 | 19.591 | 0.000 | 25.554 | 0.050 |
| 62 | 46 | 97336 | 100000 | 5 | 500000 | 18671 | 15.0 | 45.0 | 100 | 19.903 | 5.122 | 0.074 | 19.478 | 0.000 | 26.382 | 0.051 |
| 63 | 46 | 97336 | 100000 | 5 | 500000 | 147599 | 15.0 | 45.0 | 100 | 20.108 | 5.107 | 0.074 | 19.684 | 0.000 | 25.772 | 0.051 |
| 64 | 46 | 97336 | 1000000 | 5 | 5000000 | 5249 | 5.0 | 15.0 | 100 | 170.557 | 30.794 | 0.073 | 170.135 | 0.000 | 201.976 | 0.308 |
| 65 | 46 | 97336 | 1000000 | 5 | 5000000 | 18671 | 5.0 | 15.0 | 100 | 166.695 | 30.025 | 0.074 | 166.271 | 0.000 | 197.325 | 0.300 |
| 66 | 46 | 97336 | 1000000 | 5 | 5000000 | 147599 | 5.0 | 15.0 | 100 | 165.058 | 30.021 | 0.072 | 164.673 | 0.000 | 196.453 | 0.300 |
| 67 | 46 | 97336 | 1000000 | 5 | 5000000 | 5249 | 8.4 | 25.2 | 100 | 163.432 | 29.840 | 0.074 | 163.026 | 0.000 | 193.789 | 0.298 |
| 68 | 46 | 97336 | 1000000 | 5 | 5000000 | 18671 | 8.4 | 25.2 | 100 | 164.764 | 29.555 | 0.090 | 164.303 | 0.000 | 195.193 | 0.296 |
| 69 | 46 | 97336 | 1000000 | 5 | 5000000 | 147599 | 8.4 | 25.2 | 100 | 161.479 | 30.319 | 0.074 | 161.019 | 0.000 | 192.408 | 0.303 |
| 70 | 46 | 97336 | 1000000 | 5 | 5000000 | 5249 | 15.0 | 45.0 | 100 | 169.008 | 29.683 | 0.088 | 168.553 | 0.000 | 199.304 | 0.297 |
| 71 | 46 | 97336 | 1000000 | 5 | 5000000 | 18671 | 15.0 | 45.0 | 100 | 170.328 | 30.466 | 0.072 | 169.907 | 0.000 | 201.377 | 0.305 |
| 72 | 46 | 97336 | 1000000 | 5 | 5000000 | 147599 | 15.0 | 45.0 | 100 | 171.703 | 30.360 | 0.074 | 171.313 | 0.000 | 202.666 | 0.304 |
| 73 | 100 | 1000000 | 1000 | 5 | 5000 | 5249 | 5.0 | 15.0 | 100 | 83.174 | 43.307 | 54.639 | 28.169 | 0.000 | 127.837 | 0.433 |
| 74 | 100 | 1000000 | 1000 | 5 | 5000 | 18671 | 5.0 | 15.0 | 100 | 30.340 | 43.139 | 0.297 | 29.701 | 0.000 | 74.081 | 0.431 |
| 75 | 100 | 1000000 | 1000 | 5 | 5000 | 147599 | 5.0 | 15.0 | 100 | 29.649 | 43.002 | 0.072 | 29.242 | 0.000 | 73.232 | 0.430 |
| 76 | 100 | 1000000 | 1000 | 5 | 5000 | 5249 | 8.4 | 25.2 | 100 | 30.696 | 43.028 | 0.074 | 30.283 | 0.000 | 74.313 | 0.430 |
| 77 | 100 | 1000000 | 1000 | 5 | 5000 | 18671 | 8.4 | 25.2 | 100 | 31.137 | 43.476 | 0.074 | 30.679 | 0.000 | 75.163 | 0.435 |
| 78 | 100 | 1000000 | 1000 | 5 | 5000 | 147599 | 8.4 | 25.2 | 100 | 30.271 | 43.227 | 0.075 | 29.877 | 0.000 | 74.121 | 0.432 |
| 79 | 100 | 1000000 | 1000 | 5 | 5000 | 5249 | 15.0 | 45.0 | 100 | 29.438 | 43.303 | 0.073 | 28.988 | 0.000 | 73.331 | 0.433 |
| 80 | 100 | 1000000 | 1000 | 5 | 5000 | 18671 | 15.0 | 45.0 | 100 | 28.862 | 43.642 | 0.072 | 28.440 | 0.000 | 73.078 | 0.436 |
| 81 | 100 | 1000000 | 1000 | 5 | 5000 | 147599 | 15.0 | 45.0 | 100 | 30.564 | 43.656 | 0.072 | 30.177 | 0.000 | 74.905 | 0.437 |
| 82 | 100 | 1000000 | 10000 | 5 | 50000 | 5249 | 5.0 | 15.0 | 100 | 31.373 | 43.758 | 0.074 | 30.966 | 0.000 | 75.734 | 0.438 |
| 83 | 100 | 1000000 | 10000 | 5 | 50000 | 18671 | 5.0 | 15.0 | 100 | 30.568 | 43.915 | 0.075 | 30.158 | 0.000 | 75.873 | 0.439 |
| 84 | 100 | 1000000 | 10000 | 5 | 50000 | 147599 | 5.0 | 15.0 | 100 | 30.998 | 43.560 | 0.075 | 30.572 | 0.000 | 75.171 | 0.436 |
| 85 | 100 | 1000000 | 10000 | 5 | 50000 | 5249 | 8.4 | 25.2 | 100 | 30.513 | 43.220 | 0.075 | 30.095 | 0.000 | 74.331 | 0.432 |
| 86 | 100 | 1000000 | 10000 | 5 | 50000 | 18671 | 8.4 | 25.2 | 100 | 30.230 | 43.622 | 0.073 | 29.768 | 0.000 | 74.462 | 0.436 |
| 87 | 100 | 1000000 | 10000 | 5 | 50000 | 147599 | 8.4 | 25.2 | 100 | 30.135 | 43.438 | 0.073 | 29.711 | 0.000 | 74.172 | 0.434 |
| 88 | 100 | 1000000 | 10000 | 5 | 50000 | 5249 | 15.0 | 45.0 | 100 | 30.803 | 43.294 | 0.073 | 30.395 | 0.000 | 74.670 | 0.433 |
| 89 | 100 | 1000000 | 10000 | 5 | 50000 | 18671 | 15.0 | 45.0 | 100 | 30.178 | 44.056 | 0.073 | 29.774 | 0.000 | 74.838 | 0.441 |
| 90 | 100 | 1000000 | 10000 | 5 | 50000 | 147599 | 15.0 | 45.0 | 100 | 30.327 | 43.927 | 0.076 | 29.929 | 0.000 | 74.825 | 0.439 |
| 91 | 100 | 1000000 | 100000 | 5 | 500000 | 5249 | 5.0 | 15.0 | 100 | 43.917 | 45.920 | 0.093 | 43.473 | 0.000 | 91.271 | 0.459 |
| 92 | 100 | 1000000 | 100000 | 5 | 500000 | 18671 | 5.0 | 15.0 | 100 | 45.407 | 46.599 | 0.072 | 45.003 | 0.000 | 92.585 | 0.466 |
| 93 | 100 | 1000000 | 100000 | 5 | 500000 | 147599 | 5.0 | 15.0 | 100 | 44.366 | 45.712 | 0.094 | 43.903 | 0.000 | 90.654 | 0.457 |
| 94 | 100 | 1000000 | 100000 | 5 | 500000 | 5249 | 8.4 | 25.2 | 100 | 44.529 | 46.057 | 0.097 | 44.104 | 0.000 | 91.139 | 0.461 |
| 95 | 100 | 1000000 | 100000 | 5 | 500000 | 18671 | 8.4 | 25.2 | 100 | 40.171 | 45.729 | 0.072 | 39.766 | 0.000 | 86.469 | 0.457 |
| 96 | 100 | 1000000 | 100000 | 5 | 500000 | 147599 | 8.4 | 25.2 | 100 | 41.008 | 45.931 | 0.073 | 40.582 | 0.000 | 87.556 | 0.459 |
| 97 | 100 | 1000000 | 100000 | 5 | 500000 | 5249 | 15.0 | 45.0 | 100 | 46.993 | 45.815 | 0.083 | 46.533 | 0.000 | 93.418 | 0.458 |
| 98 | 100 | 1000000 | 100000 | 5 | 500000 | 18671 | 15.0 | 45.0 | 100 | 47.984 | 45.789 | 0.076 | 47.582 | 0.000 | 94.420 | 0.458 |
| 99 | 100 | 1000000 | 100000 | 5 | 500000 | 147599 | 15.0 | 45.0 | 100 | 46.675 | 46.782 | 0.073 | 46.243 | 0.000 | 94.063 | 0.468 |
| 100 | 100 | 1000000 | 1000000 | 5 | 5000000 | 5249 | 5.0 | 15.0 | 100 | 191.452 | 72.080 | 0.075 | 191.028 | 0.000 | 264.120 | 0.721 |
| 101 | 100 | 1000000 | 1000000 | 5 | 5000000 | 18671 | 5.0 | 15.0 | 100 | 202.215 | 72.091 | 0.071 | 201.818 | 0.000 | 274.901 | 0.721 |
| 102 | 100 | 1000000 | 1000000 | 5 | 5000000 | 147599 | 5.0 | 15.0 | 100 | 194.156 | 70.305 | 0.367 | 193.061 | 0.000 | 265.111 | 0.703 |
| 103 | 100 | 1000000 | 1000000 | 5 | 5000000 | 5249 | 8.4 | 25.2 | 100 | 210.457 | 71.241 | 0.091 | 209.967 | 0.000 | 283.072 | 0.712 |
| 104 | 100 | 1000000 | 1000000 | 5 | 5000000 | 18671 | 8.4 | 25.2 | 100 | 191.631 | 72.264 | 0.073 | 191.224 | 0.000 | 264.496 | 0.723 |
| 105 | 100 | 1000000 | 1000000 | 5 | 5000000 | 147599 | 8.4 | 25.2 | 100 | 208.900 | 70.688 | 0.074 | 208.460 | 0.000 | 280.192 | 0.707 |
| 106 | 100 | 1000000 | 1000000 | 5 | 5000000 | 5249 | 15.0 | 45.0 | 100 | 193.560 | 72.103 | 0.073 | 193.155 | 0.000 | 266.227 | 0.721 |

Continued on next page

*(continued from previous page)*

| Run | $N$ | ECM | CELL | FOCAD/CELL | FOCAD | FNODE | $R_c$ | $R_s$ | Steps | $t_{\text{init}}$ | $t_{\text{sim}}$ | $t_{\text{RTC}}$ | $t_{\text{initfn}}$ | $t_{\text{exit}}$ | $t_{\text{total}}$ | $t_{\text{step}}$ |
|---|---|---|---|---|---|---|---|---|---|---|---|---|---|---|---|---|
| 107 | 100 | 1000000 | 1000000 | 5 | 5000000 | 18671 | 15.0 | 45.0 | 100 | 210.240 | 71.113 | 0.072 | 209.851 | 0.000 | 281.980 | 0.711 |
| 108 | 100 | 1000000 | 1000000 | 5 | 5000000 | 147599 | 15.0 | 45.0 | 100 | 186.345 | 71.792 | 0.078 | 185.929 | 0.000 | 258.781 | 0.718 |

Table 5: RTX5000 raw benchmark results. Times are in seconds. Compilation time ($t_{\mathrm{RTC}}$) is subtracted from $t_{\mathrm{init}}$ to compute actual initialization time.

| Run | $N$ | ECM | CELL | FOCAD/CELL | FOCAD | FNODE | $R_c$ | $R_s$ | Steps | $t_{\mathrm{init}}$ | $t_{\mathrm{sim}}$ | $t_{\mathrm{RTC}}$ | $t_{\mathrm{initfn}}$ | $t_{\mathrm{exit}}$ | $t_{\mathrm{total}}$ | $t_{\mathrm{step}}$ |
|---|---|---|---|---|---|---|---|---|---|---|---|---|---|---|---|---|
| 1 | 21 | 9261 | 1000 | 5 | 5000 | 5249 | 5.0 | 15.0 | 100 | 1.517 | 0.147 | 0.320 | 0.494 | 0.000 | 2.484 | 0.001 |
| 2 | 21 | 9261 | 1000 | 5 | 5000 | 18671 | 5.0 | 15.0 | 100 | 0.828 | 0.143 | 0.070 | 0.505 | 0.000 | 1.326 | 0.001 |
| 3 | 21 | 9261 | 1000 | 5 | 5000 | 147599 | 5.0 | 15.0 | 100 | 0.823 | 0.143 | 0.069 | 0.502 | 0.000 | 1.323 | 0.001 |
| 4 | 21 | 9261 | 1000 | 5 | 5000 | 5249 | 8.4 | 25.2 | 100 | 0.871 | 0.143 | 0.069 | 0.497 | 0.000 | 1.343 | 0.001 |
| 5 | 21 | 9261 | 1000 | 5 | 5000 | 18671 | 8.4 | 25.2 | 100 | 0.828 | 0.144 | 0.070 | 0.500 | 0.000 | 1.323 | 0.001 |
| 6 | 21 | 9261 | 1000 | 5 | 5000 | 147599 | 8.4 | 25.2 | 100 | 0.830 | 0.142 | 0.069 | 0.510 | 0.000 | 1.294 | 0.001 |
| 7 | 21 | 9261 | 1000 | 5 | 5000 | 5249 | 15.0 | 45.0 | 100 | 0.833 | 0.143 | 0.070 | 0.520 | 0.000 | 1.332 | 0.001 |
| 8 | 21 | 9261 | 1000 | 5 | 5000 | 18671 | 15.0 | 45.0 | 100 | 0.814 | 0.143 | 0.072 | 0.495 | 0.000 | 1.284 | 0.001 |
| 9 | 21 | 9261 | 1000 | 5 | 5000 | 147599 | 15.0 | 45.0 | 100 | 0.810 | 0.140 | 0.072 | 0.485 | 0.000 | 1.299 | 0.001 |
| 10 | 21 | 9261 | 10000 | 5 | 50000 | 5249 | 5.0 | 15.0 | 100 | 3.065 | 0.170 | 0.072 | 2.569 | 0.000 | 3.613 | 0.002 |
| 11 | 21 | 9261 | 10000 | 5 | 50000 | 18671 | 5.0 | 15.0 | 100 | 2.864 | 0.171 | 0.070 | 2.528 | 0.000 | 3.376 | 0.002 |
| 12 | 21 | 9261 | 10000 | 5 | 50000 | 147599 | 5.0 | 15.0 | 100 | 2.827 | 0.169 | 0.069 | 2.467 | 0.000 | 3.340 | 0.002 |
| 13 | 21 | 9261 | 10000 | 5 | 50000 | 5249 | 8.4 | 25.2 | 100 | 2.922 | 0.165 | 0.070 | 2.537 | 0.000 | 3.450 | 0.002 |
| 14 | 21 | 9261 | 10000 | 5 | 50000 | 18671 | 8.4 | 25.2 | 100 | 2.871 | 0.165 | 0.071 | 2.551 | 0.000 | 3.360 | 0.002 |
| 15 | 21 | 9261 | 10000 | 5 | 50000 | 147599 | 8.4 | 25.2 | 100 | 2.871 | 0.171 | 0.070 | 2.549 | 0.000 | 3.358 | 0.002 |
| 16 | 21 | 9261 | 10000 | 5 | 50000 | 5249 | 15.0 | 45.0 | 100 | 2.889 | 0.170 | 0.068 | 2.539 | 0.000 | 3.421 | 0.002 |
| 17 | 21 | 9261 | 10000 | 5 | 50000 | 18671 | 15.0 | 45.0 | 100 | 2.886 | 0.170 | 0.069 | 2.554 | 0.000 | 3.378 | 0.002 |
| 18 | 21 | 9261 | 10000 | 5 | 50000 | 147599 | 15.0 | 45.0 | 100 | 2.953 | 0.170 | 0.070 | 2.631 | 0.000 | 3.450 | 0.002 |
| 19 | 21 | 9261 | 100000 | 5 | 500000 | 5249 | 5.0 | 15.0 | 100 | 23.362 | 0.479 | 0.070 | 23.041 | 0.000 | 24.157 | 0.005 |
| 20 | 21 | 9261 | 100000 | 5 | 500000 | 18671 | 5.0 | 15.0 | 100 | 23.088 | 0.475 | 0.070 | 22.748 | 0.000 | 23.902 | 0.005 |
| 21 | 21 | 9261 | 100000 | 5 | 500000 | 147599 | 5.0 | 15.0 | 100 | 23.830 | 0.479 | 0.070 | 23.444 | 0.000 | 24.678 | 0.005 |
| 22 | 21 | 9261 | 100000 | 5 | 500000 | 5249 | 8.4 | 25.2 | 100 | 23.435 | 0.482 | 0.070 | 22.968 | 0.000 | 24.299 | 0.005 |
| 23 | 21 | 9261 | 100000 | 5 | 500000 | 18671 | 8.4 | 25.2 | 100 | 24.205 | 0.478 | 0.070 | 23.827 | 0.000 | 25.050 | 0.005 |
| 24 | 21 | 9261 | 100000 | 5 | 500000 | 147599 | 8.4 | 25.2 | 100 | 23.468 | 0.475 | 0.070 | 23.149 | 0.000 | 24.262 | 0.005 |
| 25 | 21 | 9261 | 100000 | 5 | 500000 | 5249 | 15.0 | 45.0 | 100 | 22.845 | 0.477 | 0.071 | 22.518 | 0.000 | 23.657 | 0.005 |
| 26 | 21 | 9261 | 100000 | 5 | 500000 | 18671 | 15.0 | 45.0 | 100 | 24.394 | 0.479 | 0.070 | 24.062 | 0.000 | 25.236 | 0.005 |
| 27 | 21 | 9261 | 100000 | 5 | 500000 | 147599 | 15.0 | 45.0 | 100 | 23.445 | 0.478 | 0.069 | 23.127 | 0.000 | 24.249 | 0.005 |
| 28 | 21 | 9261 | 1000000 | 5 | 5000000 | 5249 | 5.0 | 15.0 | 100 | 221.144 | 3.303 | 0.070 | 220.822 | 0.000 | 224.807 | 0.033 |
| 29 | 21 | 9261 | 1000000 | 5 | 5000000 | 18671 | 5.0 | 15.0 | 100 | 226.115 | 3.303 | 0.071 | 225.625 | 0.000 | 229.812 | 0.033 |
| 30 | 21 | 9261 | 1000000 | 5 | 5000000 | 147599 | 5.0 | 15.0 | 100 | 232.724 | 3.302 | 0.068 | 232.379 | 0.000 | 236.414 | 0.033 |
| 31 | 21 | 9261 | 1000000 | 5 | 5000000 | 5249 | 8.4 | 25.2 | 100 | 224.707 | 3.295 | 0.070 | 224.359 | 0.000 | 228.372 | 0.033 |
| 32 | 21 | 9261 | 1000000 | 5 | 5000000 | 18671 | 8.4 | 25.2 | 100 | 218.063 | 3.297 | 0.069 | 217.706 | 0.000 | 221.733 | 0.033 |
| 33 | 21 | 9261 | 1000000 | 5 | 5000000 | 147599 | 8.4 | 25.2 | 100 | 228.411 | 3.302 | 0.069 | 228.053 | 0.000 | 232.257 | 0.033 |
| 34 | 21 | 9261 | 1000000 | 5 | 5000000 | 5249 | 15.0 | 45.0 | 100 | 227.652 | 3.286 | 0.069 | 227.267 | 0.000 | 231.334 | 0.033 |
| 35 | 21 | 9261 | 1000000 | 5 | 5000000 | 18671 | 15.0 | 45.0 | 100 | 230.264 | 3.289 | 0.069 | 229.918 | 0.000 | 234.013 | 0.033 |
| 36 | 21 | 9261 | 1000000 | 5 | 5000000 | 147599 | 15.0 | 45.0 | 100 | 223.899 | 3.291 | 0.069 | 223.559 | 0.000 | 227.596 | 0.033 |
| 37 | 46 | 97336 | 1000 | 5 | 5000 | 5249 | 5.0 | 15.0 | 100 | 3.816 | 0.212 | 0.285 | 3.241 | 0.000 | 4.387 | 0.002 |
| 38 | 46 | 97336 | 1000 | 5 | 5000 | 18671 | 5.0 | 15.0 | 100 | 3.636 | 0.209 | 0.071 | 3.002 | 0.000 | 4.230 | 0.002 |
| 39 | 46 | 97336 | 1000 | 5 | 5000 | 147599 | 5.0 | 15.0 | 100 | 3.344 | 0.211 | 0.070 | 2.973 | 0.000 | 3.889 | 0.002 |
| 40 | 46 | 97336 | 1000 | 5 | 5000 | 5249 | 8.4 | 25.2 | 100 | 3.502 | 0.208 | 0.070 | 3.173 | 0.000 | 4.074 | 0.002 |
| 41 | 46 | 97336 | 1000 | 5 | 5000 | 18671 | 8.4 | 25.2 | 100 | 3.485 | 0.214 | 0.072 | 3.161 | 0.000 | 4.033 | 0.002 |
| 42 | 46 | 97336 | 1000 | 5 | 5000 | 147599 | 8.4 | 25.2 | 100 | 3.357 | 0.208 | 0.070 | 3.033 | 0.000 | 3.893 | 0.002 |
| 43 | 46 | 97336 | 1000 | 5 | 5000 | 5249 | 15.0 | 45.0 | 100 | 3.492 | 0.212 | 0.070 | 3.160 | 0.000 | 4.029 | 0.002 |
| 44 | 46 | 97336 | 1000 | 5 | 5000 | 18671 | 15.0 | 45.0 | 100 | 3.301 | 0.218 | 0.072 | 2.970 | 0.000 | 3.844 | 0.002 |
| 45 | 46 | 97336 | 1000 | 5 | 5000 | 147599 | 15.0 | 45.0 | 100 | 3.434 | 0.210 | 0.070 | 3.112 | 0.000 | 3.976 | 0.002 |
| 46 | 46 | 97336 | 10000 | 5 | 50000 | 5249 | 5.0 | 15.0 | 100 | 5.446 | 0.242 | 0.071 | 5.108 | 0.000 | 6.027 | 0.002 |
| 47 | 46 | 97336 | 10000 | 5 | 50000 | 18671 | 5.0 | 15.0 | 100 | 5.528 | 0.234 | 0.071 | 5.200 | 0.000 | 6.093 | 0.002 |
| 48 | 46 | 97336 | 10000 | 5 | 50000 | 147599 | 5.0 | 15.0 | 100 | 5.548 | 0.237 | 0.071 | 5.203 | 0.000 | 6.114 | 0.002 |
| 49 | 46 | 97336 | 10000 | 5 | 50000 | 5249 | 8.4 | 25.2 | 100 | 5.546 | 0.232 | 0.070 | 5.182 | 0.000 | 6.095 | 0.002 |
| 50 | 46 | 97336 | 10000 | 5 | 50000 | 18671 | 8.4 | 25.2 | 100 | 5.209 | 0.239 | 0.071 | 4.827 | 0.000 | 5.782 | 0.002 |
| 51 | 46 | 97336 | 10000 | 5 | 50000 | 147599 | 8.4 | 25.2 | 100 | 5.526 | 0.234 | 0.071 | 5.199 | 0.000 | 6.125 | 0.002 |
| 52 | 46 | 97336 | 10000 | 5 | 50000 | 5249 | 15.0 | 45.0 | 100 | 5.605 | 0.233 | 0.070 | 5.188 | 0.000 | 6.219 | 0.002 |

*Continued on next page*

*(continued from previous page)*

| Run | $N$ | ECM | CELL | FOCAD/CELL | FOCAD | FNODE | $R_c$ | $R_s$ | Steps | $t_{init}$ | $t_{sim}$ | $t_{RTC}$ | $t_{initfn}$ | $t_{exit}$ | $t_{total}$ | $t_{step}$ |
|---|---|---|---|---|---|---|---|---|---|---|---|---|---|---|---|---|
| 53 | 46 | 97336 | 10000 | 5 | 50000 | 18671 | 15.0 | 45.0 | 100 | 5.432 | 0.239 | 0.070 | 5.109 | 0.000 | 6.018 | 0.002 |
| 54 | 46 | 97336 | 10000 | 5 | 50000 | 147599 | 15.0 | 45.0 | 100 | 5.252 | 0.236 | 0.069 | 4.932 | 0.000 | 5.811 | 0.002 |
| 55 | 46 | 97336 | 100000 | 5 | 500000 | 5249 | 5.0 | 15.0 | 100 | 25.698 | 0.539 | 0.069 | 25.379 | 0.000 | 26.574 | 0.005 |
| 56 | 46 | 97336 | 100000 | 5 | 500000 | 18671 | 5.0 | 15.0 | 100 | 25.912 | 0.558 | 0.072 | 25.584 | 0.000 | 26.808 | 0.006 |
| 57 | 46 | 97336 | 100000 | 5 | 500000 | 147599 | 5.0 | 15.0 | 100 | 25.092 | 0.544 | 0.069 | 24.633 | 0.000 | 25.993 | 0.005 |
| 58 | 46 | 97336 | 100000 | 5 | 500000 | 5249 | 8.4 | 25.2 | 100 | 26.050 | 0.542 | 0.070 | 25.725 | 0.000 | 26.974 | 0.005 |
| 59 | 46 | 97336 | 100000 | 5 | 500000 | 18671 | 8.4 | 25.2 | 100 | 26.210 | 0.544 | 0.069 | 25.829 | 0.000 | 27.140 | 0.005 |
| 60 | 46 | 97336 | 100000 | 5 | 500000 | 147599 | 8.4 | 25.2 | 100 | 25.716 | 0.549 | 0.069 | 25.397 | 0.000 | 26.607 | 0.005 |
| 61 | 46 | 97336 | 100000 | 5 | 500000 | 5249 | 15.0 | 45.0 | 100 | 26.188 | 0.549 | 0.070 | 25.735 | 0.000 | 27.129 | 0.005 |
| 62 | 46 | 97336 | 100000 | 5 | 500000 | 18671 | 15.0 | 45.0 | 100 | 26.085 | 0.548 | 0.071 | 25.754 | 0.000 | 27.001 | 0.005 |
| 63 | 46 | 97336 | 100000 | 5 | 500000 | 147599 | 15.0 | 45.0 | 100 | 25.424 | 0.546 | 0.070 | 25.084 | 0.000 | 26.299 | 0.005 |
| 64 | 46 | 97336 | 1000000 | 5 | 5000000 | 5249 | 5.0 | 15.0 | 100 | 226.345 | 3.425 | 0.069 | 226.016 | 0.000 | 230.132 | 0.034 |
| 65 | 46 | 97336 | 1000000 | 5 | 5000000 | 18671 | 5.0 | 15.0 | 100 | 223.927 | 3.429 | 0.071 | 223.566 | 0.000 | 227.756 | 0.034 |
| 66 | 46 | 97336 | 1000000 | 5 | 5000000 | 147599 | 5.0 | 15.0 | 100 | 232.035 | 3.418 | 0.071 | 231.670 | 0.000 | 235.851 | 0.034 |
| 67 | 46 | 97336 | 1000000 | 5 | 5000000 | 5249 | 8.4 | 25.2 | 100 | 239.333 | 3.430 | 0.072 | 238.985 | 0.000 | 243.162 | 0.034 |
| 68 | 46 | 97336 | 1000000 | 5 | 5000000 | 18671 | 8.4 | 25.2 | 100 | 232.445 | 3.415 | 0.070 | 231.942 | 0.000 | 236.390 | 0.034 |
| 69 | 46 | 97336 | 1000000 | 5 | 5000000 | 147599 | 8.4 | 25.2 | 100 | 230.692 | 3.425 | 0.071 | 229.882 | 0.000 | 234.533 | 0.034 |
| 70 | 46 | 97336 | 1000000 | 5 | 5000000 | 5249 | 15.0 | 45.0 | 100 | 236.802 | 3.413 | 0.070 | 236.286 | 0.000 | 241.425 | 0.034 |
| 71 | 46 | 97336 | 1000000 | 5 | 5000000 | 18671 | 15.0 | 45.0 | 100 | 222.252 | 3.438 | 0.069 | 221.904 | 0.000 | 226.146 | 0.034 |
| 72 | 46 | 97336 | 1000000 | 5 | 5000000 | 147599 | 15.0 | 45.0 | 100 | 221.739 | 3.414 | 0.071 | 221.175 | 0.000 | 225.599 | 0.034 |
| 73 | 100 | 1000000 | 1000 | 5 | 5000 | 5249 | 5.0 | 15.0 | 100 | 94.945 | 1.105 | 66.807 | 27.865 | 0.000 | 96.460 | 0.011 |
| 74 | 100 | 1000000 | 1000 | 5 | 5000 | 18671 | 5.0 | 15.0 | 100 | 30.078 | 1.101 | 0.069 | 29.704 | 0.000 | 31.565 | 0.011 |
| 75 | 100 | 1000000 | 1000 | 5 | 5000 | 147599 | 5.0 | 15.0 | 100 | 30.202 | 1.101 | 0.070 | 29.872 | 0.000 | 31.656 | 0.011 |
| 76 | 100 | 1000000 | 1000 | 5 | 5000 | 5249 | 8.4 | 25.2 | 100 | 29.811 | 1.099 | 0.070 | 29.485 | 0.000 | 31.265 | 0.011 |
| 77 | 100 | 1000000 | 1000 | 5 | 5000 | 18671 | 8.4 | 25.2 | 100 | 28.602 | 1.102 | 0.071 | 28.281 | 0.000 | 30.054 | 0.011 |
| 78 | 100 | 1000000 | 1000 | 5 | 5000 | 147599 | 8.4 | 25.2 | 100 | 28.744 | 1.101 | 0.072 | 28.400 | 0.000 | 30.215 | 0.011 |
| 79 | 100 | 1000000 | 1000 | 5 | 5000 | 5249 | 15.0 | 45.0 | 100 | 29.070 | 1.102 | 0.070 | 28.622 | 0.000 | 30.574 | 0.011 |
| 80 | 100 | 1000000 | 1000 | 5 | 5000 | 18671 | 15.0 | 45.0 | 100 | 28.303 | 1.105 | 0.069 | 27.775 | 0.000 | 29.803 | 0.011 |
| 81 | 100 | 1000000 | 1000 | 5 | 5000 | 147599 | 15.0 | 45.0 | 100 | 28.653 | 1.105 | 0.069 | 28.336 | 0.000 | 30.107 | 0.011 |
| 82 | 100 | 1000000 | 10000 | 5 | 50000 | 5249 | 5.0 | 15.0 | 100 | 32.311 | 1.134 | 0.069 | 31.992 | 0.000 | 33.792 | 0.011 |
| 83 | 100 | 1000000 | 10000 | 5 | 50000 | 18671 | 5.0 | 15.0 | 100 | 32.742 | 1.141 | 0.072 | 32.404 | 0.000 | 34.240 | 0.011 |
| 84 | 100 | 1000000 | 10000 | 5 | 50000 | 147599 | 5.0 | 15.0 | 100 | 31.265 | 1.131 | 0.070 | 30.945 | 0.000 | 32.758 | 0.011 |
| 85 | 100 | 1000000 | 10000 | 5 | 50000 | 5249 | 8.4 | 25.2 | 100 | 32.461 | 1.125 | 0.070 | 32.142 | 0.000 | 33.928 | 0.011 |
| 86 | 100 | 1000000 | 10000 | 5 | 50000 | 18671 | 8.4 | 25.2 | 100 | 29.916 | 1.131 | 0.070 | 29.596 | 0.000 | 31.394 | 0.011 |
| 87 | 100 | 1000000 | 10000 | 5 | 50000 | 147599 | 8.4 | 25.2 | 100 | 31.562 | 1.125 | 0.070 | 31.242 | 0.000 | 33.093 | 0.011 |
| 88 | 100 | 1000000 | 10000 | 5 | 50000 | 5249 | 15.0 | 45.0 | 100 | 30.896 | 1.135 | 0.070 | 30.571 | 0.000 | 32.402 | 0.011 |
| 89 | 100 | 1000000 | 10000 | 5 | 50000 | 18671 | 15.0 | 45.0 | 100 | 31.523 | 1.127 | 0.069 | 31.144 | 0.000 | 33.006 | 0.011 |
| 90 | 100 | 1000000 | 10000 | 5 | 50000 | 147599 | 15.0 | 45.0 | 100 | 32.172 | 1.129 | 0.070 | 31.798 | 0.000 | 33.681 | 0.011 |
| 91 | 100 | 1000000 | 100000 | 5 | 500000 | 5249 | 5.0 | 15.0 | 100 | 52.332 | 1.479 | 0.070 | 52.013 | 0.000 | 54.199 | 0.015 |
| 92 | 100 | 1000000 | 100000 | 5 | 500000 | 18671 | 5.0 | 15.0 | 100 | 50.948 | 1.482 | 0.070 | 50.613 | 0.000 | 52.808 | 0.015 |
| 93 | 100 | 1000000 | 100000 | 5 | 500000 | 147599 | 5.0 | 15.0 | 100 | 51.016 | 1.478 | 0.070 | 50.689 | 0.000 | 52.828 | 0.015 |
| 94 | 100 | 1000000 | 100000 | 5 | 500000 | 5249 | 8.4 | 25.2 | 100 | 52.233 | 1.457 | 0.071 | 51.881 | 0.000 | 54.051 | 0.015 |
| 95 | 100 | 1000000 | 100000 | 5 | 500000 | 18671 | 8.4 | 25.2 | 100 | 54.260 | 1.464 | 0.070 | 53.839 | 0.000 | 56.165 | 0.015 |
| 96 | 100 | 1000000 | 100000 | 5 | 500000 | 147599 | 8.4 | 25.2 | 100 | 50.578 | 1.467 | 0.070 | 50.194 | 0.000 | 52.427 | 0.015 |
| 97 | 100 | 1000000 | 100000 | 5 | 500000 | 5249 | 15.0 | 45.0 | 100 | 52.248 | 1.467 | 0.070 | 51.930 | 0.000 | 54.049 | 0.015 |
| 98 | 100 | 1000000 | 100000 | 5 | 500000 | 18671 | 15.0 | 45.0 | 100 | 51.760 | 1.462 | 0.069 | 51.432 | 0.000 | 53.583 | 0.015 |
| 99 | 100 | 1000000 | 100000 | 5 | 500000 | 147599 | 15.0 | 45.0 | 100 | 51.327 | 1.469 | 0.070 | 50.970 | 0.000 | 53.144 | 0.015 |
| 100 | 100 | 1000000 | 1000000 | 5 | 5000000 | 5249 | 5.0 | 15.0 | 100 | 254.959 | 4.398 | 0.071 | 254.634 | 0.000 | 259.803 | 0.044 |
| 101 | 100 | 1000000 | 1000000 | 5 | 5000000 | 18671 | 5.0 | 15.0 | 100 | 252.768 | 4.398 | 0.070 | 252.418 | 0.000 | 257.570 | 0.044 |
| 102 | 100 | 1000000 | 1000000 | 5 | 5000000 | 147599 | 5.0 | 15.0 | 100 | 250.007 | 4.405 | 0.070 | 249.638 | 0.000 | 254.949 | 0.044 |
| 103 | 100 | 1000000 | 1000000 | 5 | 5000000 | 5249 | 8.4 | 25.2 | 100 | 253.408 | 4.377 | 0.070 | 253.043 | 0.000 | 258.219 | 0.044 |
| 104 | 100 | 1000000 | 1000000 | 5 | 5000000 | 18671 | 8.4 | 25.2 | 100 | 255.036 | 4.384 | 0.070 | 254.665 | 0.000 | 259.829 | 0.044 |
| 105 | 100 | 1000000 | 1000000 | 5 | 5000000 | 147599 | 8.4 | 25.2 | 100 | 259.215 | 4.402 | 0.070 | 258.693 | 0.000 | 264.087 | 0.044 |
| 106 | 100 | 1000000 | 1000000 | 5 | 5000000 | 5249 | 15.0 | 45.0 | 100 | 249.357 | 4.412 | 0.070 | 249.018 | 0.000 | 254.177 | 0.044 |

*Continued on next page*

*(continued from previous page)*

| Run | $N$ | ECM | CELL | FOCAD/CELL | FOCAD | FNODE | $R_c$ | $R_s$ | Steps | $t_{\text{init}}$ | $t_{\text{sim}}$ | $t_{\text{RTC}}$ | $t_{\text{initfn}}$ | $t_{\text{exit}}$ | $t_{\text{total}}$ | $t_{\text{step}}$ |
|---|---|---|---|---|---|---|---|---|---|---|---|---|---|---|---|---|
| 107 | 100 | 1000000 | 1000000 | 5 | 5000000 | 18671 | 15.0 | 45.0 | 100 | 246.071 | 4.383 | 0.069 | 245.712 | 0.000 | 250.821 | 0.044 |
| 108 | 100 | 1000000 | 1000000 | 5 | 5000000 | 147599 | 15.0 | 45.0 | 100 | 249.526 | 4.386 | 0.070 | 249.172 | 0.000 | 254.333 | 0.044 |

Table 6: A100 raw benchmark results. Times are in seconds. Compilation time ($t_{\mathrm{RTC}}$) is subtracted from $t_{\mathrm{init}}$ to compute actual initialization time.

| Run | $N$ | ECM | CELL | FOCAD/CELL | FOCAD | FNODE | $R_c$ | $R_s$ | Steps | $t_{\mathrm{init}}$ | $t_{\mathrm{sim}}$ | $t_{\mathrm{RTC}}$ | $t_{\mathrm{initfn}}$ | $t_{\mathrm{exit}}$ | $t_{\mathrm{total}}$ | $t_{\mathrm{step}}$ |
|---|---|---|---|---|---|---|---|---|---|---|---|---|---|---|---|---|
| 1 | 21 | 9261 | 1000 | 5 | 5000 | 5249 | 5.0 | 15.0 | 100 | 268.143 | 0.112 | 267.363 | 0.602 | 0.000 | 269.151 | 0.001 |
| 2 | 21 | 9261 | 1000 | 5 | 5000 | 18671 | 5.0 | 15.0 | 100 | 1.736 | 0.117 | 0.096 | 1.463 | 0.000 | 3.326 | 0.001 |
| 3 | 21 | 9261 | 1000 | 5 | 5000 | 147599 | 5.0 | 15.0 | 100 | 9.954 | 0.194 | 0.093 | 9.524 | 0.000 | 10.874 | 0.002 |
| 4 | 21 | 9261 | 1000 | 5 | 5000 | 5249 | 8.4 | 25.2 | 100 | 0.864 | 0.108 | 0.094 | 0.601 | 0.000 | 2.432 | 0.001 |
| 5 | 21 | 9261 | 1000 | 5 | 5000 | 18671 | 8.4 | 25.2 | 100 | 1.724 | 0.111 | 0.095 | 1.456 | 0.000 | 2.470 | 0.001 |
| 6 | 21 | 9261 | 1000 | 5 | 5000 | 147599 | 8.4 | 25.2 | 100 | 9.852 | 0.198 | 0.091 | 9.424 | 0.000 | 10.756 | 0.002 |
| 7 | 21 | 9261 | 1000 | 5 | 5000 | 5249 | 15.0 | 45.0 | 100 | 0.871 | 0.108 | 0.095 | 0.606 | 0.000 | 1.606 | 0.001 |
| 8 | 21 | 9261 | 1000 | 5 | 5000 | 18671 | 15.0 | 45.0 | 100 | 1.721 | 0.115 | 0.093 | 1.451 | 0.000 | 2.534 | 0.001 |
| 9 | 21 | 9261 | 1000 | 5 | 5000 | 147599 | 15.0 | 45.0 | 100 | 9.900 | 0.200 | 0.092 | 9.466 | 0.000 | 10.739 | 0.002 |
| 10 | 21 | 9261 | 1000 | 50 | 50000 | 5249 | 5.0 | 15.0 | 100 | 0.865 | 0.108 | 0.094 | 0.601 | 0.000 | 1.619 | 0.001 |
| 11 | 21 | 9261 | 1000 | 50 | 50000 | 18671 | 5.0 | 15.0 | 100 | 1.902 | 0.117 | 0.286 | 1.440 | 0.000 | 2.716 | 0.001 |
| 12 | 21 | 9261 | 1000 | 50 | 50000 | 147599 | 5.0 | 15.0 | 100 | 11.037 | 0.197 | 0.310 | 9.495 | 0.000 | 12.523 | 0.002 |
| 13 | 21 | 9261 | 1000 | 50 | 50000 | 5249 | 8.4 | 25.2 | 100 | 1.078 | 0.114 | 0.296 | 0.612 | 0.000 | 1.900 | 0.001 |
| 14 | 21 | 9261 | 1000 | 50 | 50000 | 18671 | 8.4 | 25.2 | 100 | 1.717 | 0.117 | 0.094 | 1.443 | 0.000 | 2.508 | 0.001 |
| 15 | 21 | 9261 | 1000 | 50 | 50000 | 147599 | 8.4 | 25.2 | 100 | 10.064 | 0.200 | 0.092 | 9.631 | 0.000 | 11.002 | 0.002 |
| 16 | 21 | 9261 | 1000 | 50 | 50000 | 5249 | 15.0 | 45.0 | 100 | 0.887 | 0.114 | 0.093 | 0.622 | 0.000 | 1.695 | 0.001 |
| 17 | 21 | 9261 | 1000 | 50 | 50000 | 18671 | 15.0 | 45.0 | 100 | 1.735 | 0.117 | 0.095 | 1.456 | 0.000 | 2.694 | 0.001 |
| 18 | 21 | 9261 | 1000 | 50 | 50000 | 147599 | 15.0 | 45.0 | 100 | 10.073 | 0.195 | 0.092 | 9.631 | 0.000 | 10.936 | 0.002 |
| 19 | 21 | 9261 | 10000 | 5 | 50000 | 5249 | 5.0 | 15.0 | 100 | 0.864 | 0.114 | 0.093 | 0.599 | 0.000 | 1.677 | 0.001 |
| 20 | 21 | 9261 | 10000 | 5 | 50000 | 18671 | 5.0 | 15.0 | 100 | 1.710 | 0.110 | 0.092 | 1.445 | 0.000 | 2.562 | 0.001 |
| 21 | 21 | 9261 | 10000 | 5 | 50000 | 147599 | 5.0 | 15.0 | 100 | 10.604 | 0.194 | 0.095 | 9.789 | 0.000 | 11.493 | 0.002 |
| 22 | 21 | 9261 | 10000 | 5 | 50000 | 5249 | 8.4 | 25.2 | 100 | 0.871 | 0.112 | 0.095 | 0.606 | 0.000 | 1.554 | 0.001 |
| 23 | 21 | 9261 | 10000 | 5 | 50000 | 18671 | 8.4 | 25.2 | 100 | 1.769 | 0.111 | 0.094 | 1.501 | 0.000 | 2.513 | 0.001 |
| 24 | 21 | 9261 | 10000 | 5 | 50000 | 147599 | 8.4 | 25.2 | 100 | 10.305 | 0.195 | 0.093 | 9.872 | 0.000 | 11.180 | 0.002 |
| 25 | 21 | 9261 | 10000 | 5 | 50000 | 5249 | 15.0 | 45.0 | 100 | 0.871 | 0.107 | 0.094 | 0.605 | 0.000 | 1.608 | 0.001 |
| 26 | 21 | 9261 | 10000 | 5 | 50000 | 18671 | 15.0 | 45.0 | 100 | 1.703 | 0.117 | 0.095 | 1.428 | 0.000 | 2.451 | 0.001 |
| 27 | 21 | 9261 | 10000 | 5 | 50000 | 147599 | 15.0 | 45.0 | 100 | 10.062 | 0.194 | 0.092 | 9.635 | 0.000 | 10.875 | 0.002 |
| 28 | 21 | 9261 | 10000 | 50 | 500000 | 5249 | 5.0 | 15.0 | 100 | 1.071 | 0.113 | 0.295 | 0.603 | 0.000 | 1.896 | 0.001 |
| 29 | 21 | 9261 | 10000 | 50 | 500000 | 18671 | 5.0 | 15.0 | 100 | 2.002 | 0.116 | 0.297 | 1.450 | 0.000 | 2.730 | 0.001 |
| 30 | 21 | 9261 | 10000 | 50 | 500000 | 147599 | 5.0 | 15.0 | 100 | 10.585 | 0.201 | 0.297 | 9.665 | 0.000 | 11.644 | 0.002 |
| 31 | 21 | 9261 | 10000 | 50 | 500000 | 5249 | 8.4 | 25.2 | 100 | 0.884 | 0.108 | 0.097 | 0.610 | 0.000 | 2.453 | 0.001 |
| 32 | 21 | 9261 | 10000 | 50 | 500000 | 18671 | 8.4 | 25.2 | 100 | 1.708 | 0.111 | 0.093 | 1.438 | 0.000 | 2.480 | 0.001 |
| 33 | 21 | 9261 | 10000 | 50 | 500000 | 147599 | 8.4 | 25.2 | 100 | 10.170 | 0.197 | 0.093 | 9.740 | 0.000 | 10.988 | 0.002 |
| 34 | 21 | 9261 | 10000 | 50 | 500000 | 5249 | 15.0 | 45.0 | 100 | 0.869 | 0.113 | 0.093 | 0.604 | 0.000 | 2.506 | 0.001 |
| 35 | 21 | 9261 | 10000 | 50 | 500000 | 18671 | 15.0 | 45.0 | 100 | 1.724 | 0.111 | 0.094 | 1.453 | 0.000 | 2.457 | 0.001 |
| 36 | 21 | 9261 | 10000 | 50 | 500000 | 147599 | 15.0 | 45.0 | 100 | 10.113 | 0.195 | 0.094 | 9.678 | 0.000 | 10.964 | 0.002 |
| 37 | 21 | 9261 | 100000 | 5 | 500000 | 5249 | 5.0 | 15.0 | 100 | 0.866 | 0.114 | 0.095 | 0.601 | 0.000 | 1.621 | 0.001 |
| 38 | 21 | 9261 | 100000 | 5 | 500000 | 18671 | 5.0 | 15.0 | 100 | 1.714 | 0.111 | 0.093 | 1.447 | 0.000 | 2.434 | 0.001 |
| 39 | 21 | 9261 | 100000 | 5 | 500000 | 147599 | 5.0 | 15.0 | 100 | 10.101 | 0.197 | 0.092 | 9.667 | 0.000 | 10.925 | 0.002 |
| 40 | 21 | 9261 | 100000 | 5 | 500000 | 5249 | 8.4 | 25.2 | 100 | 1.169 | 0.112 | 0.384 | 0.615 | 0.000 | 2.003 | 0.001 |
| 41 | 21 | 9261 | 100000 | 5 | 500000 | 18671 | 8.4 | 25.2 | 100 | 1.733 | 0.118 | 0.094 | 1.464 | 0.000 | 2.469 | 0.001 |
| 42 | 21 | 9261 | 100000 | 5 | 500000 | 147599 | 8.4 | 25.2 | 100 | 10.046 | 0.196 | 0.093 | 9.612 | 0.000 | 10.968 | 0.002 |
| 43 | 21 | 9261 | 100000 | 5 | 500000 | 5249 | 15.0 | 45.0 | 100 | 0.876 | 0.111 | 0.094 | 0.610 | 0.000 | 2.529 | 0.001 |
| 44 | 21 | 9261 | 100000 | 5 | 500000 | 18671 | 15.0 | 45.0 | 100 | 1.719 | 0.117 | 0.092 | 1.452 | 0.000 | 2.459 | 0.001 |
| 45 | 21 | 9261 | 100000 | 5 | 500000 | 147599 | 15.0 | 45.0 | 100 | 10.668 | 0.195 | 0.319 | 9.640 | 0.000 | 11.517 | 0.002 |
| 46 | 21 | 9261 | 100000 | 50 | 5000000 | 5249 | 5.0 | 15.0 | 100 | 0.862 | 0.113 | 0.095 | 0.598 | 0.000 | 2.500 | 0.001 |
| 47 | 21 | 9261 | 100000 | 50 | 5000000 | 18671 | 5.0 | 15.0 | 100 | 1.703 | 0.112 | 0.093 | 1.436 | 0.000 | 2.486 | 0.001 |
| 48 | 21 | 9261 | 100000 | 50 | 5000000 | 147599 | 5.0 | 15.0 | 100 | 10.860 | 0.196 | 0.867 | 9.646 | 0.000 | 11.739 | 0.002 |
| 49 | 21 | 9261 | 100000 | 50 | 5000000 | 5249 | 8.4 | 25.2 | 100 | 0.863 | 0.108 | 0.096 | 0.597 | 0.000 | 1.572 | 0.001 |
| 50 | 21 | 9261 | 100000 | 50 | 5000000 | 18671 | 8.4 | 25.2 | 100 | 1.703 | 0.111 | 0.096 | 1.422 | 0.000 | 2.430 | 0.001 |
| 51 | 21 | 9261 | 100000 | 50 | 5000000 | 147599 | 8.4 | 25.2 | 100 | 10.021 | 0.202 | 0.092 | 9.587 | 0.000 | 10.945 | 0.002 |
| 52 | 21 | 9261 | 100000 | 50 | 5000000 | 5249 | 15.0 | 45.0 | 100 | 0.870 | 0.108 | 0.095 | 0.604 | 0.000 | 1.576 | 0.001 |

*Continued on next page*

(continued from previous page)

| Run | $N$ | ECM | CELL | FOCAD/CELL | FOCAD | FNODE | $R_c$ | $R_s$ | Steps | $t_{\text{init}}$ | $t_{\text{sim}}$ | $t_{\text{RTC}}$ | $t_{\text{initfn}}$ | $t_{\text{exit}}$ | $t_{\text{total}}$ | $t_{\text{step}}$ |
|---|---|---|---|---|---|---|---|---|---|---|---|---|---|---|---|---|
| 53 | 21 | 9261 | 100000 | 50 | 5000000 | 18671 | 15.0 | 45.0 | 100 | 1.707 | 0.114 | 0.093 | 1.435 | 0.000 | 2.469 | 0.001 |
| 54 | 21 | 9261 | 100000 | 50 | 5000000 | 147599 | 15.0 | 45.0 | 100 | 10.071 | 0.199 | 0.094 | 9.634 | 0.000 | 11.125 | 0.002 |
| 55 | 21 | 9261 | 1000000 | 5 | 5000000 | 5249 | 5.0 | 15.0 | 100 | 2.368 | 0.111 | 0.140 | 0.601 | 0.000 | 3.152 | 0.001 |
| 56 | 21 | 9261 | 1000000 | 5 | 5000000 | 18671 | 5.0 | 15.0 | 100 | 1.729 | 0.118 | 0.093 | 1.461 | 0.000 | 2.529 | 0.001 |
| 57 | 21 | 9261 | 1000000 | 5 | 5000000 | 147599 | 5.0 | 15.0 | 100 | 10.170 | 0.197 | 0.092 | 9.738 | 0.000 | 10.934 | 0.002 |
| 58 | 21 | 9261 | 1000000 | 5 | 5000000 | 5249 | 8.4 | 25.2 | 100 | 0.859 | 0.113 | 0.095 | 0.595 | 0.000 | 1.545 | 0.001 |
| 59 | 21 | 9261 | 1000000 | 5 | 5000000 | 18671 | 8.4 | 25.2 | 100 | 1.916 | 0.117 | 0.278 | 1.465 | 0.000 | 2.609 | 0.001 |
| 60 | 21 | 9261 | 1000000 | 5 | 5000000 | 147599 | 8.4 | 25.2 | 100 | 10.074 | 0.195 | 0.093 | 9.637 | 0.000 | 11.752 | 0.002 |
| 61 | 21 | 9261 | 1000000 | 5 | 5000000 | 5249 | 15.0 | 45.0 | 100 | 0.875 | 0.111 | 0.094 | 0.608 | 0.000 | 1.683 | 0.001 |
| 62 | 21 | 9261 | 1000000 | 5 | 5000000 | 18671 | 15.0 | 45.0 | 100 | 1.696 | 0.117 | 0.092 | 1.430 | 0.000 | 2.522 | 0.001 |
| 63 | 21 | 9261 | 1000000 | 5 | 5000000 | 147599 | 15.0 | 45.0 | 100 | 10.023 | 0.202 | 0.095 | 9.586 | 0.000 | 11.753 | 0.002 |
| 64 | 21 | 9261 | 1000000 | 50 | 50000000 | 5249 | 5.0 | 15.0 | 100 | 0.855 | 0.109 | 0.093 | 0.596 | 0.000 | 2.502 | 0.001 |
| 65 | 21 | 9261 | 1000000 | 50 | 50000000 | 18671 | 5.0 | 15.0 | 100 | 1.702 | 0.117 | 0.093 | 1.432 | 0.000 | 2.428 | 0.001 |
| 66 | 21 | 9261 | 1000000 | 50 | 50000000 | 147599 | 5.0 | 15.0 | 100 | 9.963 | 0.196 | 0.093 | 9.525 | 0.000 | 10.737 | 0.002 |
| 67 | 21 | 9261 | 1000000 | 50 | 50000000 | 5249 | 8.4 | 25.2 | 100 | 0.865 | 0.113 | 0.096 | 0.599 | 0.000 | 2.442 | 0.001 |
| 68 | 21 | 9261 | 1000000 | 50 | 50000000 | 18671 | 8.4 | 25.2 | 100 | 1.736 | 0.118 | 0.094 | 1.468 | 0.000 | 2.492 | 0.001 |
| 69 | 21 | 9261 | 1000000 | 50 | 50000000 | 147599 | 8.4 | 25.2 | 100 | 10.075 | 0.196 | 0.092 | 9.644 | 0.000 | 11.734 | 0.002 |
| 70 | 21 | 9261 | 1000000 | 50 | 50000000 | 5249 | 15.0 | 45.0 | 100 | 0.862 | 0.111 | 0.093 | 0.602 | 0.000 | 1.577 | 0.001 |
| 71 | 21 | 9261 | 1000000 | 50 | 50000000 | 18671 | 15.0 | 45.0 | 100 | 1.748 | 0.112 | 0.096 | 1.469 | 0.000 | 3.321 | 0.001 |
| 72 | 21 | 9261 | 1000000 | 50 | 50000000 | 147599 | 15.0 | 45.0 | 100 | 9.909 | 0.201 | 0.094 | 9.474 | 0.000 | 10.744 | 0.002 |
| 73 | 46 | 97336 | 1000 | 5 | 5000 | 5249 | 5.0 | 15.0 | 100 | 272.360 | 0.112 | 269.005 | 3.184 | 0.000 | 273.860 | 0.001 |
| 74 | 46 | 97336 | 1000 | 5 | 5000 | 18671 | 5.0 | 15.0 | 100 | 4.390 | 0.117 | 0.097 | 4.110 | 0.000 | 5.235 | 0.001 |
| 75 | 46 | 97336 | 1000 | 5 | 5000 | 147599 | 5.0 | 15.0 | 100 | 12.586 | 0.196 | 0.093 | 12.154 | 0.000 | 13.375 | 0.002 |
| 76 | 46 | 97336 | 1000 | 5 | 5000 | 5249 | 8.4 | 25.2 | 100 | 3.492 | 0.113 | 0.098 | 3.223 | 0.000 | 4.211 | 0.001 |
| 77 | 46 | 97336 | 1000 | 5 | 5000 | 18671 | 8.4 | 25.2 | 100 | 4.779 | 0.116 | 0.428 | 4.166 | 0.000 | 5.470 | 0.001 |
| 78 | 46 | 97336 | 1000 | 5 | 5000 | 147599 | 8.4 | 25.2 | 100 | 13.814 | 0.197 | 0.903 | 12.399 | 0.000 | 14.663 | 0.002 |
| 79 | 46 | 97336 | 1000 | 5 | 5000 | 5249 | 15.0 | 45.0 | 100 | 3.495 | 0.114 | 0.092 | 3.233 | 0.000 | 4.235 | 0.001 |
| 80 | 46 | 97336 | 1000 | 5 | 5000 | 18671 | 15.0 | 45.0 | 100 | 4.545 | 0.117 | 0.096 | 4.091 | 0.000 | 5.258 | 0.001 |
| 81 | 46 | 97336 | 1000 | 5 | 5000 | 147599 | 15.0 | 45.0 | 100 | 12.870 | 0.199 | 0.094 | 12.430 | 0.000 | 13.638 | 0.002 |
| 82 | 46 | 97336 | 1000 | 50 | 50000 | 5249 | 5.0 | 15.0 | 100 | 3.514 | 0.114 | 0.096 | 3.249 | 0.000 | 5.102 | 0.001 |
| 83 | 46 | 97336 | 1000 | 50 | 50000 | 18671 | 5.0 | 15.0 | 100 | 4.452 | 0.118 | 0.096 | 4.175 | 0.000 | 5.251 | 0.001 |
| 84 | 46 | 97336 | 1000 | 50 | 50000 | 147599 | 5.0 | 15.0 | 100 | 12.572 | 0.200 | 0.124 | 12.086 | 0.000 | 14.315 | 0.002 |
| 85 | 46 | 97336 | 1000 | 50 | 50000 | 5249 | 8.4 | 25.2 | 100 | 3.598 | 0.115 | 0.175 | 3.251 | 0.000 | 4.307 | 0.001 |
| 86 | 46 | 97336 | 1000 | 50 | 50000 | 18671 | 8.4 | 25.2 | 100 | 4.341 | 0.118 | 0.095 | 4.065 | 0.000 | 5.247 | 0.001 |
| 87 | 46 | 97336 | 1000 | 50 | 50000 | 147599 | 8.4 | 25.2 | 100 | 12.619 | 0.203 | 0.094 | 12.186 | 0.000 | 13.401 | 0.002 |
| 88 | 46 | 97336 | 1000 | 50 | 50000 | 5249 | 15.0 | 45.0 | 100 | 3.490 | 0.114 | 0.095 | 3.224 | 0.000 | 4.173 | 0.001 |
| 89 | 46 | 97336 | 1000 | 50 | 50000 | 18671 | 15.0 | 45.0 | 100 | 4.383 | 0.116 | 0.096 | 4.110 | 0.000 | 5.051 | 0.001 |
| 90 | 46 | 97336 | 1000 | 50 | 50000 | 147599 | 15.0 | 45.0 | 100 | 13.101 | 0.197 | 0.097 | 12.311 | 0.000 | 13.944 | 0.002 |
| 91 | 46 | 97336 | 10000 | 5 | 50000 | 5249 | 5.0 | 15.0 | 100 | 3.487 | 0.114 | 0.096 | 3.219 | 0.000 | 5.129 | 0.001 |
| 92 | 46 | 97336 | 10000 | 5 | 50000 | 18671 | 5.0 | 15.0 | 100 | 4.416 | 0.118 | 0.095 | 4.144 | 0.000 | 5.116 | 0.001 |
| 93 | 46 | 97336 | 10000 | 5 | 50000 | 147599 | 5.0 | 15.0 | 100 | 12.793 | 0.202 | 0.094 | 12.357 | 0.000 | 13.579 | 0.002 |
| 94 | 46 | 97336 | 10000 | 5 | 50000 | 5249 | 8.4 | 25.2 | 100 | 3.548 | 0.113 | 0.095 | 3.285 | 0.000 | 4.244 | 0.001 |
| 95 | 46 | 97336 | 10000 | 5 | 50000 | 18671 | 8.4 | 25.2 | 100 | 4.373 | 0.118 | 0.096 | 4.099 | 0.000 | 5.061 | 0.001 |
| 96 | 46 | 97336 | 10000 | 5 | 50000 | 147599 | 8.4 | 25.2 | 100 | 12.626 | 0.198 | 0.099 | 12.162 | 0.000 | 13.482 | 0.002 |
| 97 | 46 | 97336 | 10000 | 5 | 50000 | 5249 | 15.0 | 45.0 | 100 | 3.618 | 0.113 | 0.094 | 3.349 | 0.000 | 4.366 | 0.001 |
| 98 | 46 | 97336 | 10000 | 5 | 50000 | 18671 | 15.0 | 45.0 | 100 | 4.400 | 0.118 | 0.096 | 4.126 | 0.000 | 5.997 | 0.001 |
| 99 | 46 | 97336 | 10000 | 5 | 50000 | 147599 | 15.0 | 45.0 | 100 | 12.527 | 0.198 | 0.094 | 12.094 | 0.000 | 14.254 | 0.002 |
| 100 | 46 | 97336 | 10000 | 50 | 500000 | 5249 | 5.0 | 15.0 | 100 | 3.528 | 0.115 | 0.094 | 3.262 | 0.000 | 5.129 | 0.001 |
| 101 | 46 | 97336 | 10000 | 50 | 500000 | 18671 | 5.0 | 15.0 | 100 | 4.380 | 0.118 | 0.098 | 4.104 | 0.000 | 5.093 | 0.001 |
| 102 | 46 | 97336 | 10000 | 50 | 500000 | 147599 | 5.0 | 15.0 | 100 | 12.946 | 0.198 | 0.336 | 12.259 | 0.000 | 13.857 | 0.002 |
| 103 | 46 | 97336 | 10000 | 50 | 500000 | 5249 | 8.4 | 25.2 | 100 | 3.718 | 0.113 | 0.096 | 3.245 | 0.000 | 4.447 | 0.001 |
| 104 | 46 | 97336 | 10000 | 50 | 500000 | 18671 | 8.4 | 25.2 | 100 | 4.424 | 0.118 | 0.098 | 4.065 | 0.000 | 5.127 | 0.001 |
| 105 | 46 | 97336 | 10000 | 50 | 500000 | 147599 | 8.4 | 25.2 | 100 | 12.736 | 0.203 | 0.094 | 12.306 | 0.000 | 13.500 | 0.002 |
| 106 | 46 | 97336 | 10000 | 50 | 500000 | 5249 | 15.0 | 45.0 | 100 | 3.474 | 0.115 | 0.095 | 3.207 | 0.000 | 4.167 | 0.001 |

Continued on next page

(continued from previous page)

| Run | $N$ | ECM | CELL | FOCAD/CELL | FOCAD | FNODE | $R_c$ | $R_s$ | Steps | $t_{\text{init}}$ | $t_{\text{sim}}$ | $t_{\text{RTC}}$ | $t_{\text{initfn}}$ | $t_{\text{exit}}$ | $t_{\text{total}}$ | $t_{\text{step}}$ |
|---|---|---|---|---|---|---|---|---|---|---|---|---|---|---|---|---|
| 107 | 46 | 97336 | 10000 | 50 | 500000 | 18671 | 15.0 | 45.0 | 100 | 4.380 | 0.117 | 0.094 | 4.107 | 0.000 | 5.979 | 0.001 |
| 108 | 46 | 97336 | 10000 | 50 | 500000 | 147599 | 15.0 | 45.0 | 100 | 12.605 | 0.198 | 0.094 | 12.173 | 0.000 | 13.406 | 0.002 |
| 109 | 46 | 97336 | 100000 | 5 | 500000 | 5249 | 5.0 | 15.0 | 100 | 3.509 | 0.113 | 0.094 | 3.243 | 0.000 | 5.091 | 0.001 |
| 110 | 46 | 97336 | 100000 | 5 | 500000 | 18671 | 5.0 | 15.0 | 100 | 4.436 | 0.117 | 0.097 | 4.161 | 0.000 | 5.141 | 0.001 |
| 111 | 46 | 97336 | 100000 | 5 | 500000 | 147599 | 5.0 | 15.0 | 100 | 12.596 | 0.202 | 0.095 | 12.161 | 0.000 | 13.437 | 0.002 |
| 112 | 46 | 97336 | 100000 | 5 | 500000 | 5249 | 8.4 | 25.2 | 100 | 3.437 | 0.115 | 0.093 | 3.175 | 0.000 | 4.177 | 0.001 |
| 113 | 46 | 97336 | 100000 | 5 | 500000 | 18671 | 8.4 | 25.2 | 100 | 4.371 | 0.118 | 0.097 | 4.098 | 0.000 | 5.958 | 0.001 |
| 114 | 46 | 97336 | 100000 | 5 | 500000 | 147599 | 8.4 | 25.2 | 100 | 12.710 | 0.198 | 0.094 | 12.272 | 0.000 | 13.507 | 0.002 |
| 115 | 46 | 97336 | 100000 | 5 | 500000 | 5249 | 15.0 | 45.0 | 100 | 3.531 | 0.113 | 0.095 | 3.262 | 0.000 | 4.260 | 0.001 |
| 116 | 46 | 97336 | 100000 | 5 | 500000 | 18671 | 15.0 | 45.0 | 100 | 4.401 | 0.118 | 0.096 | 4.128 | 0.000 | 5.171 | 0.001 |
| 117 | 46 | 97336 | 100000 | 5 | 500000 | 147599 | 15.0 | 45.0 | 100 | 12.940 | 0.196 | 0.095 | 12.417 | 0.000 | 13.725 | 0.002 |
| 118 | 46 | 97336 | 100000 | 50 | 5000000 | 5249 | 5.0 | 15.0 | 100 | 3.456 | 0.114 | 0.096 | 3.191 | 0.000 | 4.136 | 0.001 |
| 119 | 46 | 97336 | 100000 | 50 | 5000000 | 18671 | 5.0 | 15.0 | 100 | 4.317 | 0.118 | 0.097 | 4.043 | 0.000 | 5.054 | 0.001 |
| 120 | 46 | 97336 | 100000 | 50 | 5000000 | 147599 | 5.0 | 15.0 | 100 | 12.612 | 0.199 | 0.094 | 12.176 | 0.000 | 13.408 | 0.002 |
| 121 | 46 | 97336 | 100000 | 50 | 5000000 | 5249 | 8.4 | 25.2 | 100 | 3.506 | 0.113 | 0.095 | 3.241 | 0.000 | 4.202 | 0.001 |
| 122 | 46 | 97336 | 100000 | 50 | 5000000 | 18671 | 8.4 | 25.2 | 100 | 4.299 | 0.116 | 0.096 | 4.024 | 0.000 | 5.043 | 0.001 |
| 123 | 46 | 97336 | 100000 | 50 | 5000000 | 147599 | 8.4 | 25.2 | 100 | 12.675 | 0.202 | 0.094 | 12.241 | 0.000 | 13.564 | 0.002 |
| 124 | 46 | 97336 | 100000 | 50 | 5000000 | 5249 | 15.0 | 45.0 | 100 | 3.535 | 0.114 | 0.096 | 3.266 | 0.000 | 4.252 | 0.001 |
| 125 | 46 | 97336 | 100000 | 50 | 5000000 | 18671 | 15.0 | 45.0 | 100 | 4.330 | 0.117 | 0.095 | 4.058 | 0.000 | 5.990 | 0.001 |
| 126 | 46 | 97336 | 100000 | 50 | 5000000 | 147599 | 15.0 | 45.0 | 100 | 12.645 | 0.207 | 0.098 | 12.192 | 0.000 | 13.515 | 0.002 |
| 127 | 46 | 97336 | 1000000 | 5 | 5000000 | 5249 | 5.0 | 15.0 | 100 | 3.625 | 0.116 | 0.094 | 3.357 | 0.000 | 4.393 | 0.001 |
| 128 | 46 | 97336 | 1000000 | 5 | 5000000 | 18671 | 5.0 | 15.0 | 100 | 4.407 | 0.116 | 0.096 | 4.085 | 0.000 | 5.147 | 0.001 |
| 129 | 46 | 97336 | 1000000 | 5 | 5000000 | 147599 | 5.0 | 15.0 | 100 | 12.665 | 0.198 | 0.094 | 12.232 | 0.000 | 13.572 | 0.002 |
| 130 | 46 | 97336 | 1000000 | 5 | 5000000 | 5249 | 8.4 | 25.2 | 100 | 3.510 | 0.114 | 0.094 | 3.245 | 0.000 | 4.270 | 0.001 |
| 131 | 46 | 97336 | 1000000 | 5 | 5000000 | 18671 | 8.4 | 25.2 | 100 | 4.420 | 0.118 | 0.113 | 4.130 | 0.000 | 5.854 | 0.001 |
| 132 | 46 | 97336 | 1000000 | 5 | 5000000 | 147599 | 8.4 | 25.2 | 100 | 12.777 | 0.199 | 0.094 | 12.339 | 0.000 | 13.568 | 0.002 |
| 133 | 46 | 97336 | 1000000 | 5 | 5000000 | 5249 | 15.0 | 45.0 | 100 | 3.750 | 0.115 | 0.099 | 3.478 | 0.000 | 4.449 | 0.001 |
| 134 | 46 | 97336 | 1000000 | 5 | 5000000 | 18671 | 15.0 | 45.0 | 100 | 4.391 | 0.117 | 0.096 | 4.115 | 0.000 | 5.083 | 0.001 |
| 135 | 46 | 97336 | 1000000 | 5 | 5000000 | 147599 | 15.0 | 45.0 | 100 | 12.594 | 0.200 | 0.094 | 12.156 | 0.000 | 13.372 | 0.002 |
| 136 | 46 | 97336 | 1000000 | 50 | 50000000 | 5249 | 5.0 | 15.0 | 100 | 3.496 | 0.113 | 0.097 | 3.227 | 0.000 | 4.221 | 0.001 |
| 137 | 46 | 97336 | 1000000 | 50 | 50000000 | 18671 | 5.0 | 15.0 | 100 | 4.387 | 0.118 | 0.098 | 4.111 | 0.000 | 5.086 | 0.001 |
| 138 | 46 | 97336 | 1000000 | 50 | 50000000 | 147599 | 5.0 | 15.0 | 100 | 12.599 | 0.202 | 0.095 | 12.157 | 0.000 | 13.382 | 0.002 |
| 139 | 46 | 97336 | 1000000 | 50 | 50000000 | 5249 | 8.4 | 25.2 | 100 | 3.468 | 0.113 | 0.098 | 3.193 | 0.000 | 4.142 | 0.001 |
| 140 | 46 | 97336 | 1000000 | 50 | 50000000 | 18671 | 8.4 | 25.2 | 100 | 4.320 | 0.115 | 0.097 | 4.042 | 0.000 | 5.080 | 0.001 |
| 141 | 46 | 97336 | 1000000 | 50 | 50000000 | 147599 | 8.4 | 25.2 | 100 | 12.621 | 0.196 | 0.099 | 12.173 | 0.000 | 13.463 | 0.002 |
| 142 | 46 | 97336 | 1000000 | 50 | 50000000 | 5249 | 15.0 | 45.0 | 100 | 3.508 | 0.114 | 0.097 | 3.232 | 0.000 | 4.189 | 0.001 |
| 143 | 46 | 97336 | 1000000 | 50 | 50000000 | 18671 | 15.0 | 45.0 | 100 | 4.494 | 0.118 | 0.107 | 4.111 | 0.000 | 5.199 | 0.001 |
| 144 | 46 | 97336 | 1000000 | 50 | 50000000 | 147599 | 15.0 | 45.0 | 100 | 12.674 | 0.198 | 0.094 | 12.237 | 0.000 | 13.556 | 0.002 |
| 145 | 100 | 1000000 | 1000 | 5 | 5000 | 5249 | 5.0 | 15.0 | 100 | 300.167 | 0.111 | 268.639 | 30.949 | 0.000 | 301.226 | 0.001 |
| 146 | 100 | 1000000 | 1000 | 5 | 5000 | 18671 | 5.0 | 15.0 | 100 | 31.654 | 0.116 | 0.217 | 31.252 | 0.000 | 32.358 | 0.001 |
| 147 | 100 | 1000000 | 1000 | 5 | 5000 | 147599 | 5.0 | 15.0 | 100 | 39.630 | 0.201 | 0.095 | 39.195 | 0.000 | 41.305 | 0.002 |
| 148 | 100 | 1000000 | 1000 | 5 | 5000 | 5249 | 8.4 | 25.2 | 100 | 30.532 | 0.111 | 0.098 | 30.259 | 0.000 | 31.225 | 0.001 |
| 149 | 100 | 1000000 | 1000 | 5 | 5000 | 18671 | 8.4 | 25.2 | 100 | 31.153 | 0.117 | 0.095 | 30.877 | 0.000 | 31.854 | 0.001 |
| 150 | 100 | 1000000 | 1000 | 5 | 5000 | 147599 | 8.4 | 25.2 | 100 | 40.217 | 0.195 | 0.098 | 39.773 | 0.000 | 41.892 | 0.002 |
| 151 | 100 | 1000000 | 1000 | 5 | 5000 | 5249 | 15.0 | 45.0 | 100 | 30.304 | 0.115 | 0.094 | 30.038 | 0.000 | 31.090 | 0.001 |
| 152 | 100 | 1000000 | 1000 | 5 | 5000 | 18671 | 15.0 | 45.0 | 100 | 31.646 | 0.117 | 0.099 | 31.359 | 0.000 | 32.328 | 0.001 |
| 153 | 100 | 1000000 | 1000 | 5 | 5000 | 147599 | 15.0 | 45.0 | 100 | 39.310 | 0.200 | 0.094 | 38.873 | 0.000 | 40.100 | 0.002 |
| 154 | 100 | 1000000 | 1000 | 50 | 50000 | 5249 | 5.0 | 15.0 | 100 | 30.614 | 0.114 | 0.097 | 30.338 | 0.000 | 31.392 | 0.001 |
| 155 | 100 | 1000000 | 1000 | 50 | 50000 | 18671 | 5.0 | 15.0 | 100 | 31.581 | 0.116 | 0.251 | 31.151 | 0.000 | 32.277 | 0.001 |
| 156 | 100 | 1000000 | 1000 | 50 | 50000 | 147599 | 5.0 | 15.0 | 100 | 39.702 | 0.202 | 0.102 | 39.231 | 0.000 | 40.553 | 0.002 |
| 157 | 100 | 1000000 | 1000 | 50 | 50000 | 5249 | 8.4 | 25.2 | 100 | 30.880 | 0.113 | 0.096 | 30.612 | 0.000 | 31.577 | 0.001 |
| 158 | 100 | 1000000 | 1000 | 50 | 50000 | 18671 | 8.4 | 25.2 | 100 | 31.680 | 0.118 | 0.191 | 31.304 | 0.000 | 32.375 | 0.001 |
| 159 | 100 | 1000000 | 1000 | 50 | 50000 | 147599 | 8.4 | 25.2 | 100 | 40.186 | 0.198 | 0.096 | 39.241 | 0.000 | 40.982 | 0.002 |
| 160 | 100 | 1000000 | 1000 | 50 | 50000 | 5249 | 15.0 | 45.0 | 100 | 30.378 | 0.111 | 0.097 | 30.102 | 0.000 | 31.062 | 0.001 |

*Continued on next page*

(continued from previous page)

| Run | $N$ | ECM | CELL | FOCAD/CELL | FOCAD | FNODE | $R_c$ | $R_s$ | Steps | $t_{init}$ | $t_{sim}$ | $t_{RTC}$ | $t_{initfn}$ | $t_{exit}$ | $t_{total}$ | $t_{step}$ |
|---|---|---|---|---|---|---|---|---|---|---|---|---|---|---|---|---|
| 161 | 100 | 1000000 | 1000 | 50 | 50000 | 18671 | 15.0 | 45.0 | 100 | 31.238 | 0.118 | 0.095 | 30.965 | 0.000 | 31.942 | 0.001 |
| 162 | 100 | 1000000 | 1000 | 50 | 50000 | 147599 | 15.0 | 45.0 | 100 | 39.367 | 0.193 | 0.098 | 38.924 | 0.000 | 40.221 | 0.002 |
| 163 | 100 | 1000000 | 10000 | 5 | 50000 | 5249 | 5.0 | 15.0 | 100 | 31.255 | 0.113 | 0.104 | 30.980 | 0.000 | 31.950 | 0.001 |
| 164 | 100 | 1000000 | 10000 | 5 | 50000 | 18671 | 5.0 | 15.0 | 100 | 31.009 | 0.114 | 0.099 | 30.728 | 0.000 | 31.689 | 0.001 |
| 165 | 100 | 1000000 | 10000 | 5 | 50000 | 147599 | 5.0 | 15.0 | 100 | 39.693 | 0.196 | 0.094 | 39.221 | 0.000 | 40.482 | 0.002 |
| 166 | 100 | 1000000 | 10000 | 5 | 50000 | 5249 | 8.4 | 25.2 | 100 | 30.029 | 0.112 | 0.097 | 29.757 | 0.000 | 30.727 | 0.001 |
| 167 | 100 | 1000000 | 10000 | 5 | 50000 | 18671 | 8.4 | 25.2 | 100 | 31.602 | 0.115 | 0.097 | 31.324 | 0.000 | 32.309 | 0.001 |
| 168 | 100 | 1000000 | 10000 | 5 | 50000 | 147599 | 8.4 | 25.2 | 100 | 39.834 | 0.198 | 0.097 | 39.389 | 0.000 | 40.611 | 0.002 |
| 169 | 100 | 1000000 | 10000 | 5 | 50000 | 5249 | 15.0 | 45.0 | 100 | 30.908 | 0.117 | 0.095 | 30.638 | 0.000 | 31.620 | 0.001 |
| 170 | 100 | 1000000 | 10000 | 5 | 50000 | 18671 | 15.0 | 45.0 | 100 | 31.486 | 0.119 | 0.227 | 30.994 | 0.000 | 32.255 | 0.001 |
| 171 | 100 | 1000000 | 10000 | 5 | 50000 | 147599 | 15.0 | 45.0 | 100 | 40.361 | 0.196 | 0.096 | 39.592 | 0.000 | 41.228 | 0.002 |
| 172 | 100 | 1000000 | 10000 | 50 | 500000 | 5249 | 5.0 | 15.0 | 100 | 30.136 | 0.111 | 0.098 | 29.865 | 0.000 | 30.892 | 0.001 |
| 173 | 100 | 1000000 | 10000 | 50 | 500000 | 18671 | 5.0 | 15.0 | 100 | 31.269 | 0.118 | 0.096 | 30.995 | 0.000 | 31.972 | 0.001 |
| 174 | 100 | 1000000 | 10000 | 50 | 500000 | 147599 | 5.0 | 15.0 | 100 | 39.401 | 0.200 | 0.097 | 38.955 | 0.000 | 40.181 | 0.002 |
| 175 | 100 | 1000000 | 10000 | 50 | 500000 | 5249 | 8.4 | 25.2 | 100 | 30.627 | 0.115 | 0.096 | 30.359 | 0.000 | 31.335 | 0.001 |
| 176 | 100 | 1000000 | 10000 | 50 | 500000 | 18671 | 8.4 | 25.2 | 100 | 30.925 | 0.113 | 0.099 | 30.641 | 0.000 | 31.625 | 0.001 |
| 177 | 100 | 1000000 | 10000 | 50 | 500000 | 147599 | 8.4 | 25.2 | 100 | 39.392 | 0.196 | 0.096 | 38.952 | 0.000 | 40.182 | 0.002 |
| 178 | 100 | 1000000 | 10000 | 50 | 500000 | 5249 | 15.0 | 45.0 | 100 | 30.893 | 0.114 | 0.098 | 30.619 | 0.000 | 31.575 | 0.001 |
| 179 | 100 | 1000000 | 10000 | 50 | 500000 | 18671 | 15.0 | 45.0 | 100 | 31.051 | 0.117 | 0.094 | 30.781 | 0.000 | 31.751 | 0.001 |
| 180 | 100 | 1000000 | 10000 | 50 | 500000 | 147599 | 15.0 | 45.0 | 100 | 39.480 | 0.194 | 0.097 | 39.033 | 0.000 | 40.249 | 0.002 |
| 181 | 100 | 1000000 | 100000 | 5 | 500000 | 5249 | 5.0 | 15.0 | 100 | 30.363 | 0.113 | 0.095 | 30.094 | 0.000 | 31.091 | 0.001 |
| 182 | 100 | 1000000 | 100000 | 5 | 500000 | 18671 | 5.0 | 15.0 | 100 | 31.066 | 0.119 | 0.103 | 30.785 | 0.000 | 32.688 | 0.001 |
| 183 | 100 | 1000000 | 100000 | 5 | 500000 | 147599 | 5.0 | 15.0 | 100 | 39.644 | 0.196 | 0.093 | 39.192 | 0.000 | 40.522 | 0.002 |
| 184 | 100 | 1000000 | 100000 | 5 | 500000 | 5249 | 8.4 | 25.2 | 100 | 30.598 | 0.114 | 0.098 | 30.325 | 0.000 | 32.182 | 0.001 |
| 185 | 100 | 1000000 | 100000 | 5 | 500000 | 18671 | 8.4 | 25.2 | 100 | 31.366 | 0.118 | 0.095 | 31.092 | 0.000 | 32.089 | 0.001 |
| 186 | 100 | 1000000 | 100000 | 5 | 500000 | 147599 | 8.4 | 25.2 | 100 | 39.535 | 0.198 | 0.299 | 38.776 | 0.000 | 40.379 | 0.002 |
| 187 | 100 | 1000000 | 100000 | 5 | 500000 | 5249 | 15.0 | 45.0 | 100 | 30.328 | 0.114 | 0.100 | 30.059 | 0.000 | 31.074 | 0.001 |
| 188 | 100 | 1000000 | 100000 | 5 | 500000 | 18671 | 15.0 | 45.0 | 100 | 31.282 | 0.118 | 0.099 | 30.987 | 0.000 | 31.983 | 0.001 |
| 189 | 100 | 1000000 | 100000 | 5 | 500000 | 147599 | 15.0 | 45.0 | 100 | 39.347 | 0.197 | 0.096 | 38.905 | 0.000 | 40.137 | 0.002 |
| 190 | 100 | 1000000 | 100000 | 50 | 5000000 | 5249 | 5.0 | 15.0 | 100 | 30.619 | 0.114 | 0.097 | 30.348 | 0.000 | 31.311 | 0.001 |
| 191 | 100 | 1000000 | 100000 | 50 | 5000000 | 18671 | 5.0 | 15.0 | 100 | 31.671 | 0.116 | 0.169 | 31.238 | 0.000 | 32.381 | 0.001 |
| 192 | 100 | 1000000 | 100000 | 50 | 5000000 | 147599 | 5.0 | 15.0 | 100 | 39.481 | 0.200 | 0.096 | 39.038 | 0.000 | 40.262 | 0.002 |
| 193 | 100 | 1000000 | 100000 | 50 | 5000000 | 5249 | 8.4 | 25.2 | 100 | 31.269 | 0.114 | 0.100 | 30.989 | 0.000 | 31.968 | 0.001 |
| 194 | 100 | 1000000 | 100000 | 50 | 5000000 | 18671 | 8.4 | 25.2 | 100 | 30.801 | 0.119 | 0.099 | 30.522 | 0.000 | 31.487 | 0.001 |
| 195 | 100 | 1000000 | 100000 | 50 | 5000000 | 147599 | 8.4 | 25.2 | 100 | 39.475 | 0.199 | 0.095 | 39.038 | 0.000 | 40.264 | 0.002 |
| 196 | 100 | 1000000 | 100000 | 50 | 5000000 | 5249 | 15.0 | 45.0 | 100 | 30.666 | 0.113 | 0.098 | 30.391 | 0.000 | 31.371 | 0.001 |
| 197 | 100 | 1000000 | 100000 | 50 | 5000000 | 18671 | 15.0 | 45.0 | 100 | 30.712 | 0.120 | 0.094 | 30.438 | 0.000 | 31.428 | 0.001 |
| 198 | 100 | 1000000 | 100000 | 50 | 5000000 | 147599 | 15.0 | 45.0 | 100 | 39.183 | 0.197 | 0.100 | 38.724 | 0.000 | 39.955 | 0.002 |
| 199 | 100 | 1000000 | 1000000 | 5 | 5000000 | 5249 | 5.0 | 15.0 | 100 | 30.274 | 0.111 | 0.096 | 30.007 | 0.000 | 31.885 | 0.001 |
| 200 | 100 | 1000000 | 1000000 | 5 | 5000000 | 18671 | 5.0 | 15.0 | 100 | 31.024 | 0.116 | 0.097 | 30.746 | 0.000 | 31.794 | 0.001 |
| 201 | 100 | 1000000 | 1000000 | 5 | 5000000 | 147599 | 5.0 | 15.0 | 100 | 39.727 | 0.196 | 0.093 | 39.295 | 0.000 | 40.505 | 0.002 |
| 202 | 100 | 1000000 | 1000000 | 5 | 5000000 | 5249 | 8.4 | 25.2 | 100 | 30.446 | 0.112 | 0.098 | 30.173 | 0.000 | 31.139 | 0.001 |
| 203 | 100 | 1000000 | 1000000 | 5 | 5000000 | 18671 | 8.4 | 25.2 | 100 | 31.585 | 0.116 | 0.094 | 31.315 | 0.000 | 32.343 | 0.001 |
| 204 | 100 | 1000000 | 1000000 | 5 | 5000000 | 147599 | 8.4 | 25.2 | 100 | 39.523 | 0.197 | 0.098 | 39.079 | 0.000 | 41.279 | 0.002 |
| 205 | 100 | 1000000 | 1000000 | 5 | 5000000 | 5249 | 15.0 | 45.0 | 100 | 30.288 | 0.115 | 0.100 | 30.016 | 0.000 | 31.890 | 0.001 |
| 206 | 100 | 1000000 | 1000000 | 5 | 5000000 | 18671 | 15.0 | 45.0 | 100 | 31.291 | 0.115 | 0.099 | 31.009 | 0.000 | 32.021 | 0.001 |
| 207 | 100 | 1000000 | 1000000 | 5 | 5000000 | 147599 | 15.0 | 45.0 | 100 | 40.007 | 0.197 | 0.094 | 39.564 | 0.000 | 40.796 | 0.002 |
| 208 | 100 | 1000000 | 1000000 | 50 | 50000000 | 5249 | 5.0 | 15.0 | 100 | 30.979 | 0.115 | 0.097 | 30.709 | 0.000 | 31.659 | 0.001 |
| 209 | 100 | 1000000 | 1000000 | 50 | 50000000 | 18671 | 5.0 | 15.0 | 100 | 31.297 | 0.115 | 0.093 | 31.023 | 0.000 | 31.998 | 0.001 |
| 210 | 100 | 1000000 | 1000000 | 50 | 50000000 | 147599 | 5.0 | 15.0 | 100 | 40.050 | 0.201 | 0.097 | 39.608 | 0.000 | 40.823 | 0.002 |
| 211 | 100 | 1000000 | 1000000 | 50 | 50000000 | 5249 | 8.4 | 25.2 | 100 | 30.760 | 0.111 | 0.097 | 30.495 | 0.000 | 31.461 | 0.001 |
| 212 | 100 | 1000000 | 1000000 | 50 | 50000000 | 18671 | 8.4 | 25.2 | 100 | 31.333 | 0.119 | 0.099 | 31.050 | 0.000 | 32.073 | 0.001 |
| 213 | 100 | 1000000 | 1000000 | 50 | 50000000 | 147599 | 8.4 | 25.2 | 100 | 39.251 | 0.196 | 0.095 | 38.806 | 0.000 | 40.036 | 0.002 |
| 214 | 100 | 1000000 | 1000000 | 50 | 50000000 | 5249 | 15.0 | 45.0 | 100 | 31.198 | 0.111 | 0.098 | 30.917 | 0.000 | 31.926 | 0.001 |

Continued on next page

(continued from previous page)

| Run | $N$ | ECM | CELL | FOCAD/CELL | FOCAD | FNODE | $R_c$ | $R_s$ | Steps | $t_{\text{init}}$ | $t_{\text{sim}}$ | $t_{\text{RTC}}$ | $t_{\text{initfn}}$ | $t_{\text{exit}}$ | $t_{\text{total}}$ | $t_{\text{step}}$ |
|---|---|---|---|---|---|---|---|---|---|---|---|---|---|---|---|---|
| 215 | 100 | 1000000 | 1000000 | 50 | 50000000 | 18671 | 15.0 | 45.0 | 100 | 31.077 | 0.114 | 0.095 | 30.803 | 0.000 | 31.765 | 0.001 |
| 216 | 100 | 1000000 | 1000000 | 50 | 50000000 | 147599 | 15.0 | 45.0 | 100 | 39.834 | 0.192 | 0.097 | 39.391 | 0.000 | 40.594 | 0.002 |

Table 7: H100-NVL raw benchmark results. Times are in seconds. Compilation time ($t_{\mathrm{RTC}}$) is subtracted from $t_{\mathrm{init}}$ to compute actual initialization time.

| Run | $N$ | ECM | CELL | FOCAD/CELL | FOCAD | FNODE | $R_c$ | $R_s$ | Steps | $t_{\mathrm{init}}$ | $t_{\mathrm{sim}}$ | $t_{\mathrm{RTC}}$ | $t_{\mathrm{initfn}}$ | $t_{\mathrm{exit}}$ | $t_{\mathrm{total}}$ | $t_{\mathrm{step}}$ |
|---|---|---|---|---|---|---|---|---|---|---|---|---|---|---|---|---|
| 1 | 21 | 9261 | 1000 | 5 | 5000 | 5249 | 5.0 | 15.0 | 100 | 194.459 | 0.066 | 193.723 | 0.566 | 0.000 | 195.200 | 0.001 |
| 2 | 21 | 9261 | 1000 | 5 | 5000 | 18671 | 5.0 | 15.0 | 100 | 1.623 | 0.068 | 0.105 | 1.341 | 0.000 | 2.253 | 0.001 |
| 3 | 21 | 9261 | 1000 | 5 | 5000 | 147599 | 5.0 | 15.0 | 100 | 9.053 | 0.117 | 0.103 | 8.612 | 0.000 | 10.632 | 0.001 |
| 4 | 21 | 9261 | 1000 | 5 | 5000 | 5249 | 8.4 | 25.2 | 100 | 0.820 | 0.066 | 0.104 | 0.548 | 0.000 | 1.449 | 0.001 |
| 5 | 21 | 9261 | 1000 | 5 | 5000 | 18671 | 8.4 | 25.2 | 100 | 1.595 | 0.068 | 0.103 | 1.316 | 0.000 | 2.221 | 0.001 |
| 6 | 21 | 9261 | 1000 | 5 | 5000 | 147599 | 8.4 | 25.2 | 100 | 8.970 | 0.117 | 0.103 | 8.546 | 0.000 | 9.643 | 0.001 |
| 7 | 21 | 9261 | 1000 | 5 | 5000 | 5249 | 15.0 | 45.0 | 100 | 0.826 | 0.065 | 0.103 | 0.553 | 0.000 | 1.451 | 0.001 |
| 8 | 21 | 9261 | 1000 | 5 | 5000 | 18671 | 15.0 | 45.0 | 100 | 1.594 | 0.068 | 0.102 | 1.318 | 0.000 | 2.221 | 0.001 |
| 9 | 21 | 9261 | 1000 | 5 | 5000 | 147599 | 15.0 | 45.0 | 100 | 9.065 | 0.117 | 0.103 | 8.638 | 0.000 | 9.738 | 0.001 |
| 10 | 21 | 9261 | 1000 | 50 | 50000 | 5249 | 5.0 | 15.0 | 100 | 0.830 | 0.066 | 0.103 | 0.558 | 0.000 | 1.454 | 0.001 |
| 11 | 21 | 9261 | 1000 | 50 | 50000 | 18671 | 5.0 | 15.0 | 100 | 1.587 | 0.068 | 0.103 | 1.311 | 0.000 | 2.215 | 0.001 |
| 12 | 21 | 9261 | 1000 | 50 | 50000 | 147599 | 5.0 | 15.0 | 100 | 8.879 | 0.117 | 0.102 | 8.453 | 0.000 | 9.569 | 0.001 |
| 13 | 21 | 9261 | 1000 | 50 | 50000 | 5249 | 8.4 | 25.2 | 100 | 0.834 | 0.065 | 0.104 | 0.559 | 0.000 | 1.472 | 0.001 |
| 14 | 21 | 9261 | 1000 | 50 | 50000 | 18671 | 8.4 | 25.2 | 100 | 1.657 | 0.068 | 0.102 | 1.379 | 0.000 | 2.282 | 0.001 |
| 15 | 21 | 9261 | 1000 | 50 | 50000 | 147599 | 8.4 | 25.2 | 100 | 8.968 | 0.117 | 0.103 | 8.539 | 0.000 | 9.646 | 0.001 |
| 16 | 21 | 9261 | 1000 | 50 | 50000 | 5249 | 15.0 | 45.0 | 100 | 0.829 | 0.065 | 0.104 | 0.556 | 0.000 | 1.452 | 0.001 |
| 17 | 21 | 9261 | 1000 | 50 | 50000 | 18671 | 15.0 | 45.0 | 100 | 1.582 | 0.068 | 0.103 | 1.305 | 0.000 | 2.211 | 0.001 |
| 18 | 21 | 9261 | 1000 | 50 | 50000 | 147599 | 15.0 | 45.0 | 100 | 9.007 | 0.117 | 0.102 | 8.581 | 0.000 | 9.680 | 0.001 |
| 19 | 21 | 9261 | 10000 | 5 | 50000 | 5249 | 5.0 | 15.0 | 100 | 0.817 | 0.065 | 0.103 | 0.544 | 0.000 | 1.435 | 0.001 |
| 20 | 21 | 9261 | 10000 | 5 | 50000 | 18671 | 5.0 | 15.0 | 100 | 1.594 | 0.068 | 0.102 | 1.318 | 0.000 | 2.228 | 0.001 |
| 21 | 21 | 9261 | 10000 | 5 | 50000 | 147599 | 5.0 | 15.0 | 100 | 9.136 | 0.117 | 0.102 | 8.712 | 0.000 | 10.732 | 0.001 |
| 22 | 21 | 9261 | 10000 | 5 | 50000 | 5249 | 8.4 | 25.2 | 100 | 0.828 | 0.065 | 0.103 | 0.557 | 0.000 | 1.456 | 0.001 |
| 23 | 21 | 9261 | 10000 | 5 | 50000 | 18671 | 8.4 | 25.2 | 100 | 1.596 | 0.068 | 0.102 | 1.320 | 0.000 | 2.225 | 0.001 |
| 24 | 21 | 9261 | 10000 | 5 | 50000 | 147599 | 8.4 | 25.2 | 100 | 9.063 | 0.117 | 0.103 | 8.635 | 0.000 | 9.757 | 0.001 |
| 25 | 21 | 9261 | 10000 | 5 | 50000 | 5249 | 15.0 | 45.0 | 100 | 0.844 | 0.066 | 0.104 | 0.567 | 0.000 | 1.473 | 0.001 |
| 26 | 21 | 9261 | 10000 | 5 | 50000 | 18671 | 15.0 | 45.0 | 100 | 1.587 | 0.068 | 0.103 | 1.311 | 0.000 | 2.210 | 0.001 |
| 27 | 21 | 9261 | 10000 | 5 | 50000 | 147599 | 15.0 | 45.0 | 100 | 8.950 | 0.117 | 0.103 | 8.525 | 0.000 | 9.630 | 0.001 |
| 28 | 21 | 9261 | 10000 | 50 | 500000 | 5249 | 5.0 | 15.0 | 100 | 0.831 | 0.066 | 0.104 | 0.558 | 0.000 | 1.455 | 0.001 |
| 29 | 21 | 9261 | 10000 | 50 | 500000 | 18671 | 5.0 | 15.0 | 100 | 1.601 | 0.068 | 0.103 | 1.322 | 0.000 | 2.241 | 0.001 |
| 30 | 21 | 9261 | 10000 | 50 | 500000 | 147599 | 5.0 | 15.0 | 100 | 9.015 | 0.117 | 0.102 | 8.592 | 0.000 | 9.693 | 0.001 |
| 31 | 21 | 9261 | 10000 | 50 | 500000 | 5249 | 8.4 | 25.2 | 100 | 0.830 | 0.066 | 0.103 | 0.558 | 0.000 | 1.457 | 0.001 |
| 32 | 21 | 9261 | 10000 | 50 | 500000 | 18671 | 8.4 | 25.2 | 100 | 1.632 | 0.068 | 0.103 | 1.351 | 0.000 | 2.259 | 0.001 |
| 33 | 21 | 9261 | 10000 | 50 | 500000 | 147599 | 8.4 | 25.2 | 100 | 9.009 | 0.117 | 0.103 | 8.581 | 0.000 | 9.689 | 0.001 |
| 34 | 21 | 9261 | 10000 | 50 | 500000 | 5249 | 15.0 | 45.0 | 100 | 0.830 | 0.065 | 0.103 | 0.558 | 0.000 | 1.448 | 0.001 |
| 35 | 21 | 9261 | 10000 | 50 | 500000 | 18671 | 15.0 | 45.0 | 100 | 1.581 | 0.068 | 0.104 | 1.302 | 0.000 | 2.231 | 0.001 |
| 36 | 21 | 9261 | 10000 | 50 | 500000 | 147599 | 15.0 | 45.0 | 100 | 8.875 | 0.117 | 0.103 | 8.447 | 0.000 | 9.574 | 0.001 |
| 37 | 21 | 9261 | 100000 | 5 | 500000 | 5249 | 5.0 | 15.0 | 100 | 0.831 | 0.065 | 0.105 | 0.556 | 0.000 | 1.454 | 0.001 |
| 38 | 21 | 9261 | 100000 | 5 | 500000 | 18671 | 5.0 | 15.0 | 100 | 1.587 | 0.068 | 0.104 | 1.307 | 0.000 | 2.219 | 0.001 |
| 39 | 21 | 9261 | 100000 | 5 | 500000 | 147599 | 5.0 | 15.0 | 100 | 8.997 | 0.117 | 0.103 | 8.569 | 0.000 | 9.686 | 0.001 |
| 40 | 21 | 9261 | 100000 | 5 | 500000 | 5249 | 8.4 | 25.2 | 100 | 0.829 | 0.065 | 0.106 | 0.550 | 0.000 | 1.460 | 0.001 |
| 41 | 21 | 9261 | 100000 | 5 | 500000 | 18671 | 8.4 | 25.2 | 100 | 1.595 | 0.068 | 0.104 | 1.314 | 0.000 | 3.125 | 0.001 |
| 42 | 21 | 9261 | 100000 | 5 | 500000 | 147599 | 8.4 | 25.2 | 100 | 8.919 | 0.117 | 0.104 | 8.489 | 0.000 | 10.499 | 0.001 |
| 43 | 21 | 9261 | 100000 | 5 | 500000 | 5249 | 15.0 | 45.0 | 100 | 0.829 | 0.065 | 0.107 | 0.552 | 0.000 | 1.454 | 0.001 |
| 44 | 21 | 9261 | 100000 | 5 | 500000 | 18671 | 15.0 | 45.0 | 100 | 1.600 | 0.068 | 0.104 | 1.319 | 0.000 | 2.225 | 0.001 |
| 45 | 21 | 9261 | 100000 | 5 | 500000 | 147599 | 15.0 | 45.0 | 100 | 9.015 | 0.117 | 0.103 | 8.589 | 0.000 | 9.690 | 0.001 |
| 46 | 21 | 9261 | 100000 | 50 | 5000000 | 5249 | 5.0 | 15.0 | 100 | 0.832 | 0.066 | 0.105 | 0.558 | 0.000 | 1.455 | 0.001 |
| 47 | 21 | 9261 | 100000 | 50 | 5000000 | 18671 | 5.0 | 15.0 | 100 | 1.588 | 0.069 | 0.103 | 1.310 | 0.000 | 3.120 | 0.001 |
| 48 | 21 | 9261 | 100000 | 50 | 5000000 | 147599 | 5.0 | 15.0 | 100 | 9.057 | 0.117 | 0.104 | 8.625 | 0.000 | 9.757 | 0.001 |
| 49 | 21 | 9261 | 100000 | 50 | 5000000 | 5249 | 8.4 | 25.2 | 100 | 0.825 | 0.065 | 0.106 | 0.549 | 0.000 | 1.456 | 0.001 |
| 50 | 21 | 9261 | 100000 | 50 | 5000000 | 18671 | 8.4 | 25.2 | 100 | 1.611 | 0.068 | 0.104 | 1.331 | 0.000 | 2.233 | 0.001 |
| 51 | 21 | 9261 | 100000 | 50 | 5000000 | 147599 | 8.4 | 25.2 | 100 | 9.102 | 0.117 | 0.104 | 8.674 | 0.000 | 9.777 | 0.001 |
| 52 | 21 | 9261 | 100000 | 50 | 5000000 | 5249 | 15.0 | 45.0 | 100 | 0.829 | 0.066 | 0.105 | 0.555 | 0.000 | 1.449 | 0.001 |

*Continued on next page*

(continued from previous page)

| Run | $N$ | ECM | CELL | FOCAD/CELL | FOCAD | FNODE | $R_c$ | $R_s$ | Steps | $t_{init}$ | $t_{sim}$ | $t_{RTC}$ | $t_{initfn}$ | $t_{exit}$ | $t_{total}$ | $t_{step}$ |
|---|---|---|---|---|---|---|---|---|---|---|---|---|---|---|---|---|
| 53 | 21 | 9261 | 100000 | 50 | 5000000 | 18671 | 15.0 | 45.0 | 100 | 1.627 | 0.068 | 0.104 | 1.348 | 0.000 | 2.253 | 0.001 |
| 54 | 21 | 9261 | 100000 | 50 | 5000000 | 147599 | 15.0 | 45.0 | 100 | 8.918 | 0.116 | 0.104 | 8.488 | 0.000 | 9.597 | 0.001 |
| 55 | 21 | 9261 | 1000000 | 5 | 5000000 | 5249 | 5.0 | 15.0 | 100 | 0.832 | 0.065 | 0.105 | 0.558 | 0.000 | 1.457 | 0.001 |
| 56 | 21 | 9261 | 1000000 | 5 | 5000000 | 18671 | 5.0 | 15.0 | 100 | 1.600 | 0.069 | 0.104 | 1.321 | 0.000 | 2.229 | 0.001 |
| 57 | 21 | 9261 | 1000000 | 5 | 5000000 | 147599 | 5.0 | 15.0 | 100 | 9.048 | 0.117 | 0.104 | 8.619 | 0.000 | 9.737 | 0.001 |
| 58 | 21 | 9261 | 1000000 | 5 | 5000000 | 5249 | 8.4 | 25.2 | 100 | 0.820 | 0.065 | 0.105 | 0.546 | 0.000 | 1.442 | 0.001 |
| 59 | 21 | 9261 | 1000000 | 5 | 5000000 | 18671 | 8.4 | 25.2 | 100 | 1.599 | 0.068 | 0.104 | 1.318 | 0.000 | 2.268 | 0.001 |
| 60 | 21 | 9261 | 1000000 | 5 | 5000000 | 147599 | 8.4 | 25.2 | 100 | 8.995 | 0.117 | 0.103 | 8.568 | 0.000 | 9.704 | 0.001 |
| 61 | 21 | 9261 | 1000000 | 5 | 5000000 | 5249 | 15.0 | 45.0 | 100 | 0.834 | 0.065 | 0.106 | 0.558 | 0.000 | 1.466 | 0.001 |
| 62 | 21 | 9261 | 1000000 | 5 | 5000000 | 18671 | 15.0 | 45.0 | 100 | 1.593 | 0.068 | 0.107 | 1.312 | 0.000 | 2.224 | 0.001 |
| 63 | 21 | 9261 | 1000000 | 5 | 5000000 | 147599 | 15.0 | 45.0 | 100 | 9.039 | 0.117 | 0.104 | 8.612 | 0.000 | 9.726 | 0.001 |
| 64 | 21 | 9261 | 1000000 | 50 | 50000000 | 5249 | 5.0 | 15.0 | 100 | 0.833 | 0.066 | 0.105 | 0.559 | 0.000 | 1.461 | 0.001 |
| 65 | 21 | 9261 | 1000000 | 50 | 50000000 | 18671 | 5.0 | 15.0 | 100 | 1.671 | 0.068 | 0.104 | 1.392 | 0.000 | 2.292 | 0.001 |
| 66 | 21 | 9261 | 1000000 | 50 | 50000000 | 147599 | 5.0 | 15.0 | 100 | 8.985 | 0.117 | 0.104 | 8.553 | 0.000 | 9.673 | 0.001 |
| 67 | 21 | 9261 | 1000000 | 50 | 50000000 | 5249 | 8.4 | 25.2 | 100 | 0.830 | 0.066 | 0.107 | 0.554 | 0.000 | 1.453 | 0.001 |
| 68 | 21 | 9261 | 1000000 | 50 | 50000000 | 18671 | 8.4 | 25.2 | 100 | 1.590 | 0.068 | 0.104 | 1.312 | 0.000 | 2.211 | 0.001 |
| 69 | 21 | 9261 | 1000000 | 50 | 50000000 | 147599 | 8.4 | 25.2 | 100 | 8.995 | 0.117 | 0.104 | 8.565 | 0.000 | 9.674 | 0.001 |
| 70 | 21 | 9261 | 1000000 | 50 | 50000000 | 5249 | 15.0 | 45.0 | 100 | 0.825 | 0.065 | 0.104 | 0.551 | 0.000 | 1.452 | 0.001 |
| 71 | 21 | 9261 | 1000000 | 50 | 50000000 | 18671 | 15.0 | 45.0 | 100 | 1.599 | 0.068 | 0.104 | 1.320 | 0.000 | 2.223 | 0.001 |
| 72 | 21 | 9261 | 1000000 | 50 | 50000000 | 147599 | 15.0 | 45.0 | 100 | 8.923 | 0.117 | 0.104 | 8.494 | 0.000 | 9.623 | 0.001 |
| 73 | 46 | 97336 | 1000 | 5 | 5000 | 5249 | 5.0 | 15.0 | 100 | 196.582 | 0.066 | 193.416 | 2.996 | 0.000 | 197.225 | 0.001 |
| 74 | 46 | 97336 | 1000 | 5 | 5000 | 18671 | 5.0 | 15.0 | 100 | 4.049 | 0.068 | 0.106 | 3.764 | 0.000 | 4.674 | 0.001 |
| 75 | 46 | 97336 | 1000 | 5 | 5000 | 147599 | 5.0 | 15.0 | 100 | 11.389 | 0.116 | 0.104 | 10.962 | 0.000 | 12.070 | 0.001 |
| 76 | 46 | 97336 | 1000 | 5 | 5000 | 5249 | 8.4 | 25.2 | 100 | 3.277 | 0.066 | 0.105 | 3.003 | 0.000 | 3.911 | 0.001 |
| 77 | 46 | 97336 | 1000 | 5 | 5000 | 18671 | 8.4 | 25.2 | 100 | 4.019 | 0.068 | 0.102 | 3.744 | 0.000 | 4.649 | 0.001 |
| 78 | 46 | 97336 | 1000 | 5 | 5000 | 147599 | 8.4 | 25.2 | 100 | 11.344 | 0.117 | 0.102 | 10.919 | 0.000 | 12.924 | 0.001 |
| 79 | 46 | 97336 | 1000 | 5 | 5000 | 5249 | 15.0 | 45.0 | 100 | 3.278 | 0.066 | 0.103 | 3.009 | 0.000 | 3.901 | 0.001 |
| 80 | 46 | 97336 | 1000 | 5 | 5000 | 18671 | 15.0 | 45.0 | 100 | 4.050 | 0.068 | 0.102 | 3.774 | 0.000 | 4.671 | 0.001 |
| 81 | 46 | 97336 | 1000 | 5 | 5000 | 147599 | 15.0 | 45.0 | 100 | 11.431 | 0.116 | 0.103 | 11.006 | 0.000 | 12.125 | 0.001 |
| 82 | 46 | 97336 | 1000 | 50 | 50000 | 5249 | 5.0 | 15.0 | 100 | 3.330 | 0.066 | 0.104 | 3.058 | 0.000 | 3.953 | 0.001 |
| 83 | 46 | 97336 | 1000 | 50 | 50000 | 18671 | 5.0 | 15.0 | 100 | 4.014 | 0.068 | 0.103 | 3.737 | 0.000 | 4.647 | 0.001 |
| 84 | 46 | 97336 | 1000 | 50 | 50000 | 147599 | 5.0 | 15.0 | 100 | 11.479 | 0.117 | 0.102 | 11.056 | 0.000 | 13.061 | 0.001 |
| 85 | 46 | 97336 | 1000 | 50 | 50000 | 5249 | 8.4 | 25.2 | 100 | 3.315 | 0.066 | 0.103 | 3.043 | 0.000 | 3.974 | 0.001 |
| 86 | 46 | 97336 | 1000 | 50 | 50000 | 18671 | 8.4 | 25.2 | 100 | 4.072 | 0.068 | 0.103 | 3.795 | 0.000 | 4.701 | 0.001 |
| 87 | 46 | 97336 | 1000 | 50 | 50000 | 147599 | 8.4 | 25.2 | 100 | 11.387 | 0.117 | 0.102 | 10.964 | 0.000 | 12.074 | 0.001 |
| 88 | 46 | 97336 | 1000 | 50 | 50000 | 5249 | 15.0 | 45.0 | 100 | 3.314 | 0.066 | 0.148 | 2.997 | 0.000 | 3.953 | 0.001 |
| 89 | 46 | 97336 | 1000 | 50 | 50000 | 18671 | 15.0 | 45.0 | 100 | 4.240 | 0.068 | 0.102 | 3.962 | 0.000 | 4.879 | 0.001 |
| 90 | 46 | 97336 | 1000 | 50 | 50000 | 147599 | 15.0 | 45.0 | 100 | 11.518 | 0.117 | 0.103 | 11.092 | 0.000 | 12.216 | 0.001 |
| 91 | 46 | 97336 | 10000 | 5 | 50000 | 5249 | 5.0 | 15.0 | 100 | 3.251 | 0.066 | 0.104 | 2.980 | 0.000 | 4.777 | 0.001 |
| 92 | 46 | 97336 | 10000 | 5 | 50000 | 18671 | 5.0 | 15.0 | 100 | 4.084 | 0.068 | 0.103 | 3.804 | 0.000 | 5.191 | 0.001 |
| 93 | 46 | 97336 | 10000 | 5 | 50000 | 147599 | 5.0 | 15.0 | 100 | 11.564 | 0.117 | 0.103 | 11.139 | 0.000 | 12.238 | 0.001 |
| 94 | 46 | 97336 | 10000 | 5 | 50000 | 5249 | 8.4 | 25.2 | 100 | 3.313 | 0.067 | 0.103 | 3.043 | 0.000 | 3.933 | 0.001 |
| 95 | 46 | 97336 | 10000 | 5 | 50000 | 18671 | 8.4 | 25.2 | 100 | 4.031 | 0.068 | 0.103 | 3.752 | 0.000 | 4.664 | 0.001 |
| 96 | 46 | 97336 | 10000 | 5 | 50000 | 147599 | 8.4 | 25.2 | 100 | 11.398 | 0.116 | 0.102 | 10.972 | 0.000 | 12.091 | 0.001 |
| 97 | 46 | 97336 | 10000 | 5 | 50000 | 5249 | 15.0 | 45.0 | 100 | 3.281 | 0.066 | 0.104 | 3.009 | 0.000 | 3.899 | 0.001 |
| 98 | 46 | 97336 | 10000 | 5 | 50000 | 18671 | 15.0 | 45.0 | 100 | 4.060 | 0.068 | 0.103 | 3.783 | 0.000 | 4.709 | 0.001 |
| 99 | 46 | 97336 | 10000 | 5 | 50000 | 147599 | 15.0 | 45.0 | 100 | 11.563 | 0.117 | 0.103 | 11.136 | 0.000 | 12.258 | 0.001 |
| 100 | 46 | 97336 | 10000 | 50 | 500000 | 5249 | 5.0 | 15.0 | 100 | 3.242 | 0.065 | 0.104 | 2.969 | 0.000 | 4.444 | 0.001 |
| 101 | 46 | 97336 | 10000 | 50 | 500000 | 18671 | 5.0 | 15.0 | 100 | 4.042 | 0.068 | 0.103 | 3.763 | 0.000 | 4.673 | 0.001 |
| 102 | 46 | 97336 | 10000 | 50 | 500000 | 147599 | 5.0 | 15.0 | 100 | 11.874 | 0.116 | 0.103 | 11.449 | 0.000 | 12.546 | 0.001 |
| 103 | 46 | 97336 | 10000 | 50 | 500000 | 5249 | 8.4 | 25.2 | 100 | 3.299 | 0.066 | 0.104 | 3.024 | 0.000 | 3.940 | 0.001 |
| 104 | 46 | 97336 | 10000 | 50 | 500000 | 18671 | 8.4 | 25.2 | 100 | 4.044 | 0.068 | 0.102 | 3.767 | 0.000 | 4.681 | 0.001 |
| 105 | 46 | 97336 | 10000 | 50 | 500000 | 147599 | 8.4 | 25.2 | 100 | 11.457 | 0.117 | 0.104 | 11.030 | 0.000 | 12.151 | 0.001 |
| 106 | 46 | 97336 | 10000 | 50 | 500000 | 5249 | 15.0 | 45.0 | 100 | 3.338 | 0.067 | 0.103 | 3.067 | 0.000 | 3.975 | 0.001 |

Continued on next page

*(continued from previous page)*

| Run | $N$ | ECM | CELL | FOCAD/CELL | FOCAD | FNODE | $R_c$ | $R_s$ | Steps | $t_{init}$ | $t_{sim}$ | $t_{RTC}$ | $t_{initfn}$ | $t_{exit}$ | $t_{total}$ | $t_{step}$ |
|---|---|---|---|---|---|---|---|---|---|---|---|---|---|---|---|---|
| 107 | 46 | 97336 | 10000 | 50 | 500000 | 18671 | 15.0 | 45.0 | 100 | 4.019 | 0.068 | 0.103 | 3.739 | 0.000 | 4.645 | 0.001 |
| 108 | 46 | 97336 | 10000 | 50 | 500000 | 147599 | 15.0 | 45.0 | 100 | 11.443 | 0.117 | 0.103 | 11.018 | 0.000 | 13.027 | 0.001 |
| 109 | 46 | 97336 | 100000 | 5 | 500000 | 5249 | 5.0 | 15.0 | 100 | 3.222 | 0.066 | 0.105 | 2.949 | 0.000 | 4.451 | 0.001 |
| 110 | 46 | 97336 | 100000 | 5 | 500000 | 18671 | 5.0 | 15.0 | 100 | 4.077 | 0.068 | 0.103 | 3.797 | 0.000 | 4.703 | 0.001 |
| 111 | 46 | 97336 | 100000 | 5 | 500000 | 147599 | 5.0 | 15.0 | 100 | 11.623 | 0.117 | 0.102 | 11.199 | 0.000 | 12.311 | 0.001 |
| 112 | 46 | 97336 | 100000 | 5 | 500000 | 5249 | 8.4 | 25.2 | 100 | 3.257 | 0.066 | 0.105 | 2.984 | 0.000 | 3.880 | 0.001 |
| 113 | 46 | 97336 | 100000 | 5 | 500000 | 18671 | 8.4 | 25.2 | 100 | 3.981 | 0.069 | 0.102 | 3.705 | 0.000 | 4.601 | 0.001 |
| 114 | 46 | 97336 | 100000 | 5 | 500000 | 147599 | 8.4 | 25.2 | 100 | 11.414 | 0.116 | 0.103 | 10.986 | 0.000 | 12.107 | 0.001 |
| 115 | 46 | 97336 | 100000 | 5 | 500000 | 5249 | 15.0 | 45.0 | 100 | 3.303 | 0.066 | 0.103 | 3.032 | 0.000 | 3.930 | 0.001 |
| 116 | 46 | 97336 | 100000 | 5 | 500000 | 18671 | 15.0 | 45.0 | 100 | 4.114 | 0.068 | 0.104 | 3.835 | 0.000 | 4.740 | 0.001 |
| 117 | 46 | 97336 | 100000 | 5 | 500000 | 147599 | 15.0 | 45.0 | 100 | 11.443 | 0.117 | 0.103 | 11.014 | 0.000 | 12.154 | 0.001 |
| 118 | 46 | 97336 | 100000 | 50 | 5000000 | 5249 | 5.0 | 15.0 | 100 | 3.311 | 0.066 | 0.104 | 3.037 | 0.000 | 3.934 | 0.001 |
| 119 | 46 | 97336 | 100000 | 50 | 5000000 | 18671 | 5.0 | 15.0 | 100 | 4.046 | 0.068 | 0.103 | 3.768 | 0.000 | 4.679 | 0.001 |
| 120 | 46 | 97336 | 100000 | 50 | 5000000 | 147599 | 5.0 | 15.0 | 100 | 11.455 | 0.117 | 0.104 | 11.028 | 0.000 | 12.158 | 0.001 |
| 121 | 46 | 97336 | 100000 | 50 | 5000000 | 5249 | 8.4 | 25.2 | 100 | 3.318 | 0.066 | 0.104 | 3.044 | 0.000 | 3.937 | 0.001 |
| 122 | 46 | 97336 | 100000 | 50 | 5000000 | 18671 | 8.4 | 25.2 | 100 | 4.102 | 0.068 | 0.103 | 3.822 | 0.000 | 4.727 | 0.001 |
| 123 | 46 | 97336 | 100000 | 50 | 5000000 | 147599 | 8.4 | 25.2 | 100 | 11.400 | 0.117 | 0.103 | 10.974 | 0.000 | 12.086 | 0.001 |
| 124 | 46 | 97336 | 100000 | 50 | 5000000 | 5249 | 15.0 | 45.0 | 100 | 3.293 | 0.066 | 0.104 | 3.020 | 0.000 | 3.925 | 0.001 |
| 125 | 46 | 97336 | 100000 | 50 | 5000000 | 18671 | 15.0 | 45.0 | 100 | 3.986 | 0.069 | 0.103 | 3.706 | 0.000 | 4.615 | 0.001 |
| 126 | 46 | 97336 | 100000 | 50 | 5000000 | 147599 | 15.0 | 45.0 | 100 | 11.637 | 0.116 | 0.103 | 11.209 | 0.000 | 12.342 | 0.001 |
| 127 | 46 | 97336 | 1000000 | 5 | 5000000 | 5249 | 5.0 | 15.0 | 100 | 3.330 | 0.066 | 0.104 | 3.058 | 0.000 | 3.958 | 0.001 |
| 128 | 46 | 97336 | 1000000 | 5 | 5000000 | 18671 | 5.0 | 15.0 | 100 | 4.194 | 0.068 | 0.104 | 3.915 | 0.000 | 4.823 | 0.001 |
| 129 | 46 | 97336 | 1000000 | 5 | 5000000 | 147599 | 5.0 | 15.0 | 100 | 11.438 | 0.117 | 0.102 | 11.012 | 0.000 | 12.130 | 0.001 |
| 130 | 46 | 97336 | 1000000 | 5 | 5000000 | 5249 | 8.4 | 25.2 | 100 | 3.250 | 0.066 | 0.104 | 2.978 | 0.000 | 3.875 | 0.001 |
| 131 | 46 | 97336 | 1000000 | 5 | 5000000 | 18671 | 8.4 | 25.2 | 100 | 4.103 | 0.068 | 0.103 | 3.824 | 0.000 | 4.736 | 0.001 |
| 132 | 46 | 97336 | 1000000 | 5 | 5000000 | 147599 | 8.4 | 25.2 | 100 | 11.428 | 0.117 | 0.103 | 11.003 | 0.000 | 12.113 | 0.001 |
| 133 | 46 | 97336 | 1000000 | 5 | 5000000 | 5249 | 15.0 | 45.0 | 100 | 3.262 | 0.066 | 0.103 | 2.991 | 0.000 | 3.887 | 0.001 |
| 134 | 46 | 97336 | 1000000 | 5 | 5000000 | 18671 | 15.0 | 45.0 | 100 | 3.974 | 0.068 | 0.103 | 3.694 | 0.000 | 4.601 | 0.001 |
| 135 | 46 | 97336 | 1000000 | 5 | 5000000 | 147599 | 15.0 | 45.0 | 100 | 11.357 | 0.116 | 0.103 | 10.932 | 0.000 | 12.943 | 0.001 |
| 136 | 46 | 97336 | 1000000 | 50 | 50000000 | 5249 | 5.0 | 15.0 | 100 | 3.291 | 0.066 | 0.104 | 3.017 | 0.000 | 3.915 | 0.001 |
| 137 | 46 | 97336 | 1000000 | 50 | 50000000 | 18671 | 5.0 | 15.0 | 100 | 3.993 | 0.068 | 0.103 | 3.712 | 0.000 | 5.522 | 0.001 |
| 138 | 46 | 97336 | 1000000 | 50 | 50000000 | 147599 | 5.0 | 15.0 | 100 | 11.443 | 0.116 | 0.103 | 11.014 | 0.000 | 12.137 | 0.001 |
| 139 | 46 | 97336 | 1000000 | 50 | 50000000 | 5249 | 8.4 | 25.2 | 100 | 3.236 | 0.066 | 0.104 | 2.964 | 0.000 | 3.858 | 0.001 |
| 140 | 46 | 97336 | 1000000 | 50 | 50000000 | 18671 | 8.4 | 25.2 | 100 | 4.025 | 0.068 | 0.103 | 3.745 | 0.000 | 4.656 | 0.001 |
| 141 | 46 | 97336 | 1000000 | 50 | 50000000 | 147599 | 8.4 | 25.2 | 100 | 11.459 | 0.116 | 0.103 | 11.031 | 0.000 | 12.141 | 0.001 |
| 142 | 46 | 97336 | 1000000 | 50 | 50000000 | 5249 | 15.0 | 45.0 | 100 | 3.269 | 0.066 | 0.103 | 2.999 | 0.000 | 3.903 | 0.001 |
| 143 | 46 | 97336 | 1000000 | 50 | 50000000 | 18671 | 15.0 | 45.0 | 100 | 4.020 | 0.068 | 0.103 | 3.742 | 0.000 | 5.549 | 0.001 |
| 144 | 46 | 97336 | 1000000 | 50 | 50000000 | 147599 | 15.0 | 45.0 | 100 | 11.428 | 0.116 | 0.103 | 11.002 | 0.000 | 12.128 | 0.001 |
| 145 | 100 | 1000000 | 1000 | 5 | 5000 | 5249 | 5.0 | 15.0 | 100 | 221.873 | 0.066 | 193.367 | 28.336 | 0.000 | 222.527 | 0.001 |
| 146 | 100 | 1000000 | 1000 | 5 | 5000 | 18671 | 5.0 | 15.0 | 100 | 29.694 | 0.069 | 0.106 | 29.413 | 0.000 | 30.331 | 0.001 |
| 147 | 100 | 1000000 | 1000 | 5 | 5000 | 147599 | 5.0 | 15.0 | 100 | 36.521 | 0.117 | 0.103 | 36.095 | 0.000 | 37.233 | 0.001 |
| 148 | 100 | 1000000 | 1000 | 5 | 5000 | 5249 | 8.4 | 25.2 | 100 | 28.280 | 0.066 | 0.105 | 28.005 | 0.000 | 28.914 | 0.001 |
| 149 | 100 | 1000000 | 1000 | 5 | 5000 | 18671 | 8.4 | 25.2 | 100 | 29.073 | 0.069 | 0.103 | 28.794 | 0.000 | 30.615 | 0.001 |
| 150 | 100 | 1000000 | 1000 | 5 | 5000 | 147599 | 8.4 | 25.2 | 100 | 36.705 | 0.117 | 0.105 | 36.274 | 0.000 | 37.398 | 0.001 |
| 151 | 100 | 1000000 | 1000 | 5 | 5000 | 5249 | 15.0 | 45.0 | 100 | 28.008 | 0.066 | 0.103 | 27.738 | 0.000 | 28.654 | 0.001 |
| 152 | 100 | 1000000 | 1000 | 5 | 5000 | 18671 | 15.0 | 45.0 | 100 | 29.334 | 0.069 | 0.105 | 29.053 | 0.000 | 29.975 | 0.001 |
| 153 | 100 | 1000000 | 1000 | 5 | 5000 | 147599 | 15.0 | 45.0 | 100 | 36.317 | 0.117 | 0.103 | 35.891 | 0.000 | 37.020 | 0.001 |
| 154 | 100 | 1000000 | 1000 | 50 | 50000 | 5249 | 5.0 | 15.0 | 100 | 28.598 | 0.066 | 0.105 | 28.324 | 0.000 | 29.242 | 0.001 |
| 155 | 100 | 1000000 | 1000 | 50 | 50000 | 18671 | 5.0 | 15.0 | 100 | 28.923 | 0.069 | 0.103 | 28.646 | 0.000 | 29.576 | 0.001 |
| 156 | 100 | 1000000 | 1000 | 50 | 50000 | 147599 | 5.0 | 15.0 | 100 | 36.276 | 0.117 | 0.105 | 35.847 | 0.000 | 36.961 | 0.001 |
| 157 | 100 | 1000000 | 1000 | 50 | 50000 | 5249 | 8.4 | 25.2 | 100 | 28.734 | 0.066 | 0.103 | 28.462 | 0.000 | 29.389 | 0.001 |
| 158 | 100 | 1000000 | 1000 | 50 | 50000 | 18671 | 8.4 | 25.2 | 100 | 29.293 | 0.069 | 0.105 | 29.012 | 0.000 | 29.929 | 0.001 |
| 159 | 100 | 1000000 | 1000 | 50 | 50000 | 147599 | 8.4 | 25.2 | 100 | 36.776 | 0.117 | 0.103 | 36.350 | 0.000 | 37.485 | 0.001 |
| 160 | 100 | 1000000 | 1000 | 50 | 50000 | 5249 | 15.0 | 45.0 | 100 | 28.617 | 0.066 | 0.105 | 28.343 | 0.000 | 29.248 | 0.001 |

*Continued on next page*

(continued from previous page)

| Run | $N$ | ECM | CELL | FOCAD/CELL | FOCAD | FNODE | $R_c$ | $R_s$ | Steps | $t_{init}$ | $t_{sim}$ | $t_{RTC}$ | $t_{initfn}$ | $t_{exit}$ | $t_{total}$ | $t_{step}$ |
|---|---|---|---|---|---|---|---|---|---|---|---|---|---|---|---|---|
| 161 | 100 | 1000000 | 1000 | 50 | 50000 | 18671 | 15.0 | 45.0 | 100 | 28.761 | 0.069 | 0.103 | 28.482 | 0.000 | 30.301 | 0.001 |
| 162 | 100 | 1000000 | 1000 | 50 | 50000 | 147599 | 15.0 | 45.0 | 100 | 36.216 | 0.117 | 0.126 | 35.764 | 0.000 | 36.906 | 0.001 |
| 163 | 100 | 1000000 | 10000 | 5 | 50000 | 5249 | 5.0 | 15.0 | 100 | 28.391 | 0.066 | 0.103 | 28.120 | 0.000 | 29.048 | 0.001 |
| 164 | 100 | 1000000 | 10000 | 5 | 50000 | 18671 | 5.0 | 15.0 | 100 | 29.323 | 0.069 | 0.105 | 29.041 | 0.000 | 29.958 | 0.001 |
| 165 | 100 | 1000000 | 10000 | 5 | 50000 | 147599 | 5.0 | 15.0 | 100 | 36.810 | 0.117 | 0.103 | 36.384 | 0.000 | 37.514 | 0.001 |
| 166 | 100 | 1000000 | 10000 | 5 | 50000 | 5249 | 8.4 | 25.2 | 100 | 30.746 | 0.066 | 0.106 | 30.471 | 0.000 | 31.381 | 0.001 |
| 167 | 100 | 1000000 | 10000 | 5 | 50000 | 18671 | 8.4 | 25.2 | 100 | 29.078 | 0.068 | 0.103 | 28.800 | 0.000 | 29.724 | 0.001 |
| 168 | 100 | 1000000 | 10000 | 5 | 50000 | 147599 | 8.4 | 25.2 | 100 | 36.564 | 0.117 | 0.105 | 36.135 | 0.000 | 37.272 | 0.001 |
| 169 | 100 | 1000000 | 10000 | 5 | 50000 | 5249 | 15.0 | 45.0 | 100 | 28.552 | 0.066 | 0.105 | 28.280 | 0.000 | 30.091 | 0.001 |
| 170 | 100 | 1000000 | 10000 | 5 | 50000 | 18671 | 15.0 | 45.0 | 100 | 28.945 | 0.069 | 0.105 | 28.663 | 0.000 | 30.486 | 0.001 |
| 171 | 100 | 1000000 | 10000 | 5 | 50000 | 147599 | 15.0 | 45.0 | 100 | 36.398 | 0.117 | 0.103 | 35.971 | 0.000 | 37.122 | 0.001 |
| 172 | 100 | 1000000 | 10000 | 50 | 500000 | 5249 | 5.0 | 15.0 | 100 | 28.483 | 0.066 | 0.106 | 28.207 | 0.000 | 29.122 | 0.001 |
| 173 | 100 | 1000000 | 10000 | 50 | 500000 | 18671 | 5.0 | 15.0 | 100 | 28.920 | 0.069 | 0.103 | 28.642 | 0.000 | 29.574 | 0.001 |
| 174 | 100 | 1000000 | 10000 | 50 | 500000 | 147599 | 5.0 | 15.0 | 100 | 36.182 | 0.117 | 0.105 | 35.751 | 0.000 | 36.904 | 0.001 |
| 175 | 100 | 1000000 | 10000 | 50 | 500000 | 5249 | 8.4 | 25.2 | 100 | 28.615 | 0.066 | 0.104 | 28.343 | 0.000 | 29.269 | 0.001 |
| 176 | 100 | 1000000 | 10000 | 50 | 500000 | 18671 | 8.4 | 25.2 | 100 | 29.316 | 0.069 | 0.106 | 29.034 | 0.000 | 29.977 | 0.001 |
| 177 | 100 | 1000000 | 10000 | 50 | 500000 | 147599 | 8.4 | 25.2 | 100 | 36.156 | 0.117 | 0.103 | 35.726 | 0.000 | 36.856 | 0.001 |
| 178 | 100 | 1000000 | 10000 | 50 | 500000 | 5249 | 15.0 | 45.0 | 100 | 28.536 | 0.066 | 0.106 | 28.258 | 0.000 | 29.172 | 0.001 |
| 179 | 100 | 1000000 | 10000 | 50 | 500000 | 18671 | 15.0 | 45.0 | 100 | 29.065 | 0.069 | 0.103 | 28.785 | 0.000 | 29.724 | 0.001 |
| 180 | 100 | 1000000 | 10000 | 50 | 500000 | 147599 | 15.0 | 45.0 | 100 | 36.638 | 0.117 | 0.106 | 36.206 | 0.000 | 37.319 | 0.001 |
| 181 | 100 | 1000000 | 100000 | 5 | 500000 | 5249 | 5.0 | 15.0 | 100 | 28.235 | 0.066 | 0.103 | 27.964 | 0.000 | 28.885 | 0.001 |
| 182 | 100 | 1000000 | 100000 | 5 | 500000 | 18671 | 5.0 | 15.0 | 100 | 29.211 | 0.069 | 0.105 | 28.930 | 0.000 | 29.854 | 0.001 |
| 183 | 100 | 1000000 | 100000 | 5 | 500000 | 147599 | 5.0 | 15.0 | 100 | 36.291 | 0.117 | 0.104 | 35.862 | 0.000 | 37.008 | 0.001 |
| 184 | 100 | 1000000 | 100000 | 5 | 500000 | 5249 | 8.4 | 25.2 | 100 | 28.380 | 0.066 | 0.106 | 28.103 | 0.000 | 29.012 | 0.001 |
| 185 | 100 | 1000000 | 100000 | 5 | 500000 | 18671 | 8.4 | 25.2 | 100 | 29.403 | 0.069 | 0.104 | 29.124 | 0.000 | 30.055 | 0.001 |
| 186 | 100 | 1000000 | 100000 | 5 | 500000 | 147599 | 8.4 | 25.2 | 100 | 36.708 | 0.117 | 0.108 | 36.269 | 0.000 | 38.302 | 0.001 |
| 187 | 100 | 1000000 | 100000 | 5 | 500000 | 5249 | 15.0 | 45.0 | 100 | 28.557 | 0.066 | 0.105 | 28.284 | 0.000 | 29.229 | 0.001 |
| 188 | 100 | 1000000 | 100000 | 5 | 500000 | 18671 | 15.0 | 45.0 | 100 | 28.749 | 0.069 | 0.106 | 28.463 | 0.000 | 29.387 | 0.001 |
| 189 | 100 | 1000000 | 100000 | 5 | 500000 | 147599 | 15.0 | 45.0 | 100 | 37.303 | 0.117 | 0.105 | 36.871 | 0.000 | 38.013 | 0.001 |
| 190 | 100 | 1000000 | 100000 | 50 | 5000000 | 5249 | 5.0 | 15.0 | 100 | 28.176 | 0.066 | 0.107 | 27.900 | 0.000 | 28.817 | 0.001 |
| 191 | 100 | 1000000 | 100000 | 50 | 5000000 | 18671 | 5.0 | 15.0 | 100 | 28.767 | 0.069 | 0.105 | 28.485 | 0.000 | 29.509 | 0.001 |
| 192 | 100 | 1000000 | 100000 | 50 | 5000000 | 147599 | 5.0 | 15.0 | 100 | 36.359 | 0.118 | 0.203 | 35.827 | 0.000 | 37.077 | 0.001 |
| 193 | 100 | 1000000 | 100000 | 50 | 5000000 | 5249 | 8.4 | 25.2 | 100 | 29.158 | 0.066 | 0.104 | 28.887 | 0.000 | 29.809 | 0.001 |
| 194 | 100 | 1000000 | 100000 | 50 | 5000000 | 18671 | 8.4 | 25.2 | 100 | 28.822 | 0.069 | 0.107 | 28.539 | 0.000 | 30.365 | 0.001 |
| 195 | 100 | 1000000 | 100000 | 50 | 5000000 | 147599 | 8.4 | 25.2 | 100 | 36.663 | 0.118 | 0.104 | 36.232 | 0.000 | 37.370 | 0.001 |
| 196 | 100 | 1000000 | 100000 | 50 | 5000000 | 5249 | 15.0 | 45.0 | 100 | 28.027 | 0.066 | 0.107 | 27.750 | 0.000 | 28.662 | 0.001 |
| 197 | 100 | 1000000 | 100000 | 50 | 5000000 | 18671 | 15.0 | 45.0 | 100 | 29.668 | 0.069 | 0.104 | 29.387 | 0.000 | 30.331 | 0.001 |
| 198 | 100 | 1000000 | 100000 | 50 | 5000000 | 147599 | 15.0 | 45.0 | 100 | 36.441 | 0.117 | 0.108 | 35.998 | 0.000 | 37.131 | 0.001 |
| 199 | 100 | 1000000 | 1000000 | 5 | 5000000 | 5249 | 5.0 | 15.0 | 100 | 28.548 | 0.066 | 0.104 | 28.275 | 0.000 | 29.196 | 0.001 |
| 200 | 100 | 1000000 | 1000000 | 5 | 5000000 | 18671 | 5.0 | 15.0 | 100 | 29.391 | 0.069 | 0.106 | 29.107 | 0.000 | 30.036 | 0.001 |
| 201 | 100 | 1000000 | 1000000 | 5 | 5000000 | 147599 | 5.0 | 15.0 | 100 | 36.555 | 0.117 | 0.105 | 36.122 | 0.000 | 38.145 | 0.001 |
| 202 | 100 | 1000000 | 1000000 | 5 | 5000000 | 5249 | 8.4 | 25.2 | 100 | 28.718 | 0.066 | 0.361 | 28.186 | 0.000 | 29.356 | 0.001 |
| 203 | 100 | 1000000 | 1000000 | 5 | 5000000 | 18671 | 8.4 | 25.2 | 100 | 28.767 | 0.074 | 0.104 | 28.488 | 0.000 | 29.432 | 0.001 |
| 204 | 100 | 1000000 | 1000000 | 5 | 5000000 | 147599 | 8.4 | 25.2 | 100 | 37.550 | 0.117 | 0.107 | 37.114 | 0.000 | 38.563 | 0.001 |
| 205 | 100 | 1000000 | 1000000 | 5 | 5000000 | 5249 | 15.0 | 45.0 | 100 | 28.772 | 0.066 | 0.104 | 28.500 | 0.000 | 29.428 | 0.001 |
| 206 | 100 | 1000000 | 1000000 | 5 | 5000000 | 18671 | 15.0 | 45.0 | 100 | 29.173 | 0.069 | 0.106 | 28.889 | 0.000 | 29.810 | 0.001 |
| 207 | 100 | 1000000 | 1000000 | 5 | 5000000 | 147599 | 15.0 | 45.0 | 100 | 36.811 | 0.117 | 0.104 | 36.378 | 0.000 | 37.524 | 0.001 |
| 208 | 100 | 1000000 | 1000000 | 50 | 50000000 | 5249 | 5.0 | 15.0 | 100 | 28.495 | 0.066 | 0.107 | 28.217 | 0.000 | 29.132 | 0.001 |
| 209 | 100 | 1000000 | 1000000 | 50 | 50000000 | 18671 | 5.0 | 15.0 | 100 | 29.191 | 0.069 | 0.104 | 28.912 | 0.000 | 29.857 | 0.001 |
| 210 | 100 | 1000000 | 1000000 | 50 | 50000000 | 147599 | 5.0 | 15.0 | 100 | 37.534 | 0.117 | 0.107 | 37.098 | 0.000 | 38.233 | 0.001 |
| 211 | 100 | 1000000 | 1000000 | 50 | 50000000 | 5249 | 8.4 | 25.2 | 100 | 28.333 | 0.066 | 0.104 | 28.061 | 0.000 | 28.990 | 0.001 |
| 212 | 100 | 1000000 | 1000000 | 50 | 50000000 | 18671 | 8.4 | 25.2 | 100 | 28.362 | 0.069 | 0.107 | 28.078 | 0.000 | 29.004 | 0.001 |
| 213 | 100 | 1000000 | 1000000 | 50 | 50000000 | 147599 | 8.4 | 25.2 | 100 | 36.980 | 0.117 | 0.105 | 36.547 | 0.000 | 37.704 | 0.001 |
| 214 | 100 | 1000000 | 1000000 | 50 | 50000000 | 5249 | 15.0 | 45.0 | 100 | 28.405 | 0.066 | 0.107 | 28.127 | 0.000 | 29.053 | 0.001 |

Continued on next page

*(continued from previous page)*

| Run | $N$ | ECM | CELL | FOCAD/CELL | FOCAD | FNODE | $R_c$ | $R_s$ | Steps | $t_{\text{init}}$ | $t_{\text{sim}}$ | $t_{\text{RTC}}$ | $t_{\text{initfn}}$ | $t_{\text{exit}}$ | $t_{\text{total}}$ | $t_{\text{step}}$ |
|---|---|---|---|---|---|---|---|---|---|---|---|---|---|---|---|---|
| 215 | 100 | 1000000 | 1000000 | 50 | 50000000 | 18671 | 15.0 | 45.0 | 100 | 28.983 | 0.069 | 0.104 | 28.703 | 0.000 | 29.643 | 0.001 |
| 216 | 100 | 1000000 | 1000000 | 50 | 50000000 | 147599 | 15.0 | 45.0 | 100 | 36.932 | 0.117 | 0.106 | 36.496 | 0.000 | 37.626 | 0.001 |

### F. Agent API Summary

This section provides a comprehensive overview of all pre-defined variables and functions associated with each agent type, detailing their purpose, structure, and role within the model to facilitate understanding, customization, and extension of the framework.

#### CELL agent

| CELL | |
|---|---|
| **Variable** | **Description** |
| id | int, unique cell id |
| x | float, cell-center position [um] |
| y | float |
| z | float |
| vx | float, cell velocity [um/s] |
| vy | float |
| vz | float |
| trajectory_length | float, cumulative path length since birth/latest division [um] |
| trajectory_time | float, elapsed tracked lifetime since birth/latest division [s] |
| birth_x | float, reference position for effective speed [um] |
| birth_y | float |
| birth_z | float |
| orx | float, cell polarity/orientation unit vector |
| ory | float |
| orz | float |
| cell_type | int, to represent different phenotypes (e.g. different cell lines). The specific meaning of the values assigned to this variable is up to the user and is not defined by the model. |
| k_elast | float, cell stiffness [nN/um] per-type; set during init (unused in the current implementation) |
| d_dumping | float, cell damping coefficient [nN*s/um] per-type; set during init (unused in the current implementation) |
| alignment | float, alignment score with the local fibre field [-] (unused in the current implementation) |
| k_consumption | float[N_SPECIES], per-species consumption rate constants |
| k_production | float[N_SPECIES], per-species production rate constants |
| k_reaction | float[N_SPECIES], per-species reaction rate constants |
| C_sp | float[N_SPECIES], per-species cell-associated concentration state |
| M_sp | float[N_SPECIES], per-species cell-associated mass state |
| speed_ref | float, per-type; set during init |
| radius | float, per-type; set during init |
| nucleus_radius | float, per-type; set during init |
| cc_dvx | float, [um/s] velocity contribution from cell_cell_interaction |
| cc_dvy | float |
| cc_dvz | float |
| cf_dvx | float, [um/s] velocity contribution from cell_fnode_repulsion |
| cf_dvy | float |
| cf_dvz | float |
| cycle_phase | int, [1:G1] [2:S] [3:G2] [4:M] |
| clock | float, internal clock of the cell to switch phases |
| completed_cycles | int, number of completed cell cycles |
| max_global_cell_id | int, cached global max CELL id (to atomically track newly created cells) |
| damage | float, accumulated damage score in [0,1], where 1 is lethal threshold |

| CELL (continued) | |
|---|---|
| **Variable** | **Description** |
| dead | int, 0: alive, 1: dead (dead cells can be kept as debris or removed using the DEAD_CELLS_DISAPPEAR flag) |
| dead_by | int, -1:none, 0:hypoxia, 1:starvation, 2:mechanical, 3:cumulative_damage |
| mother_id | int, id of the parent cell if this cell is a daughter |
| daughter_id | int, id of the daughter created in the latest division |
| just_divided | int, 1 during the step immediately after division |
| marked_for_removal | int, 1 if the cell should be removed |
| fnode_birth_cooldown | float, refractory time before creating another FNODE [s] |
| focad_birth_cooldown | float, refractory time before creating another FOCAD [s] |
| x_i | float[N_ANCHOR_POINTS], focal-adhesion anchor point positions on the cell nucleus surface. Unused if INCLUDE_FOCAL_ADHESIONS is False |
| y_i | float[N_ANCHOR_POINTS] |
| z_i | float[N_ANCHOR_POINTS] |
| u_ref_x_i | float[N_ANCHOR_POINTS], unit direction vector from the cell center to the anchor point in the reference configuration (used for elastic force calculation). Unused if INCLUDE_FOCAL_ADHESIONS is False |
| u_ref_y_i | float[N_ANCHOR_POINTS] |
| u_ref_z_i | float[N_ANCHOR_POINTS] |
| eps_xx | float, strain tensor |
| eps_yy | float |
| eps_zz | float |
| eps_xy | float |
| eps_xz | float |
| eps_yz | float |
| sig_xx | float, stress tensor [kPa] |
| sig_yy | float |
| sig_zz | float |
| sig_xy | float |
| sig_xz | float |
| sig_yz | float |
| sig_eig_1 | float, principal stresses (eigen values) [kPa] |
| sig_eig_2 | float |
| sig_eig_3 | float |
| sig_eigvec1_x | float, first principal-stress direction |
| sig_eigvec1_y | float |
| sig_eigvec1_z | float |
| sig_eigvec2_x | float, second principal-stress direction |
| sig_eigvec2_y | float |
| sig_eigvec2_z | float |
| sig_eigvec3_x | float, third principal-stress direction |
| sig_eigvec3_y | float |
| sig_eigvec3_z | float |
| eps_eig_1 | float, principal strains (eigen values)[-] |
| eps_eig_2 | float |
| eps_eig_3 | float |
| eps_eigvec1_x | float, first principal-strain direction |
| eps_eigvec1_y | float |
| eps_eigvec1_z | float |
| eps_eigvec2_x | float, second principal-strain direction |
| eps_eigvec2_y | float |
| eps_eigvec2_z | float |

| CELL (continued) | |
|---|---|
| **Variable** | **Description** |
| eps_eigvec3_x | float, third principal-strain direction |
| eps_eigvec3_y | float |
| eps_eigvec3_z | float |
| chemotaxis_sensitivity | float[N_SPECIES], per-species chemotactic sensitivity weights |

##### Functions

| CELL | | |
|---|---|---|
| **Function** | **Input** | **Output** |
| cell_spatial_location_data | None | cell_spatial_location_message |
| cell_cell_interaction | cell_spatial_location_message | None |
| cell_fnode_repulsion | fnode_spatial_location_message | None |
| cell_ecm_interaction_metabolism | ecm_grid_location_message | None |
| cell_move | None | None |
| cell_bucket_location_data | None | cell_bucket_location_message |
| cell_MaxID_update | None | None |

#### FOCAD agent

| FOCAD | |
|---|---|
| **Variable** | **Description** |
| id | int, unique focal-adhesion id |
| cell_id | int, id of the owner cell |
| cell_type | int, cell type of the owner cell (for per-type property lookups) |
| fnode_id | int, id of the interacting fibre node if attached (-1 if not) |
| x | float, focal-adhesion position [um] |
| y | float |
| z | float |
| vx | float, focal-adhesion velocity [um/s] |
| vy | float |
| vz | float |
| fx | float, net force on the focal adhesion [nN] |
| fy | float |
| fz | float |
| anchor_id | int, index of the associated cell anchor point |
| x_i | float, anchor-point position on the cell surface [um] |
| y_i | float |
| z_i | float |
| x_c | float, owner-cell center position [um] |
| y_c | float |
| z_c | float |
| orx | float, owner-cell orientation unit vector |
| ory | float |
| orz | float |
| rest_length_0 | float, rest length at adhesion birth [um] |
| rest_length | float, current effective rest length [um] |

| FOCAD (continued) | |
|---|---|
| **Variable** | **Description** |
| k_fa | float, focal-adhesion stiffness [nN/um] |
| f_max | float, maximum sustainable traction force [nN] |
| attached | int, 1 if attached to a fibre node |
| active | uint8, 1 if the adhesion is active in the simulation |
| v_c | float, rest-length shortening speed [um/s] |
| fa_state | uint8, adhesion state code: 1 nascent, 2 mature, 3 disassembling |
| age | float, adhesion lifetime [s] |
| detached_age | float, time spent detached [s] |
| k_on | float, attachment rate [1/s] |
| k_off_0 | float, baseline detachment rate [1/s] |
| f_c | float, force scale in the slip-bond law [nN] |
| k_reinf | float, force-dependent reinforcement rate [1/s] |
| f_mag | float, \|F_FA\| traction magnitude [nN] at current step |
| is_front | int, 1 if adhesion is classified in the cell front hemisphere, else 0 |
| is_rear | int, 1 if adhesion is classified in the cell rear hemisphere, else 0 |
| attached_front | int, 1 if attached and in front |
| attached_rear | int, 1 if attached and in rear |
| frontness_front | float, frontness score used for front-biased kinetics (front branch). Polarity score p in [-1,1] from orientation vs anchor direction (cell center towards anchor) |
| frontness_rear | float, rearness score used for rear-biased kinetics |
| k_on_eff_front | float, effective attachment rate used for front-side update [1/s] |
| k_on_eff_rear | float, effective attachment rate used for rear-side update [1/s] |
| k_off_0_eff_front | float, effective baseline detachment rate at front [1/s] |
| k_off_0_eff_rear | float, effective baseline detachment rate at rear [1/s] |
| linc_prev_total_length | float, previous-step LINC internal length state for Kelvin-Voigt-in-series solve [um] |

##### Functions

| FOCAD | | |
|---|---|---|
| **Function** | **Input** | **Output** |
| focad_bucket_location_data | None | focad_bucket_location_message |
| focad_spatial_location_data | None | focad_spatial_location_message |
| focad_anchor_update | cell_bucket_location_message | None |
| focad_move | fnode_bucket_location_message | None |

#### FNODE agent

| FNODE | |
|---|---|
| **Variable** | **Description** |
| id | int, unique fibre-node id |
| x | float, fibre-node position [um] |
| y | float |
| z | float |
| vx | float, fibre-node velocity [um/s] |
| vy | float |

| FNODE (continued) | |
|---|---|
| **Variable** | **Description** |
| vz | float |
| fx | float, net force on the fibre node [nN] |
| fy | float |
| fz | float |
| k_elast | float, effective segment stiffness [nN/um] |
| d_dumping | float, effective segment damping [nN*s/um] |
| equilibrium_distance | float[MAX_CONNECTIVITY], each segment can have a different equilibrium distance depending on the rest length assigned during network generation |
| boundary_fx | float, boundary_f[A]: normal force coming from boundary [A] when elastic boundaries option is selected. |
| boundary_fy | float |
| boundary_fz | float |
| f_bx_pos | float, f_b[A]_[B]: normal force transmitted to the boundary [A]_[B] when agent is clamped |
| f_bx_neg | float |
| f_by_pos | float |
| f_by_neg | float |
| f_bz_pos | float |
| f_bz_neg | float |
| f_bx_pos_y | float, f_b[A]_[B]_[C]: shear force transmitted to the boundary [A]_[B] in the direction [C] when agent is clamped |
| f_bx_pos_z | float |
| f_bx_neg_y | float |
| f_bx_neg_z | float |
| f_by_pos_x | float |
| f_by_pos_z | float |
| f_by_neg_x | float |
| f_by_neg_z | float |
| f_bz_pos_x | float |
| f_bz_pos_y | float |
| f_bz_neg_x | float |
| f_bz_neg_y | float |
| f_extension | float, tensile load carried by connected segments [nN] |
| f_compression | float, compressive load carried by connected segments [nN] |
| elastic_energy | float, stored elastic energy [nN*um] |
| connectivity_count | uint8, number of linked neighbour nodes |
| degradation | float, accumulated degradation state [-] |
| reinforcement | float, accumulated reinforcement state [-] |
| secreted | int, 1 if this node was newly secreted by a cell |
| marked_for_removal | int, 1 if the node should be deleted |
| closest_fnode_id | int, id of the nearest neighbouring FNODE |
| second_closest_fnode_id | int, id of the second-nearest neighbouring FNODE |
| linked_nodes | float[MAX_CONNECTIVITY], ids of connected neighbour FNODEs |
| clamped_bx_pos | uint8, boundary clamp flags for each face (1 = clamped) |
| clamped_bx_neg | uint8 |
| clamped_by_pos | uint8 |
| clamped_by_neg | uint8 |
| clamped_bz_pos | uint8 |
| clamped_bz_neg | uint8 |

| FNODE (continued) | |
|---|---|
| **Variable** | **Description** |
| unstable_move | uint8, flag used to indicate when unrealistically large movements occur within a single simulation step, enabling warnings to the user or termination of the execution |

##### Functions

| FNODE | | |
|---|---|---|
| **Function** | **Input** | **Output** |
| fnode_spatial_location_data | None | fnode_spatial_location_message |
| fnode_bucket_location_data | None | fnode_bucket_location_message |
| fnode_boundary_interaction | None | None |
| fnode_update_links | fnode_bucket_location_message | None |
| fnode_fnode_spatial_interaction | fnode_spatial_location_message | None |
| fnode_fnode_bucket_interaction | fnode_bucket_location_message | None |
| fnode_remodel | cell_spatial_location_message | None |
| fnode_move | None | None |
| fnode_focad_interaction | focad_spatial_location_message | None |
| fnode_cell_repulsion | cell_spatial_location_message | None |

#### ECM agent

| ECM | |
|---|---|
| **Variable** | **Description** |
| id | int, unique ECM-agent id |
| x | float, ECM grid-point spatial position [um] |
| y | float |
| z | float |
| grid_lin_id | int, linear index in the 3D grid that maps to i,j,k positions |
| grid_i | uint8, grid index, (i,j,k) maps to (x,y,z) |
| grid_j | uint8 |
| grid_k | uint8 |
| D_sp | float[N_SPECIES], diffusion coefficient of each species at the agent location (used for heterogeneous diffusion) |
| C_sp | float[N_SPECIES], species concentrations at this ECM node |
| C_sp_sat | float[N_SPECIES], saturation concentrations for each species |
| k_elast | float, ECM spring stiffness [nN/um] (used only for smooth grid adapation if boundaries move) |
| d_dumping | float, ECM damping coefficient [nN*s/um] (used only for smooth grid adapation if boundaries move) |
| vx | float, ECM grid-point velocity [um/s] (used only for smooth grid adapation if boundaries move) |
| vy | float |
| vz | float |
| fx | float, net force on the ECM grid-point [nN] (used only for smooth grid adapation if boundaries move) |
| fy | float |

| ECM (continued) | |
|---|---|
| **Variable** | **Description** |
| fz | float |
| clamped_bx_pos | uint8, boundary clamp flags for each face (1 = clamped) (used only for smooth grid adapation if boundaries move) |
| clamped_bx_neg | uint8 |
| clamped_by_pos | uint8 |
| clamped_by_neg | uint8 |
| clamped_bz_pos | uint8 |
| clamped_bz_neg | uint8 |

**Functions**

| ECM | | |
|---|---|---|
| **Function** | **Input** | **Output** |
| ecm_grid_location_data | None | ecm_grid_location_message |
| ecm_ecm_interaction | ecm_grid_location_message | None |
| ecm_boundary_concentration_conditions | None | None |
| ecm_Csp_update | None | None |
| ecm_Dsp_update | fnode_spatial_location_message | None |
| ecm_move | None | None |

##### BCORNER agent

This agent does not participate in the simulation dynamics and serves solely as a visual aid to represent the position and extent of the domain boundaries.

| BCORNER | |
|---|---|
| **Variable** | **Description** |
| id | int, unique boundary-corner id |
| x | float, boundary corner position [um] |
| y | float |
| z | float |

**Functions**

| BCORNER | | |
|---|---|---|
| **Function** | **Input** | **Output** |
| bcorner_output_location_data | None | bcorner_location_message |
| bcorner_move | None | None |

#### G. Agents' C++ Functions Summary

Table 18: Function summary table

■ ECM ■ CELL ■ FOCAD ■ FNODES ■ BOUNDARIES

| Function name | Input type | Output type | Description |
|---|---|---|---|
| bcorner_output_location_data | None | Spatial | Publish BCORNER identifiers and coordinates to spatial messages. |
| cell_spatial_location_data | None | Spatial | Broadcast CELL kinematics and metabolic parameters over a spatial message list. |
| ecm_grid_location_data | None | Array | Publish ECM voxel-centered state into the Array3D message for neighborhood reads. |
| cell_bucket_location_data | None | Bucket | Export CELL state required by bucket-based readers (e.g., focal adhesion updates). |
| focad_anchor_update | Bucket | None | Re-anchor each FOCAD agent to a CELL nucleus anchor point read from bucket messages keyed by cell_id. |
| fnode_spatial_location_data | None | Spatial | Broadcast FNODE position for spatial proximity queries. |
| fnode_bucket_location_data | None | Bucket | Export FNODE state and connectivity arrays into a bucket message keyed by node id. |
| ecm_boundary_ concentration_conditions | None | None | Apply boundary concentration conditions to ECM agents located near domain faces. |
| fnode_boundary_interaction | None | None | Compute boundary reaction forces on FNODE agents near domain boundaries, including optional elastic and damping contributions per face. |
| cell_fnode_remodel | Spatial | None | Probabilistically create a single FNODE around a CELL and request reciprocal parent-link update through environment macros. |
| fnode_remodel | Spatial | None | Update FNODE degradation/reinforcement state from nearby CELLs and register removal requests when net degradation reaches 1. |
| fnode_apply_remodel_ updates | Spatial | None | Apply remodeling topology updates and optionally remove terminally degraded nodes. |
| fnode_update_links | Bucket | None | Update FNODE link list using bucket messages keyed by linked node id. If a linked node has no bucket message (e.g., removed), clear that link. |
| cell_ecm_interaction_ metabolism | Array | None | Couple each CELL to its nearest ECM voxel for species exchange and run intracellular metabolic reactions with mass-consistent updates. |
| cell_MaxID_update | None | None | Synchronize the agent's max_global_cell_id variable with the environment macro property. |
| cell_cycle | None | None | Agent function for cell cycle progression, division, and death. Handles cell phase transitions, damage accumulation, and division logic. |
| cell_bucket_location_data | None | Bucket | Export CELL state required by bucket-based readers (e.g., focal adhesion updates). |
| focad_post_cycle_update | Bucket | None | Update FOCAD anchor association after cell division, switching anchor points between parent and daughter cells. Ensures spatial and orientation variables are updated for correct cell association. |

Continued on next page

■ ECM ■ CELL ■ FOCAD ■ FNODES ■ BOUNDARIES

| Function name | Input type | Output type | Description |
|---|---|---|---|
| ecm_Csp_update | None | None | Refresh each ECM voxel concentration array from the global macro property buffer. |
| ecm_Dsp_update | Spatial | None | Compute local FNODE crowding around each ECM voxel and downscale diffusion coefficients to represent heterogeneous transport in dense regions. |
| ecm_ecm_interaction | Array | None | Execute ECM voxel-to-voxel mechanical coupling and multi-species diffusion on the same neighborhood pass. |
| ecm_boundary_ concentration_conditions | None | None | Apply boundary concentration conditions to ECM agents located near domain faces. |
| fnode_fnode_spatial_ interaction | Spatial | None | Apply short-range repulsion between nearby FNODE agents to prevent overlap. |
| fnode_fnode_bucket_ interaction | Bucket | None | Compute spring-damper forces along explicit FNODE connectivity links and accumulate network mechanical metrics (extension/compression/elastic energy). |
| focad_fnode_interaction | Spatial | None | Manage FOCAD-FNODE attachment dynamics and compute traction forces stored on FOCAD for subsequent FNODE-side force transfer. |
| focad_spatial_location_data | None | Spatial | Broadcast active adhesion position/force state for local spatial interaction queries. |
| focad_bucket_location_data | None | Bucket | Publish full FOCAD state for bucket-keyed readers (mainly CELL/FOCAD coupling steps). |
| fnode_focad_interaction | Spatial | None | Transfer precomputed FOCAD traction forces onto the corresponding FNODE. |
| cell_focad_update | Bucket | None | Reads all focal adhesion (FOCAD) messages in a bucket keyed by this cell id. |
| cell_cell_interaction | Spatial | None | Compute short-range CELL-CELL mechanics with strong contact repulsion and weak finite-range adhesion shell (soft cohesion) to promote aggregate compactness while allowing escape under other motility cues. |
| cell_fnode_repulsion | Spatial | None | Prevent CELL centers from approaching FNODE points closer than an exclusion distance by adding a short-range repulsive velocity component. |
| fnode_cell_repulsion | Spatial | None | Prevent FNODE points from being pushed into CELL centers by adding a short-range repulsive force (Newton-3 counterpart of cell_fnode_repulsion). |
| cell_move | None | None | Update CELL velocity/orientation-driven migration by combining Brownian, chemotactic, and durotactic components, then advance position. |
| fnode_move | None | None | Update FNODE positions/velocities under internal, boundary, and transmitted forces while enforcing clamp and sliding boundary behavior. |
| focad_move | Bucket | None | Update focal adhesion positions by either following attached FNODEs or executing bounded exploratory motion when detached/inactive. |

Continued on next page

■ ECM ■ CELL ■ FOCAD ■ FNODES ■ BOUNDARIES

| Function name | Input type | Output type | Description |
|---|---|---|---|
| bcorner_move | None | None | Synchronize each BCORNER agent position with the current domain boundary coordinates. |
| ecm_move | None | None | Advance ECM agent motion from accumulated forces, then enforce boundary clamping/sliding rules and boundary-driven kinematics. |

#### H. Video Captions

**Video Supp.Video.1.** Showcase of 100 cells migrating through the extracellular matrix, illustrating how adhesion formation, force transmission, and detachment contribute to the resulting trajectories and long-range force transmission. CELL agents are represented as semi-transparent spheres, including nuclei (coloured by deformation) and nuclei anchor points (white). FOCAD agents are shown as straight rods coloured by adhesion state (green means attached). The surrounding matrix is visualised through the positions of FNODE agents (small light blue spheres on the right panel), joined by lines (fibres) whose colour reflect their local elastic energy. This video highlights the dynamic interplay between cellular motility and matrix mechanics in a complex microenvironment.

Link to video 1: `https://drive.google.com/file/d/1gWTfK3YCguf6KjD9w4iat7Ccx3IyUnZS/view?usp=sharing`

**Video Supp.Video.2.** This video demonstrates the model configurator workflow, including agent definition and visual interaction linking. More information can be found in its own repository at `https://github.com/cborau/flamegpu2_uiconfig`.

Link to video 2: `https://drive.google.com/file/d/1fJTrgfqePtavwgBk20KPtsRoFw5TPJoG/view?usp=sharing`

**Video Supp.Video.3.** This video illustrates the integration of chemical and mechanical interactions in the model, where cell motility and matrix remodelling are influenced by local biochemical cues, while cells also actively modify their microenvironment through secretion and consumption of diffusible factors. In this example, 100 cells are shown. The cells consume species 0 (e.g. nutrients present in the environment) and release species 1, generating heterogeneous concentration patterns in the surrounding domain. The blobs are coloured and scaled according to the local concentration values of species 1.

Link to video 3: `https://drive.google.com/file/d/13crL01TN6DYW49jp_UcKSprt5S1MRUSo/view?usp=sharing`

**Video Supp.Video.4.** Matrix degradation by a migrating cell. Surrounding fibre nodes (FNODEs) within the interaction range are represented as blobs whose size and colour reflect their local degradation state. As degradation progresses and reaches a value of 1, the corresponding FNODE is removed from the simulation, thereby modelling local matrix breakdown, relaxing mechanical constraints and enabling cell invasion through the matrix.

Link to video 4: `https://drive.google.com/file/d/1WIe3hLTs1z95FOxNvR_n9mjT8uojnrge/view?usp=sharing`

**Video Supp.Video.5.** Matrix reinforcement driven by a migrating cell. Surrounding fibre nodes (FNODEs) within the interaction range are shown as blobs whose size and colour indicate the local reinforcement level. Reinforcement promotes the generation of new FNODEs and connected fibres, enabling progressive matrix deposition and remodelling around the cell. This can lead to local stiffening and densification of the matrix, which in turn can influence cell motility, force transmission and diffusion of species.

Link to video 5: `https://drive.google.com/file/d/1wPA-thez7thd5JEsg343QJJkIa3xvKsl/view?usp=sharing`

**Video Supp.Video.6.** Organoid growth driven by the proliferation of three cell types that progressively assemble into a compact three-dimensional structure. Cells are coloured by cell type in the left panel and by cell-cycle phase in the right panel, highlighting both the emerging tissue composition and the heterogeneous proliferative dynamics during organoid formation.

Link to video 6: https://drive.google.com/file/d/17AsDhWSK0Fx9J_6w3zD_WH_-enS1vSah/view?usp=sharing
